# EWS/FLI mediated reprogramming of 3D chromatin promotes an altered transcriptional state in Ewing sarcoma

**DOI:** 10.1101/2021.09.30.462658

**Authors:** Iftekhar A. Showpnil, Julia Selich-Anderson, Cenny Taslim, Megann A. Boone, Jesse C. Crow, Emily R. Theisen, Stephen L. Lessnick

## Abstract

Ewing sarcoma is a prototypical fusion transcription factor-associated pediatric cancer that expresses EWS/FLI or highly related fusions. EWS/FLI dysregulates transcription to induce and maintain sarcomagenesis, but the mechanisms utilized are not fully understood. We therefore sought to define the global effects of EWS/FLI on chromatin conformation and transcription in Ewing sarcoma. We found that EWS/FLI (and EWS/ERG) genomic localization is largely conserved across multiple patient-derived Ewing sarcoma cell lines. EWS/FLI binding is primarily associated with compartment activation, establishment of topologically-associated domain (TAD) boundaries, enhancer-promoter looping that involve both intra- and inter-TAD interactions, and gene activation. Importantly, local chromatin features provide the basis for transcriptional heterogeneity in regulation of direct EWS/FLI target genes across different Ewing sarcoma cell lines. These data demonstrate a key role of EWS/FLI in mediating genome-wide changes in chromatin configuration and support the notion that fusion transcription factors serve as master regulators through three-dimensional reprogramming of chromatin.

## INTRODUCTION

Ewing sarcoma is a highly aggressive pediatric bone cancer characterized by FET/ETS fusion oncoprotein expression (Delattre et al., 1992; Sankar and Lessnick, 2011). The FET family of RNA binding proteins, FUS, EWS, and TAF15 contain largely conserved low complexity domains (LCD) at the amino terminus, while the ETS (E26 transformation specific) family of transcription factors contain a highly conserved winged helix-loop-helix DNA-binding domain (DBD) (Schwartz et al., 2015; Seth and Watson, 2005). EWS/FLI, the most common FET/ETS fusion found in Ewing sarcoma (in ∼85% of cases), is generated by the (11;22)(q24;q12) chromosomal translocation that fuses the amino terminus of the conserved FET LCD of EWS to the carboxyl terminus of FLI containing the ETS DBD (Delattre et al., 1992; Turc-Carel et al., 1988; Turc-Carel et al., 1984).

The FLI ETS DBD of EWS/FLI is crucial for DNA binding and oncogenesis in Ewing sarcoma (May et al., 1993a; Smith et al., 2006). Genome-wide localization studies have identified GGAA-repeat elements (GGAA-microsatellites/GGAA-μsats) as well as consensus ETS sites (ACC<u>GGAA</u>GTG) as EWS/FLI response elements in Ewing sarcoma (Gangwal et al., 2008; May et al., 1993b). GGAA-μsat binding by EWS/FLI is critical for its oncogenic function (Boulay et al., 2018; Gangwal et al., 2008; Johnson et al., 2017).

The conserved FET LCD of EWS in EWS/FLI can undergo liquid-liquid phase transition and multimerization *in vitro* (Kwon et al., 2013). This multimerization property of the EWS LCD is required for EWS/FLI binding, chromatin accessibility, enhancer establishment, and formation of transcriptional hubs at GGAA-μsats *in vivo* (Boulay et al., 2017; Boulay et al., 2018; Chong et al., 2018; Johnson et al., 2017; Riggi et al., 2014). The ability to bind, multimerize, and form transcriptional hubs at GGAA-μsats, suggests a model in which GGAA-μsats bound by EWS/FLI interact with other EWS/FLI bound DNA to promote chromatin looping (Nacev et al., 2020), altered 3D chromatin conformation, and dysregulated gene expression in Ewing sarcoma.

We evaluated this model using an unbiased whole-genome approach, *in situ* Hi-C, to define the global changes in chromatin structure mediated by EWS/FLI in Ewing sarcoma (Rao et al., 2014). Integration of Hi-C with CUT&Tag (cleavage under target and tagmentation), RNA-sequencing and 4C (circular chromatin conformation capture) defined how EWS/FLI binding to chromatin alters the 3D chromatin landscape to facilitate an oncogenic transcriptional state in Ewing sarcoma.

## RESULTS

### EWS/FLI reprograms the global interaction profile in Ewing sarcoma

To understand how EWS/FLI affects 3D chromatin conformation in Ewing sarcoma, we depleted EWS/FLI expression from patient derived Ewing sarcoma cells and compared chromatin features between EWS/FLI expressing and depleted states. Endogenous EWS/FLI (EF-Endo) was knocked-down from A673 cells using RNAi (EF-KD), and subsequently rescued with ectopic expression of a 3X-FLAG tagged EWS/FLI cDNA construct (EF-Rescue), using a well-validated knock-down/rescue system (Smith et al., 2006; Theisen et al., 2019). The EF-Rescue experiment was used to confirm that the changes observed between EF-Endo and EF-KD were not off-target effects of RNAi. EWS/FLI knock-down and rescue of expression were verified using quantitative reverse transcription polymerase chain reaction (qRT-PCR) and western blot analysis (Figures S1A-C). *In situ* Hi-C experiments were performed, in duplicate, in EF-Endo, EF-KD and EF-Rescue A673 cells to obtain 201 – 311 million unique paired-end (PE) reads per replicate. In this study, chromatin features in EWS/FLI expressing (EF-WT) states, i.e. EF-Endo and EF-Rescue were compared to chromatin features in the EWS/FLI depleted (EF-KD) state.

We first generated genome-wide contact maps using the Hi-C data for EF-Endo, EF-KD and EF-Rescue (Figure S1D). Off diagonal inter-chromosomal signals represent chromosomal rearrangements in A673 cells, many of which were reported previously (e.g. t(1;9), t(1;13), t(3;16), t(5;8), t(9;13), t(11;13) and t(11;22) (Martinez-Ramirez et al., 2003). To determine whether EWS/FLI promotes global changes in the chromatin interaction profile, we performed multidimensional scaling (MDS) analysis of the top 1000 interactions at 1 megabase [mb] resolution in EF-Endo, EF-KD and EF-Rescue (Figure 1A). EF-Endo clustered with EF-Rescue, and were separated from the EF-KD replicates, demonstrating that the changes in interaction profiles observed with EWS/FLI depletion are largely reversed by re-expression of the “rescue” EWS/FLI.

**Figure 1.**
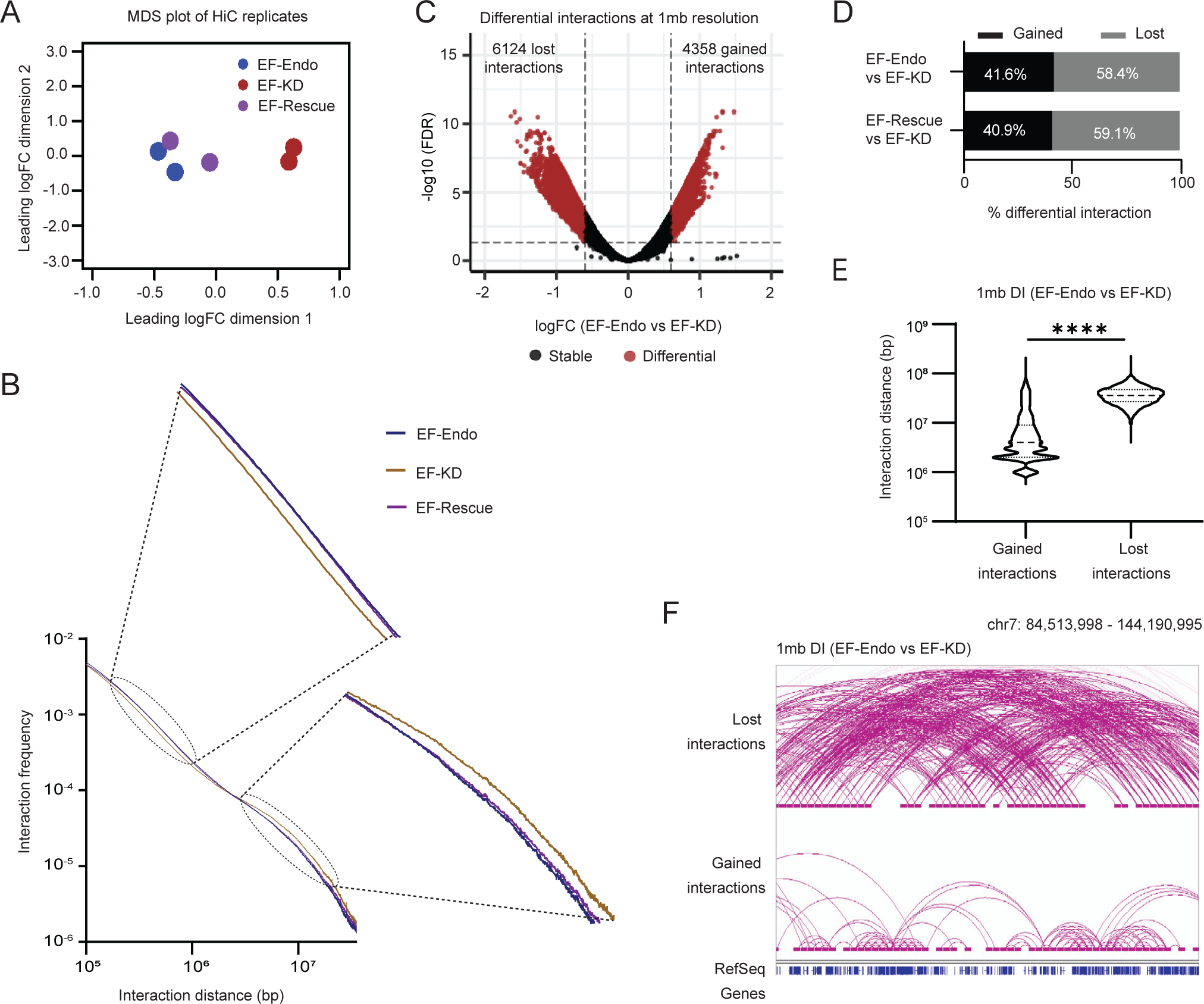
**EWS/FLI reprograms the global interaction profile in Ewing sarcoma** (A) Multidimensional scaling (MDS) plot of top 1000 interactions (1mb resolution) in each Hi-C replicate. (B) Genome-wide interaction frequency over interaction distance (bp) curves for EF-Endo, EF-KD and EF-Rescue. (C) Volcano plot showing differential interactions (DI) detected at 1mb resolution for EF-Endo vs. EF-KD A673 cells using diffHiC R package. DI, FDR <0.05 and |logFC| > 0.6. (D) Percent DIs gained (FDR <0.05 & logFC > 0.6) or lost (FDR<0.05 & logFC < -0.6). (E) Violin plots of interaction distance (bp) for DIs (EF-Endo vs. EF-KD). **** P value < 0.0001 (Unpaired t-test). (F) Example of lost and gained interactions (EF-Endo vs. EF-KD). Loop anchors depicted by horizontal bars (1mb). Stable interactions not shown.

We next asked whether long-range intra-chromosomal interactions were globally altered by the presence of EWS/FLI. We plotted interaction frequency over linear genomic distance for all intra-chromosomal interactions. We found a decrease in long distance (2 – 20 mb) interactions and an increase in shorter distance (0.1 – 1 mb) interactions in EF-WT cells as compared to EF-KD cells (Figure 1B). To determine whether these differences were significant, differential interaction (DI) analysis was performed to identify higher order interactions at low resolution (1 mb bins) that are different in EF-WT as compared to EF-KD cells. DIs were identified and separated into gained or lost interactions (Figures 1C, S1E). We found that more interactions were lost than gained with EWS/FLI expression (Figure 1D) and lost interactions were significantly larger than gained interactions (Figures 1E, S1F). Example genomic regions are shown in Figures 1F and S1G. Taken together, these data demonstrate that expression of EWS/FLI results in a highly-significant reduction of interactions between distant loci in favor of increased interactions between nearby loci.

### Alterations in compartment structure associate highly with enhancer landscape and gene expression changes

The megabase scale changes identified in intra-chromosomal interactions suggest that these may impact large-scale chromatin organizational features, such as chromatin compartment organization. Principal component analysis (PCA) of Hi-C data has been previously shown to identify and distinguish chromatin regions into one of two compartmental states: compartment A (relatively transcriptionally active) and compartment B (relatively transcriptionally inactive; Lieberman-Aiden et al., 2009). To identify A and B compartmental organization for EF-Endo, EF-KD and EF-Rescue cells, PCA of the respective interaction profiles were performed and the first principal component (PC1) values were assigned to each 20 kilobase (kb) region in the genome. Here, positive PC1 values assigned to a region indicate A compartmentalization and negative PC1 values indicate B compartmentalization. We found that the majority of compartmental assignments remained unchanged in the presence or absence of EWS/FLI, but small segments of the genome showed noticeable changes (red highlights, Figure 2A). These findings were confirmed using Pearson correlation analyses (Figure S2A).

**Figure 2.**
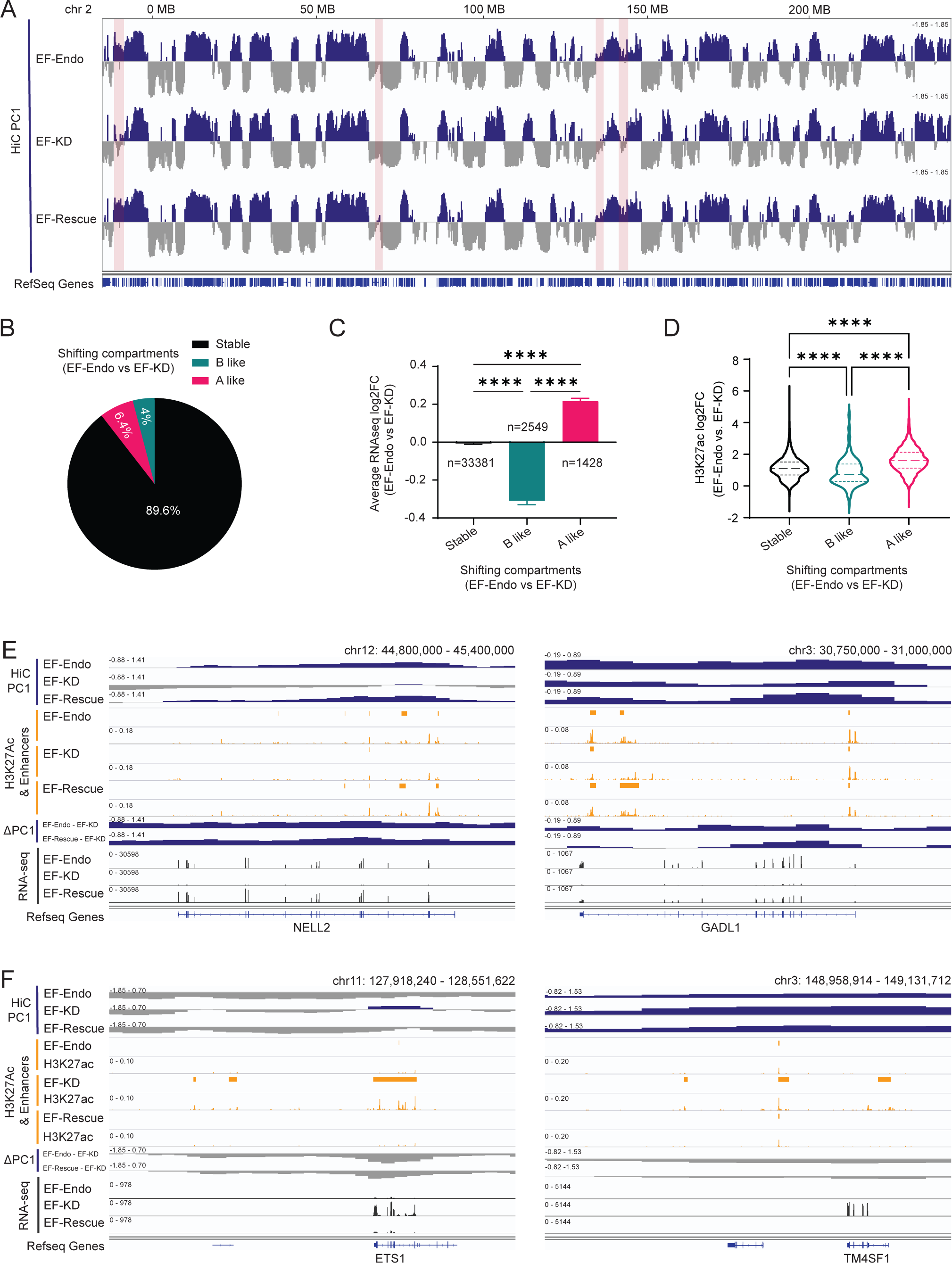
**Alterations in compartment structure associate highly with enhancer landscape and gene expression changes** (A) Hi-C eigenvectors (PC1) for chromosome 2 showing active (A, blue) and inactive (B, gray) compartments in EF-Endo, EF-KD and EF-Rescue. Highlighted regions show compartment alterations in EF-KD compared to EF-Endo and EF-Rescue. (B) Pie chart showing percentage of the genome undergoing compartment shifting in EF-Endo vs. EF-KD. A-like shift, ΔPC1 ≥ 0.4; B-like shift, ΔPC1 ≤ -0.4. (C) Average log_2_ fold-change (log_2_FC) of gene expression at shifting compartments (EF-Endo vs. EF-KD). Mean ± SEM shown. **** P value < 0.0001 (Šídák’s multiple comparisons test). (D) Violin plots showing log_2_FC enrichment of H3K27Ac at shifting compartments for EF-Endo vs. EF-KD. ****P value < 0.0001 (Tukey’s multiple comparisons test). Differential H3K27Ac analysis was performed using diffBind and DESeq2 R packages. (E & F) Integrative genome viewer (IGV) traces showing examples of A-like (positive ΔPC1) (F) and B-like (negative ΔPC1) (G) compartment shifts in EF-Endo and EF-Rescue compared to EF-KD. Corresponding compartment profiles (PC1), enhancers, H3K27Ac levels, and gene expression (RNA-seq) are also shown.

We next identified shifting compartments between EF-WT and EF-KD where regions that show a positive change in PC1 values (ΔPC1 ≥ 0.4) shift into a more “A-like” compartment state while those that show a negative change in PC1 values (ΔPC1 ≤ -0.4) shift to a more “B-like” compartment state. Overall, 10.4% (6.4% A-like and 4% B-like) and 8.7% (3.2% A-like and 5.5% B-like) of the genome showed altered compartments in EF-Endo vs. EF-KD, and EF-Rescue vs. EF-KD, respectively (Figure 2B, S2B).

We next asked whether EWS/FLI-mediated changes in compartmentalization were associated with gene expression and enhancer alterations. We compared previously-published RNA-seq and H3K27Ac CUT&Tag data in A673 cells (Boone et al., 2021; Theisen et al., 2020) to our compartmental data, and found that genes that shifted in an A-like direction displayed significant increases in EWS/FLI-mediated gene expression, while genes mapping to B-like shifts showed significant decreases in EWS/FLI-mediated gene expression (Figure 2C, S2C). Genes in stable compartments demonstrated little change in gene expression. Similarly, H3K27Ac enrichment (a common marker of enhancers) was significantly increased with A-like compartmental shifts while H3K27Ac enrichment was significantly decreased with B-like compartmental shifts when compared to stable compartments (Figure 2D, S2D). Taken together, these results show strong global associations between compartment activation, gain of enhancers, and upregulation of gene expression that can be appreciated at representative gene loci (Figure 2E). Conversely, strong associations between compartment inactivation, enhancer loss, and downregulation of gene expression can be similarly appreciated (Figure 2F).

### EWS/FLI promotes active compartmentalization of chromatin

We next hypothesized that EWS/FLI binding to chromatin might directly affect compartmental changes in Ewing sarcoma. To test this, we first developed a set of “high-confidence” EWS/FLI binding sites by performing genome-wide localization studies for EWS/FLI in A673, TC71, SK-N-MC, and EWS-502 cells or the EWS/ERG fusion in TTC-466 cells using CUT&Tag (Kaya-Okur et al., 2019). We identified 4658 overlapping regions, containing 5637 EWS/FLI peaks in A673, that were conserved across all five lines (Figure S3A). The conserved EWS/FLI peaks in A673 cells were used in all subsequent analyses as a “high-confidence” peak set. Motif analysis of these peaks showed GGAA-μsats and conserved ETS sequences as the top enriched motifs (Figure S3B), as anticipated, thus confirming the quality of this dataset. Peak traces show highly conserved chromatin localization profiles for EWS/FLI and EWS/ERG in all five cell lines (Figure 3A) at both GGAA-μsats (Figure S3C) and conventional ETS sites (Figure S3D). Analysis of EWS/FLI enrichment showed that enhancers that were gained in EF-WT (as compared to EF-KD) had significantly higher EWS/FLI enrichment as compared to stable or lost enhancers, supporting the previously reported role of EWS/FLI in activating enhancer activity (Figures 3B, S3E, S3F; Riggi et al., 2014).

**Figure 3.**
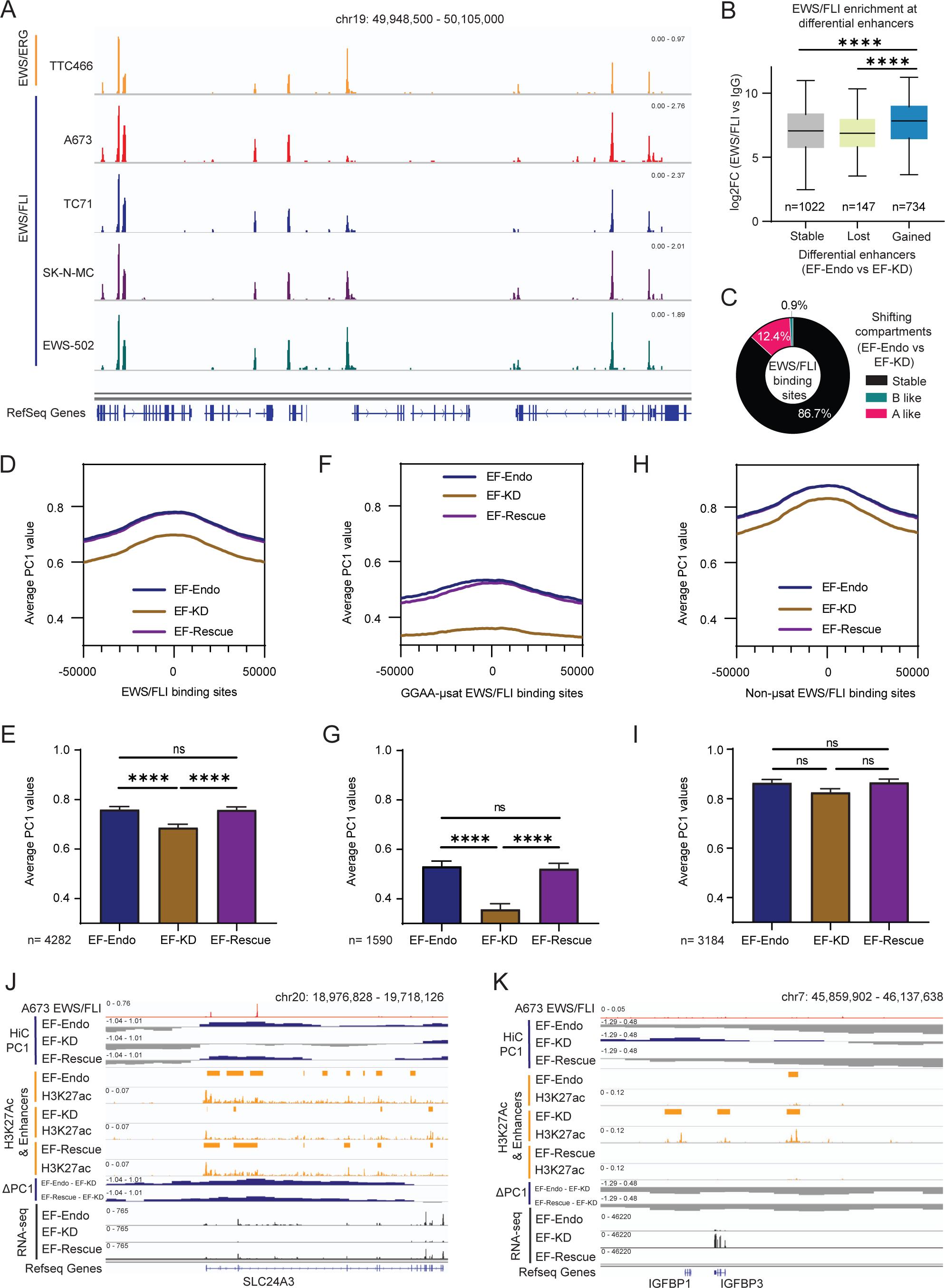
**EWS/FLI promotes active compartmentalization of chromatin** (A) Representative example of EWS/FLI and EWS/ERG localization in 5 Ewing sarcoma cell lines (TTC466, A673, TC71, SK-N-MC and EWS-502) over a 157 kb region of chromosome 19. Tracks represent log_2_ ratio of EWS/FLI or EWS/ERG vs. IgG signal. (B) Boxplots showing EWS/FLI enrichment in A673 cells at differential enhancer regions in EF-Endo vs. EF-KD. **\*\*\*\***P value < 0.0001 (Tukey’s multiple comparisons test). Differential enhancers shown in Figure S3E. (C) Doughnut chart showing proportion of EWS/FLI occupancy in A673 cells at shifting compartment regions between EF-Endo and EF-KD. (D, F, H) Histogram plots of average PC1 values within 50kb of EWS/FLI binding sites in EF-Endo, EF-KD, and EF-Rescue. (D) All conserved EWS/FLI binding sites in A673 cells. (F) EWS/FLI GGAA-μsat binding sites. (H) EWS/FLI non-μsat binding sites. (E, G, I) Bar charts showing average PC1 values in EF-Endo, EF-KD and EF-Rescue for 20kb regions overlapping EWS/FLI peaks. (E) All conserved EWS/FLI binding sites in A673. (G) GGAA-μsat binding sites. (I) Non-μsat binding sites. Mean ± SEM shown. **** P value < 0.0001 (Tukey’s multiple comparisons test). (J, K) IGV traces showing EWS/FLI binding in A673 cells with corresponding compartment profiles, H3K27Ac levels, enhancers, and gene expression in EF-Endo, EF-KD and EF-Rescue. ΔPC1 shows (J) A-like & (K) B-like compartment shifts in EF-Endo and EF-Rescue compared to EF-KD.

To evaluate the relationship between EWS/FLI binding and compartmental shifts, we annotated each conserved EWS/FLI peak to its corresponding shifting compartmental segment in the genome. We found that 86.7% of EWS/FLI peaks bound to stable compartments (including both A and B compartments that remained stable), while 12.4% bound to A-like and only 0.9% bound to B-like compartment shifts in EF-Endo vs. EF-KD (Figure 3C). This suggests a direct role for EWS/FLI in compartment activation, but a minimal role in direct compartment inactivation. Similar results were observed for EF-Rescue vs. EF-KD (Figure S3G). Analysis of average PC1 values (i.e., compartmentalization) within 50 kb of EWS/FLI binding sites demonstrated that EWS/FLI binding was associated with statistically-significant compartment activation (demonstrated as an increase in PC1 values) in both EF-Endo and EF-Rescue compared to EF-KD, with little difference in PC1 values between EF-Endo and EF-Rescue (Figure 3D, 3E). Most compartment activation was associated with EWS/FLI bound at GGAA-μsats (Figure 3F, 3G), with no statistically-significant difference observed at non-μsat sites (Figure 3H, 3I). Representative loci demonstrate compartment activation mediated by EWS/FLI binding, with associated enhancer formation and upregulation of gene expression (Figure 3J, S3H). Conversely, representative loci demonstrate compartment inactivation, associated with loss of enhancers and decreased gene expression (Figure 3K, S3I). These data support a model in which EWS/FLI binding to GGAA-μsats induces new enhancer formation, A-compartmentalization, and subsequent target gene upregulation. Conversely, these data suggest that EWS/FLI seldom has a direct effect on inducing B-compartmentalization and gene repression.

### EWS/FLI perturbs TAD boundaries

Large scale changes in chromatin features demonstrated by alterations to compartments and megabase scale interactions next prompted us to evaluate the effect of EWS/FLI on sub-megabase TAD structures. TADs are known to be highly conserved across different cell types and are usually bounded by CCCTC-binding factor (CTCF) that tends to limit enhancer activity to genes that are present within the same TAD (Pombo and Dillon, 2015; Rao et al., 2014). We used Hi-C data to identify TAD boundaries in EF-Endo, EF-KD and EF-Rescue and found that ∼80% (6129) of TAD boundaries are conserved (stable boundaries), while ∼4% (306) of boundaries were found only in EF-KD and ∼5.8% (441) of boundaries were specific to both the EF-WT conditions (Figure 4A).

**Figure 4.**
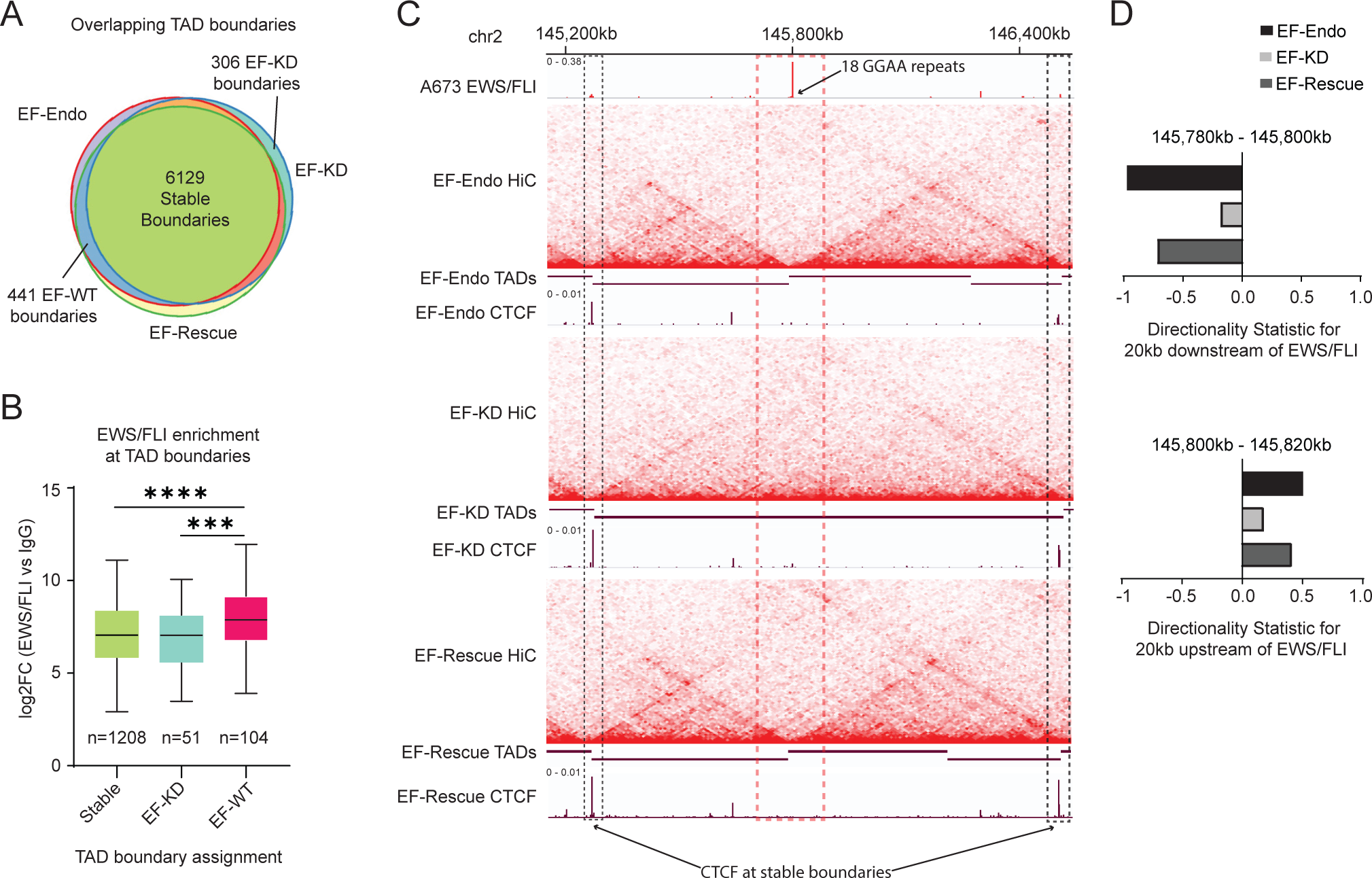
**EWS/FLI perturbs TAD boundaries** (A) Venn diagram overlap of TAD boundaries in EF-Endo, EF-KD, and EF-Rescue. TAD boundaries were identified as 20kb regions that allow minimal contact between its upstream and downstream regions compared to its neighboring regions, P value <0.05. (B) Boxplots showing EWS/FLI enrichment at EF-KD, EF-WT, and stable boundaries in A673. *** P value < 0.001, **** P value < 0.0001 (Tukey’s multiple comparisons test). (C) Example of EF-WT (dashed orange box) and stable boundaries (dashed black boxes) over a 1.4mb region in chromosome 2. Heatmaps show Hi-C interactions for EF-Endo, EF-KD, and EF-Rescue. Intensity of red spots correspond to contact frequency. TAD calls are depicted using maroon horizontal bars where a break corresponds to a TAD boundary. EWS/FLI and CTCF localization are also shown. (D) Directionality statistic (DS) for 20 kb upstream (top panel) and downstream (bottom panel) of EWS/FLI binding site (orange dashed box) in EF-Endo, EF-KD, and EF-Rescue. Positive and negative signs reflect the preference for downstream and upstream interactions, respectively. The absolute value of DS reflects the magnitude of interaction directionality.

Alterations of CTCF binding have been previously shown to disrupt TAD boundaries (Nora et al., 2017; Sun et al., 2018). We therefore asked whether EWS/FLI altered TAD boundaries by altering CTCF binding in Ewing sarcoma cells. We performed CUT&Tag for CTCF in EF-Endo, EF-KD and EF-Rescue cells. Heatmap analysis of CTCF binding sites at boundary regions showed that CTCF occupancy was largely unchanged between EF-Endo, EF-KD and EF-Rescue at all TAD boundaries (stable, EF-WT and EF-KD; Figure S4A). Evaluation of CTCF peak tracks also showed similar CTCF localization in EF-Endo, EF-KD and EF-Rescue (Figure S4B-D).

The similarity of CTCF binding at TAD boundaries suggested a CTCF-independent role in the boundary perturbations we observed in A673 Ewing sarcoma cells. We therefore investigated whether EWS/FLI itself was involved in directly altering TAD boundaries. We found that EWS/FLI enrichment was significantly higher at EF-WT boundaries as compared to stable and EF–KD boundaries (Figure 4B), suggesting a role for EWS/FLI enrichment in formation of TAD boundaries. Examination of representative EF-WT boundaries occupied by EWS/FLI (dashed orange box) showed boundary formation in both EF-Endo and EF-Rescue, as compared to EF-KD as shown by Hi-C heatmaps and TAD calls (Figures 4C, S4E, S4F). Directionality statistic (DS), a measure of the direction and size of the interaction preference calculated for 20 kb regions adjacent to EWS/FLI binding sites, showed higher absolute values, but in opposite directions, for each 20kb region in EF-Endo and EF-Rescue as compared to EF-KD (Figures 4D, S4G, S4H), suggesting an insulating function, similar to that of CTCF, for EWS/FLI at these sites (Dixon et al., 2012; Lun and Smyth, 2015). Furthermore, we do not detect CTCF occupancy at these EWS/FLI-specific boundaries, while CTCF binding clearly demarks the nearby stable boundaries (dashed black boxes). These data suggest a direct role for EWS/FLI in insulating chromatin interactions to promote TAD boundaries at specific loci, independent of CTCF binding.

### EWS/FLI mediates chromatin looping

Identification of EWS/FLI at TAD boundaries and in changing directionality of interactions suggested that EWS/FLI might reorganize chromatin looping in Ewing sarcoma. We identified changes in chromatin looping at a high resolution (20 kb bins) between EF-WT and EF-KD cells. We found 1608 gained and 3878 lost loops in EF-Endo vs. EF-KD (Figure 5A) and 706 gained and 2066 lost loops in EF-Rescue vs. EF-KD (Figure S5A). Comparison of these data to the differential enhancer analysis above (Figure S4E) revealed that gained enhancers showed significant increases in loop enrichment, while lost enhancers showed significant decreases in loop enrichment, supporting a strong association between enhancer establishment and loop enrichment (Figures 5B, S5B). Annotation of differential loops to genes within 20kb of loop anchors demonstrated that lost loops are strongly associated with downregulation of gene expression (Figure 5C, S5C), while gained loops associate strongly with upregulation of gene expression (Figure 5D, S5D).

**Figure 5.**
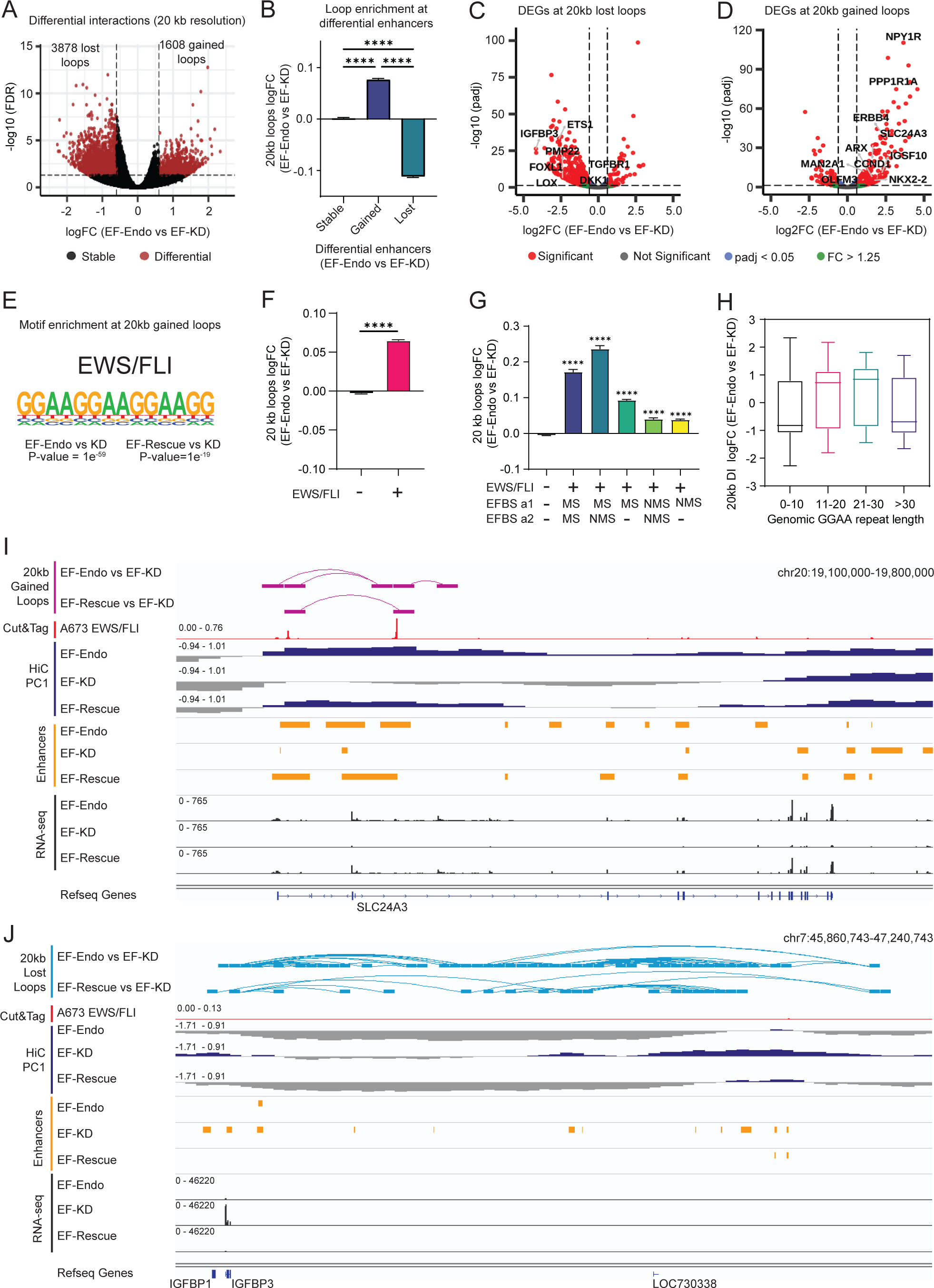
**EWS/FLI mediates chromatin looping** (A) Volcano plot of differential loops (DL) identified at 20 kb resolution in EF-Endo vs. EF-KD A673 cells using diffHiC. DL, FDR < 0.05 and |logFC| > 0.6. (B) Average logFC enrichment of all 20kb loops at differential enhancers for EF-Endo vs. EF-KD. Mean ± SEM are shown. **** P value < 0.0001 (Tukey’s multiple comparisons test). (C) & (D) Volcano plots of differentially expressed genes (DEG) mapping to anchors of (C) lost (FDR<0.05, logFC < -0.6) and (D) gained (FDR<0.05, logFC > 0.6) 20kb loops in EF-Endo vs. EF-KD. Significantly altered genes (red dots): Adjusted P value (padj) < 0.05 and fold change (FC) > 1.25. (E) Top motif enriched at anchors of 20kb gained loops for EF-Endo and EF-Rescue compared to EF-KD. P values for enriched motifs were identified by comparing to matched, randomized background regions using Homer. (F) Average logFC enrichment of 20kb loops in EF-Endo vs. EF-KD associated with an EWS/FLI binding site (+) or no EWS/FLI binding site (-). Mean ± SEM shown. **** P value < 0.0001 (Unpaired t-test). (G) Average logFC (EF-Endo vs. EF-KD) enrichment of 20kb loops associated with the type of EWS/FLI binding site at each loop anchor. EFBS = EWS/FLI binding site, a1= first anchor of a loop, a2=second anchor of a loop, MS= GGAA-μsat bound EWS/FLI, NMS= non-μsat bound EWS/FLI. Mean ± SEM shown. **** P value < 0.0001 (Dunnett’s multiple comparisons test). All comparisons were made to the EWS/FLI (-) loops. (H) Boxplots showing enrichment of DLs (EF-Endo vs. EF-KD) mapping to *total* number of GGAA motifs (sequence between two GGAA motifs ≤ 20-bp). (I, J) Examples showing gained (I, purple) and lost (J, blue) loops in EF-Endo vs. EF-KD and EF-Rescue vs. EF-KD. Corresponding EWS/FLI localization (CUT&Tag) in A673, enhancers, compartment profiles (PC1) and gene expression (RNA-seq) in EF-Endo, EF-KD and EF-Rescue are also shown.

To identify features that promote looping in EF-WT cells, we performed motif analysis of gained loops and found GGAA-μsats as the top enriched motif (Figure 5E), suggesting a direct role for EWS/FLI bound GGAA-μsats in mediating chromatin looping. To determine whether EWS/FLI binding to chromatin promotes loop enrichment, we compared the fold change of interactions either with or without a conserved EWS/FLI peak at an anchor. DNA bound by EWS/FLI showed significantly higher loop enrichment compared to DNA with no EWS/FLI (Figure 5F, S5E), supporting a direct role for EWS/FLI in mediating loop formation.

To interrogate the effect of EWS/FLI specific binding site on loop formation, we sub-categorized EWS/FLI bound loops into 5 categories: (i) GGAA-μsat bound EWS/FLI at both loop anchors (MS-MS); (ii) GGAA-μsat bound EWS/FLI at one anchor and non-μsat bound EWS/FLI at the other anchor (MS-NMS); (iii) GGAA-μsat bound EWS/FLI at one anchor only (MS); (iv) non-μsat bound EWS/FLI at both anchors (NMS-NMS); (v) non-μsat bound EWS/FLI at one anchor only (NMS). Loops in categories (i) and (ii) showed higher enrichment compared to loops in any of the other categories (Figure 5G, S5F), supporting the notion that GGAA-μsat bound EWS/FLI promotes looping to other EWS/FLI bound sites.

To determine whether EWS/FLI-mediated chromatin loops are insulated within TADs, we next identified 143 significantly gained loops in EF-Endo vs. EF-KD, and 43 significantly gained loops in EF-Rescue vs. EF-KD in categories (i) and (ii) combined and compared these to TAD regions. We found 92 (and 28) intra-TAD loops, and 51 (and 15) inter-TAD loops in EF-Endo vs. EF-KD (and EF-Rescue vs. EF-KD). These data not only suggest a direct role for EWS/FLI in formation of loops within TADs, but also indicates that EWS/FLI bound GGAA-μsats can bypass TAD boundaries to interact with other EWS/FLI bound loci. Representative examples are shown in Figure S5G.

Previous work suggested an optimal length of 18–26 total GGAA repeats for GGAA-μsats in EWS/FLI binding and gene expression (Johnson et al., 2017; Monument et al., 2014; Zuo et al., 2021). We therefore asked whether similar GGAA-μsats lengths were also optimal for EWS/FLI-mediated looping. We evaluated the enrichment of differential loops against the underlying number of GGAA-μsat motifs and found the highest level of loop enrichment at 21-30 *total* GGAA repeats (Figures 5H, S6A). We found similar strong correlations between the number of *consecutive* GGAA repeats and differential loop enrichment (Figures S6B, S6C). Taken together, these results show that EWS/FLI mediated loop enrichment is dependent on the number of GGAA motifs underlying the loop anchors, and that EWS/FLI mediated gained loops strongly associate with enhancer and gene activation (Figures 5I, S6D). Conversely, lost loops do not generally associate with EWS/FLI binding but still show strong association with enhancer loss and downregulation of gene expression (Figures 5J and S6E).

### Cell-specific local chromatin structure affects gene regulation

Multiple studies have previously identified putative EWS/FLI target genes (Aynaud et al., 2020; Hancock and Lessnick, 2008; Kinsey et al., 2006; Prieur et al., 2004; Smith et al., 2006). These studies, however, lacked genome-wide chromatin loop data associated with EWS/FLI to accurately annotate target genes to specific EWS/FLI bound regulatory elements. We therefore sought to define a set of “direct EWS/FLI target genes” as those with (1) transcription start sites (TSS) either within 20kb of an EWS/FLI binding site or overlapping an EWS/FLI-bound gained loop anchor (20 kb resolution), and (2) differential expression in EF-Endo vs. EF-KD in A673 cells. We found 707 genes to be directly upregulated and 339 genes to be directly downregulated by EWS/FLI. Tables S1 and S2 list these direct up- and down-regulated genes along with the type of EWS/FLI binding site regulating them. Inspection of the EWS/FLI binding sites regulating gene expression in these data showed that GGAA-usat bound EWS/FLI plays a predominantly activating role by directly upregulating 324 genes compared to directly downregulating 49 genes. Conversely, non-μsat bound EWS/FLI was found to play a more equal role in both activation and repression of genes by directly upregulating 383 genes, and directly downregulating 290 genes.

To investigate whether EWS/FLI directly regulates similar target genes in an alternate Ewing sarcoma cell line, we knocked-down EWS/FLI in TC71 cells and performed RNA-seq for EWS/FLI WT (EF-Endo) and EWS/FLI knock-down (EF-KD) cells (Figures S7A-C). PCA analysis revealed strong clustering of RNA-seq replicates in the first principal component (PC1) axis (79% of the variance in the data; Figure S7D). Upon differential gene expression analysis, we found that the total number of genes regulated by EWS/FLI in TC71 was lower than in A673 (2,179 versus 4,648, respectively; Figure S7E), likely due to the less efficient depletion of EWS/FLI (compare Figures S7A-C to Figures S1A-C). GSEA analysis using the direct up- and down-regulated gene sets in A673 showed significant enrichment for the same genes in TC71 cells (NES=+2.76 and -2.16 for up- and down-regulated gene sets respectively), demonstrating that many genes are regulated similarly by EWS/FLI in both cell lines (Figures 6A, 6B). However, we also observed critical direct targets in A673, e.g. *NKX2-2*, that were regulated differently in TC71. In A673, depletion of EWS/FLI results in depletion of *NKX2-2* expression and loss of oncogenesis, but not in TC71 (Figure 6C; Smith et al., 2006). Since EWS/FLI localization is conserved between A673 and TC71 at the *NKX2-2* locus, we asked whether heterogeneity in regulation was a result of altered chromatin structure between the two cell lines. Hi-C data in A673 showed EWS/FLI bound GGAA-μsat looping to *NKX2-2* to regulate gene expression (Figure 6D). We therefore performed 4C experiments to compare the local chromatin structure of TC71 and A673 cells at the *NKX2-2* locus, using the right most GGAA-μsat (RMS) as the viewpoint (Figure 6E). We observed high interaction intensity, possibly within an insulated domain in both A673 and TC71, beyond which interaction intensity drops (boundary highlighted in yellow). However, in A673, the intensity of interactions at this boundary is much depleted compared to TC71, allowing EWS/FLI to significantly interact near *NKX2-2* in A673 (black boxes). No significant interactions between RMS and *NKX2-2* were detected in TC71, suggesting direct EWS/FLI regulation of *NKX2-2* in A673 but not in TC71. CTCF localization showed increased insulation of EWS/FLI interactions at the CTCF peak upstream of the *NKX2-2* locus (highlighted in yellow) in TC71, which is largely absent in A673, allowing EWS/FLI bound GGAA-μsat to loop to *NKX2-2* in EF-Endo and EF-Rescue A673 cells. These data demonstrate that local chromatin structure, such as CTCF localization and chromatin loop formation, may account for differences in EWS/FLI-mediated gene regulation in different Ewing sarcoma cell lines and tumors.

**Figure 6.**
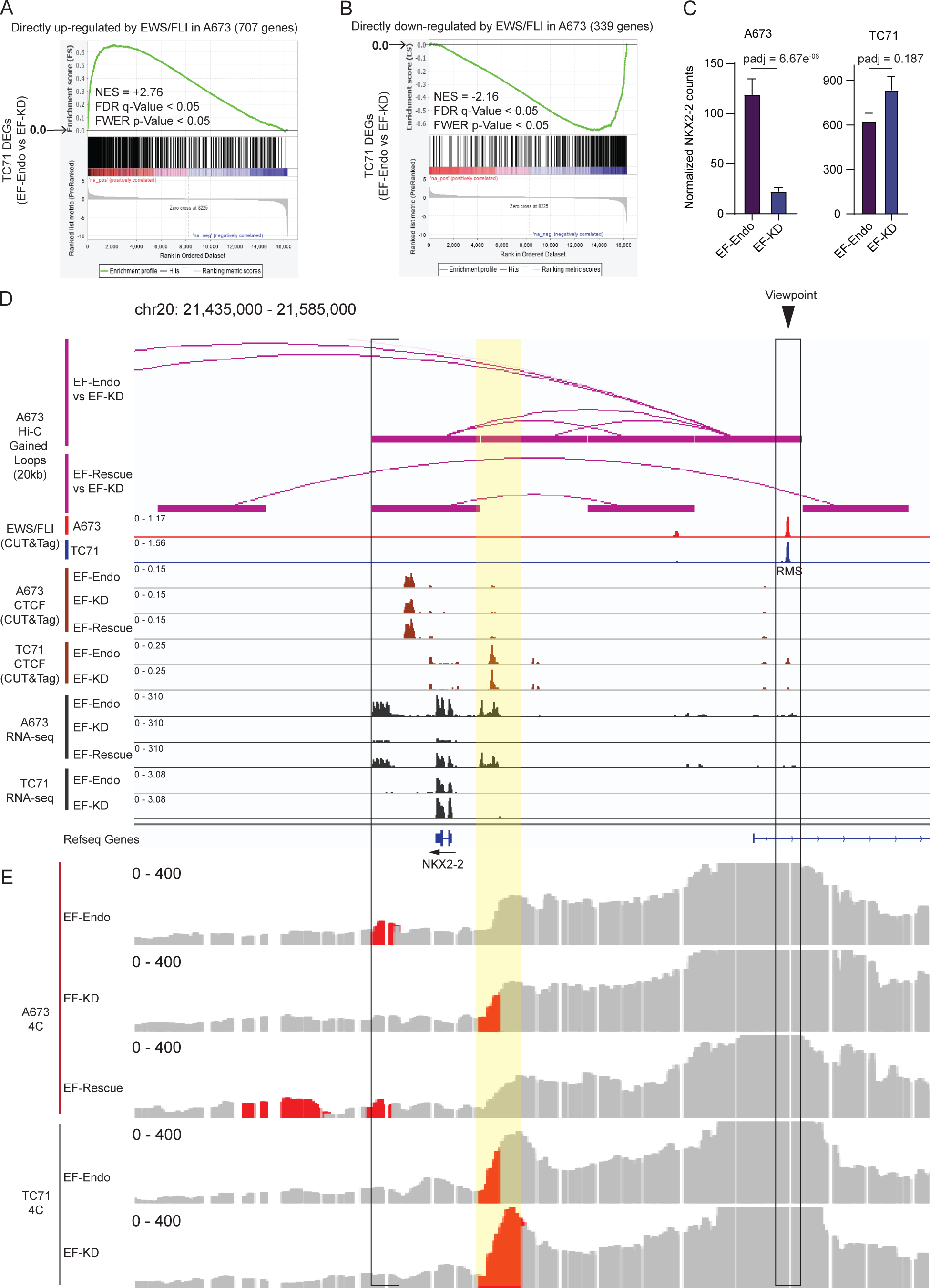
**Cell-specific local chromatin structure affects gene regulation** (A, B) GSEA plots showing functional enrichment of genes directly (A) upregulated or (B) downregulated by EWS/FLI in EF-Endo vs. EF-KD A673 cells to rank ordered DEGs in TC71 cells (EF-Endo vs. EF-KD). Significant, |NES|> 1.5, FDR q-value < 0.05 and FWER p-value < 0.05. (C) Normalized *NKX2-2* expression counts from RNA-sequencing in A673 and TC71 EF-Endo and EF-KD cells. Adjusted p-value (padj) was calculated using DeSeq2 R package. Significant, padj < 0.05. (D) Altered regulation of *NKX2-2* in A673 compared to TC71. IGV traces show gained loops at 20kb resolution for EF-Endo vs. EF-KD and EF-Rescue vs. EF-KD in A673 (purple). CUT&Tag data show EWS/FLI and CTCF occupancy at the *NKX2-2* locus in A673 and TC71. RNA sequencing traces show gene expression for EF-Endo, EF-KD and EF-Rescue in A673 and EF-Endo and EF-KD in TC71. (E) 4C data showing significant loops (in red) (P value < 0.05) with the viewpoint (region of interest; RMS, Right GGAA-μsat) in A673 and TC71. Significance testing performed using non-parametric statistics by ranking 4C signal compared to a background model.

**Figure 7.**
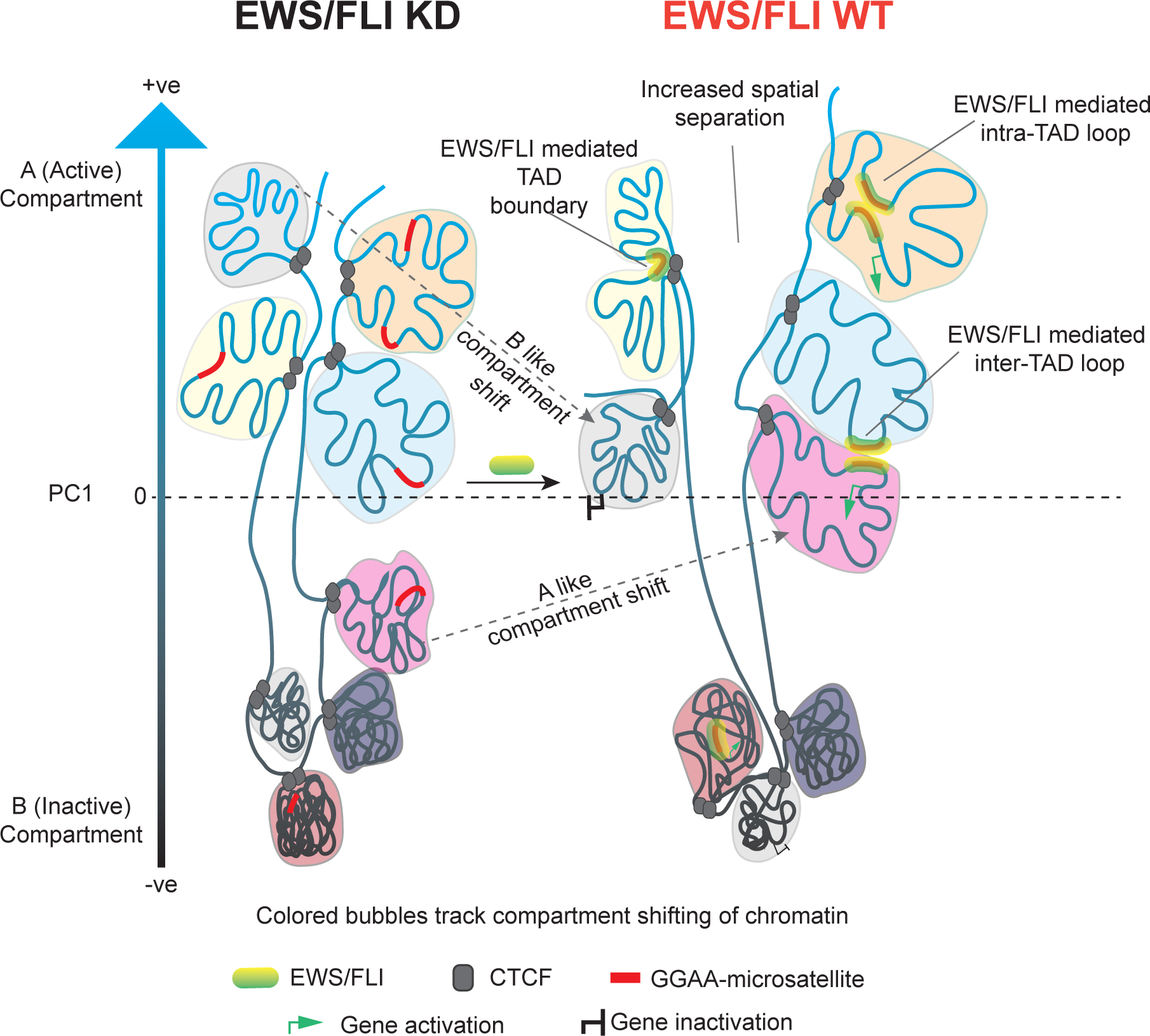
**Model illustrating EWS/FLI mediated chromatin alterations**

## DISCUSSION

Our study of 3D chromatin organization revealed widespread reprogramming of higher order chromatin interactions, A/B compartments, TAD boundaries, and enhancer-promoter type chromatin loops by EWS/FLI in patient derived Ewing sarcoma cells. Chromatin changes that occur upon depletion of EWS/FLI can be largely reversed by re-expression of the “rescue” EWS/FLI. These results provide evidence for significant EWS/FLI mediated remodeling of 3D chromatin to promote a transcriptional state specific to Ewing sarcoma.

We highlight a role for EWS/FLI bound GGAA-μsats in directly anchoring chromatin loops (both inter- and intra-TAD loops) and TAD boundaries. EWS/FLI binding to GGAA-μsats also promotes active compartmentalization of chromatin and has a strong looping preference for other EWS/FLI bound DNA. This in turn may promote active compartmentalization of DNA interacting with GGAA-μsat bound EWS/FLI, thus upregulating gene expression in these compartments. Recent research suggest that STAG2 promotes extrusion of CTCF-anchored loops in Ewing sarcoma, whereby a subset of EWS/FLI-bound enhancers can undergo intra-TAD looping to alter gene expression (Adane et al., 2021; Surdez et al., 2021). Loop extrusion is a mechanism by which insulated chromatin neighborhoods and intra-TAD loops are formed (Fudenberg et al., 2016). Identification of EWS/FLI mediated inter-TAD looping in our study therefore suggests a role for mechanism(s), other than STAG2 mediated loop extrusion, by which EWS/FLI promotes inter-TAD loops that do not respect TAD boundaries. Investigation into alternative mechanisms, e.g. phase separation and/or weak multivalent interactions, are therefore necessary to better understand the role of EWS/FLI in chromatin looping.

Past and current work have shown more genes are downregulated than upregulated in the presence of EWS/FLI in Ewing sarcoma (Prieur et al., 2004; Smith et al., 2006). In line with this observation, several studies show direct mechanisms for EWS/FLI in transcriptional repression (Riggi et al., 2014; Sankar et al., 2013). In contrast, our research provides evidence for indirect modes of chromatin restructuring by EWS/FLI, such as compartment inactivation and loop loss, that are directly associated with enhancer loss and gene repression. This suggests involvement of factors, other than EWS/FLI, in enhancer depletion, loop loss, compartment inactivation, and gene repression. It is therefore important to interrogate these regulatory elements to identify critical factors involved in gene repression and to understand the role of EWS/FLI in indirectly promoting such repressive chromatin states in Ewing sarcoma.

EWS/FLI alters the transcriptome by directly regulating hundreds of genes, which in turn leads to dysregulation of thousands of downstream targets, leading to oncogenesis. Hence, identification of direct EWS/FLI target genes is crucial to define the primary set of genes that initiate this transcriptional mayhem in Ewing sarcoma. Using EWS/FLI mediated chromatin looping data, we fill a significant knowledge gap in Ewing sarcoma by directly annotating EWS/FLI regulatory elements to target genes that trigger this wide range of transcriptional alteration. Our list of direct EWS/FLI target genes is however, limited by the resolution of Hi-C experiments as well as the reference genome used. Publicly available reference genomes do not include structural and small variants present in cancer cells (Ciriello et al., 2013; Futreal et al., 2004). Existence of any such variants at repetitive GGAA-μsats in Ewing sarcoma cells would be of great consequence to EWS/FLI binding, enhancer establishment, A/B compartmentalization, and chromatin looping. Hence, construction of custom reference genomes along with higher resolution looping data for each cell line are necessary to accurately identify the complete set of direct EWS/FLI target genes in Ewing sarcoma. We detected heterogeneity in regulation of these direct target genes in two distinct patient derived Ewing sarcoma cell lines, suggesting that factors other than EWS/FLI localization are at play in promoting such altered regulation. GGAA-μsat polymorphisms have been linked to altered gene expression and disease susceptibility in Ewing sarcoma (Grunewald et al., 2015; Monument et al., 2014). Here we show that heterogenous regulation of genes in different Ewing sarcoma cells may also occur as a result of altered local chromatin structure, suggesting that such disparity in chromatin organization may also effect disease susceptibility, clinical presentation, therapeutic response, and patient outcomes.

In conclusion, our study demonstrates that the EWS/FLI fusion oncoprotein plays a significant role in chromatin reorganization in Ewing sarcoma. It is therefore likely that other similar fusion oncoproteins found in different cancers may also be involved in reogranization of chromatin to promote oncogenesis. Although not studied at multiple levels of organization, recent research show that the NUP98-HOXA9 oncogenic fusion transcription factor, found in leukemias, can undergo phase transition and promote chromatin looping to regulate gene expression and oncogenesis (Ahn et al., 2021). Our results therefore provide direct evidence for a significant role for fusion oncoproteins in reprogramming the 3D chromatin landscape to promote an altered transcriptional state in a fusion mediated cancer.

## MATERIALS AND METHODS

### Cell lines and culture conditions

HEK-293EBNA, female, human (Invitrogen). Cells were cultured in DMEM (Corning Cellgro 10-013-CV) supplemented with 10% heat inactivated fetal bovine serum (FBS) (Gibco 16000-044), 1% penicillin/streptomycin/glutamine (PSQ) (Gibco 10378-016), and 0.3mg/mL geneticin (Gibco 10131-027) at 37°C and 5% CO_2_. Cell line was authenticated by short tandem repeat (STR) profiling through Genetica (LabCorp).

A673, female, human (ATCC). Cells were cultured in DMEM with L-Glutamine, supplemented with 10% FBS, 1% PSQ and 1% sodium pyruvate (Gibco 11360-070) at 37°C and 5% CO_2_. Cell line was authenticated by short tandem repeat (STR) profiling through Genetica (LabCorp).

TC71, male, human (E.S. Kleinerman, The University of Texas, M. D. Anderson Cancer Center, Houston, Texas, USA). Cells were cultured in RPMI 1640 without L-Glutamine, supplemented with 10% FBS and 1% PSQ at 37°C and 5% CO_2._ Cell line was authenticated by short tandem repeat (STR) profiling through Genetica (LabCorp).

TTC-466, female, human (T. Triche, Children’s Hospital Los Angeles, California, USA). Cells were cultured in RPMI 1640 without L-Glutamine, supplemented with 10% FBS and 1% PSQ at 37°C and 5% CO_2_. Cell line was authenticated by short tandem repeat (STR) profiling through Genetica (LabCorp).

EWS-502, sex unspecified, human (J.A. Fletcher, Brigham & Women’s Hospital). Cells were cultured in RPMI 1640 without L-Glutamine, supplemented with 15% FBS and 1% PSQ at 37°C and 5% CO_2_. Cell line was authenticated by short tandem repeat (STR) profiling through Genetica (LabCorp).

SK-N-MC, female, human (ATCC). Cells were cultured in DMEM with L-Glutamine, supplemented with 10% FBS and 1% PSQ at 37°C and 5% CO_2_. Cell line was authenticated by short tandem repeat (STR) profiling through Genetica (LabCorp).

### Constructs and retroviruses

Oligonucleotide sequences for shRNAs targeting either luciferase (luciferase-RNAi, iLuc forward primer: 5’- GATCCCCCTTACGCTGAGTACTTCGATTCAAGAGATCGAAGTACTCAGCGTAAGTTTTTGGAAC-3’; iLuc reverse primer: 5’- TCGAGTTCCAAAAACTTACGCTGAGTACTTCGATCTCTTGAATCGAAGTACTCAGCGTAAGGG G-3’) or the 3’-UTR of endogenous *EWS/FLI* mRNA (EF-2-RNAi, iEF forward primer: 5’-GATCCCCATAGACCTGGGAAGCTTATTTCAAGAGAATAAGCTTCCCACCTCTATTTTTTGGAAC-3’; iEF reverse primer: 5’- TCGAGTTCCAAAAAATAGAGGTGGGAAGCTTATTCTCTTGAAATAAGCTTCCCACCTCTATGGG-3’) were previously cloned downstream of the H1 promoter of the puromycin resistant pSRP RNAi retroviral vector as described (Smith et al., 2006). A 3X-FLAG tagged EWS/FLI cDNA (type 4 breakpoint), was previously cloned into the pMSCV-Hygro retroviral vector as described (Kinsey et al., 2006; Smith et al., 2006).

### Antibodies

Antibodies used: M2-anti-FLAG (Sigma F3165), anti-FLI1 (Abcam ab15289), anti-H3 total (D1H2; Cell Signaling Technology 4499), Rabbit anti-mouse IgG (Abcam ab46540), anti-CTCF (Millipore, 07-729), guinea pig anti-rabbit IgG (Antibodies-Online ABIN101961), IRDye® 800CW goat-anti-rabbit IgG (LI-COR Biosciences 926-32211), IRDye® 680LT goat anti-rabbit IgG (LI-COR Biosciences 926-68021), and anti-rabbit secondary (HRP Cell Signaling 7074), recombinant anti-ERG antibody.

### EWS/FLI knock-down/rescue and selection

EWS/FLI knock-down was performed as described previously (Smith et al., 2006) via retroviral infection with shRNA targeting the 3’ UTR of *EWSR1-FLI1* transcript (iEF2); shRNA targeting luciferase was used as control for knock-down (iLuc). Rescue of EWS/FLI knock-down was carried out using either a 3X-FLAG tagged EWS/FLI cDNA construct devoid of 3’ UTR (3X-FLAG-EF) or an empty pMSCV-hygro vector (e.v) 4 hours after infection with iEF2 to generate the EF-Rescue and EF-KD cells respectively. The control iLuc infection was rescued with e.v to generate the EF-Endo cells expressing endogenous levels of EWS/FLI. Cells were kept in A673 culture media and double selection in A673 media supplemented with 2μg/mL puromycin (Sigma P8833) and 150μg/mL hygromycin B (Thermo Fisher 10687010) was started 2-3 days post infection with the rescue constructs. Cells were selected for 7 to 11 days before harvesting.

EWS/FLI knock-down in TC71 cells was performed via retroviral infection using the iEF2 construct (EF-KD); the iLuc construct was used as a shRNA control (EF-Endo). Cells were grown in TC71 media for 2 days and selection was started in 2μg/mL puromycin. Cells were selected for 2 days and harvested.

### RNA isolation and qRT-PCR

Total mRNA was extracted from frozen cell pellets using the RNeasy Extraction Kit (Qiagen 74136). Reverse transcription and qPCR was performed using the iTaq^TM^ Universal SYBR® Green 1-Step Kit (BioRad 1725151) on a Bio-Rad CFX Connect Real-Time System. Fold change was determined relative to the control sample after normalization to the internal control gene *RPL30*. Primer sequences are as follows: *EWS/FLI* Forward: 5’-CAGTCACTGCACCTCCATCC *EWS/FLI* Reverse: 5’-TTCATGTTATTGCCCCAAGC *RPL30* Forward: 5’-GGGGTACAAGCAGACTCTGAAG *RPL30* Reverse: 5’-ATGGACACCAGTTTTAGCCAAC

### Nuclear protein isolation

Nuclear proteins were isolated from fresh cell pellets re-suspended in 500 μL hypotonic buffer (20 mM HEPES [pH 8.0], 10% glycerol, 10 mM NaCl, 1.5 mM MgCl2, 1 mM EDTA, and 1 mM DTT) with protease inhibitor for 15 minutes. 12.5 μL of IGEPAL® CA-630 (Sigma I8896) was added (final volume of 0.5%) and cells vortexed vigorously for 10 seconds to lyse the cytoplasm. Nuclei were pelleted at 1000 rcf for 5 minutes at 4°C and washed in 20 mM HEPES [pH 8.0], 10% glycerol, 140 mM NaCl, 1.5 mM MgCl2, 1 mM EDTA, 1 mM DTT, and 1% IGEPAL® CA-630 with 1X protease inhibitor. Nuclei were pelleted at 1000 rcf for 5 minutes at 4°C, snap frozen, and stored at -80°C. Proteins were extracted by incubation with RIPA buffer and protease inhibitor cocktail (Roche, 4693159001) on ice for 1 hour with occasional vortexing. Lysates were clarified by centrifugation at max speed for 1 hour at 4°C. Protein concentration was measured using a BCA assay (Thermo Fisher 23225).

### Western blot and densitometry

Protein samples were prepared in 1X SDS loading buffer and boiled for 5 minutes prior to gel loading. Western blots were run on 7.5% Mini-PROTEAN® TGX™ Precast Protein Gels (Bio-Rad, 4561024) for 10 minutes at 90V and 45 minutes at 120V. Proteins were blotted onto nitrocellulose using the iBlot™2 (Thermo Fisher) and developed using LiCor Odyssey CLx Infrared Imaging System.

Protein bands were converted to 8bit images and intensities were calculated using ImageJ 1.51j8. EWS/FLI band intensities were normalized to the respective Histone H3 band as a fraction of the EF-Endo samples for both A673 and TC71 cells.

### CUT&Tag experiment

CUT&Tag experiments were performed as described (Kaya-Okur et al., 2019) with slight modifications. All CUT&Tag experiments were performed in duplicate. EWS/FLI CUT&Tag was performed in A673, TC71, EWS502 and SKNMC cells. EWS/ERG CUT&Tag was performed in TTC-466 cells. CTCF CUT&Tag was performed for A673 EF-Endo, EF-KD and EF-Rescue cells and TC71 EF-Endo and EF-KD cells. H3K27Ac CUT&Tag data was previously performed by (Theisen et al., 2020). BioMag® Plus Concanavalin A-coated magnetic beads (ConA beads, Bangs Laboratories, BP531; 10 µL beads per sample) were washed twice with Binding buffer (20 mM HEPES-KOH pH 7.9, 10 mM KCl, 1 mM CaCl_2_, 1 mM MnCl_2_). 250,000 cells per sample were washed twice with Wash buffer (20 mM HEPES-NaOH pH 7.5, 150 mM NaCl, 0.5 mM Spermidine, 1X protease Inhibitor) and rotated with 10µL ConA beads for 10 minutes at room temperature (RT). Beads bound to cells were separated using magnet stand and re-suspended in 100 µL Antibody buffer (20 mM HEPES-NaOH pH 7.5, 150 mM NaCl, 0.5 mM Spermidine, 0.05% digitonin, 2 mM EDTA, 1x protease Inhibitor). Antibodies (FLI; ERG; CTCF; Rabbit anti-mouse IgG) were added at a dilution of 1:100. Samples were rotated overnight at 4°C. The samples were cleared on a magnet stand and beads were washed with Dig-wash buffer (20 mM HEPES-NaOH pH 7.5, 150 mM NaCl, 0.5 mM Spermidine, 0.05% digitonin. Beads were re-suspended in 100 µL Dig-wash buffer and incubated with guinea pig anti-rabbit IgG at a dilution of 1:100 on a rotator for 1 hour at 4°C. After 3 washes with Dig-wash buffer, beads were re-suspended in 100 µL Dig-300 buffer (20 mM HEPES-NaOH pH 7.5, 300 mM NaCl, 0.5 mM Spermidine, 0.01% Digitonin) with a 1:250 dilution of Protein A-Tn5 transposase fusion protein containing *E.coli* DNA. Samples were rotated for 1 hour at RT. After 3 washes with Dig-300 buffer, beads were re-suspended in 300 µL tagmentation buffer (Dig-300 buffer with 10 mM MgCl_2_) and incubated for 1 hour at 37°C. Tagmentation was stopped by adding 10 µL 0.5 M EDTA, 3 µL 10% SDS, and 2.5 µL 20 mg/mL Proteinase K to each sample, vortexing 5 seconds, and incubating for 1 hour at 50°C. DNA was purified using phenol-chloroform extraction and ethanol precipitation. DNA pellet was dried and re-suspended in 30 µL 10 mM Tris-Cl, pH 8 with 1 mM EDTA and 1/400 RNase A and incubated at 37°C for 10 min. Libraries were amplified using dual-indexed primers. 21 µL of DNA, and 2 µL of each primer (10 µM) were added to 25 µL of NEBNext HiFi 2X PCR master mix. Libraries were amplified using the following cycling conditions: 72°C for 5 min, 98°C for 30 seconds, 15 cycles of 98°C for 10 seconds and 63°C for 10 seconds, 72°C for 1 minute. Amplified libraries were purified by adding 55 µL Agencourt AMPure XP magnetic beads (Beckman Coulter, A63880) to the PCR reactions, incubating 15 minutes, washing twice with 400 µL 80% ethanol, drying the DNA pellet, and eluting purified libraries in 25 µL Tris-Cl, pH 8. Illumina HiSeq4000 system (Nationwide Children’s Hospital Institute for Genomic Medicine) was used to sequence the CUT&Tag libraries using 2X 150 bp paired-end (PE) run.

### CUT&Tag data processing

Quality control on raw sequencing reads were performed using FastQC (v0.11.4; Andrews, 2010). Adapter sequences and/or low quality reads were trimmed using Trim Galore (0.4.4_dev; Krueger, 2015). Reads were aligned to human (hg19) and spike-in *Escherichia coli* (Escherichia_coli_K_12_DH10B NCBI 2008-03-17) genomes using Bowtie2 (v2.3.4.3; Langmead and Salzberg, 2012; Langmead et al., 2019) with the following options ’--no-unal --no-mixed --no-discordant --dovetail --phred33 -q -I 10 -X 700’. The ‘--very-sensitive’ option was added when aligning to spike-in genome. SamTools (v1.9; Danecek et al., 2021) was used to convert sam to bam with ‘-bq 10’ option. CUT&Tag reads were spike-in normalized using DESeq2 median ratio method (Love et al., 2014) in R (v4.0.0; Team, 2020a) to eliminate bias across different samples, minimize the effect of outliers, and appropriately account for global occupancy changes in. Spike-in normalized bigwig tracks were generated and averaged across biological replicates using deepTools (Ramirez et al., 2016). Peaks were called with spike-in normalization using corresponding IgG as controls, and accounting for variation between the biological replicates using MACS2 (v 2.2.7.1; Zhang et al., 2008), DiffBind (v2.14.0; Ross-Innes et al., 2012; Stark R., 2011) and DESeq2. All duplicate reads were kept in the analysis. Irreproducible Discovery Rate (IDR; v 2.0.3; Li et al., 2011) was used to identify reproducible and consistent peaks across replicates. To ensure high quality peaks that are most likely to represent biological signals, the final peak lists were generated using custom thresholds for IDR, log_2_FC, mean normalized counts of signal, and False Discovery Rate (FDR). Default parameters: IDR < 0.01, log_2_FC > 3, mean normalized counts of signal > 80, and FDR < 0.05. Bigwig tracks for each experiment was further compared to the respective IgG control bigwigs using bigwigCompare from deepTools package to generate tracks of log_2_ ratio between experimental and IgG CUT&Tag signal.

### H3K27Ac CUT&Tag analysis

H3K27Ac peak calling: peaks were called after spike-in normalization using the following parameters: FDR < 0.05, IDR < 0.005, mean counts ≥ 50, and log_2_FC ≥ 3.

H3K27Ac differential binding analysis: DiffBind and DESeq2 were used to identify differential H3K27Ac localization for EF-Endo vs. EF-KD and EF-Rescue vs. EF-KD from spike-in normalized H3K27Ac data.

Enhancers and super enhancer detection: Ranked order of super-enhancers (ROSE; Loven et al., 2013; Whyte et al., 2013) was used to identify all enhancers (including both typical enhancers and super-enhancers) from H3K27Ac localization data using the following command: python ROSE_main.py -g hg19 -i H3K27ac.gff -r H3K27ac_sorted.bam -c IgG_sorted.bam -o ROSE_output -t 2500. Peak files for H3K27Ac (.narrowPeak) were converted to GFF (.gff) file format. BAM files generated for each replicate during peak calling were merged, sorted and indexed using SAMtools for both H3K27Ac and IgG controls for use in ROSE analysis. Enhancer BED files from ROSE and H3K27Ac BIGWIG files were visualized using Integrative genomics viewer (IGV; Robinson et al., 2011).

Differential enhancer detection: ROSE was used to identify all enhancer regions (including super enhancers) for each H3K27Ac replicate separately. Differential enhancer regions for EF-Endo vs. EF-KD and EF-Rescue vs. EF-KD were subsequently identified using the DiffBind R package. DESeq2 was used to normalize read counts and identify enhancer regions that are differentially enriched for H3K27Ac signals (FDR < 0.05). Enhancers with significantly increased H3K27Ac signal were identified as gained enhancers (FDR <0.05 & log_2_FC > 0), while enhancers with significantly decreased H3K27Ac signal were identified as lost enhancers (FDR <0.05 & log_2_FC < 0). Enhancers with no significant change in H3K27Ac signal were identified as stable enhancers (FDR

> 0.05).

### EWS/FLI and EWS/ERG CUT&Tag analysis

EWS/FLI and EWS/ERG peaks were called after spike-in normalization using minimal cut-off parameters: FDR < 0.05, IDR < 0.05, mean counts ≥ 1, and log_2_FC ≥ 2 to identify all putative binding locations for EWS/FLI and EWS/ERG in each cell line. A highly conversed list of EWS/FLI and EWS/ERG peaks were identified by overlapping peaks from all 5 cell lines (A673, SK-N-MC, EWS-502, TC71 and TTC-466) using R packages ChIPpeakAnno, Genomic Ranges, VennDiagram and Vennerable (Chen, 2018; Lawrence et al., 2013; Swinton, 2020; Zhu et al., 2010). Regions overlapping peaks in all 5 cell lines were then extracted and overlapped with EWS/FLI peak calls in A673 cell line using BedTools’ intersect tool to identify a list of highly conserved EWS/FLI peaks (Quinlan and Hall, 2010). Homer motif finding tool was used to determine enriched motifs associated with this list of conserved EWS/FLI peaks utilizing the findMotifsGenome.pl script (Heinz et al., 2010). EWS/FLI peaks at differential enhancer regions were identified using BEDTools’ intersect tool.

### RNA-sequencing experiment, data processing, and analysis

RNA-sequencing was performed on two biological replicates each of EF-Endo and EF-KD TC71 samples. Total mRNA were extracted from frozen cell pellets using RNeasy Extraction Kit (Qiagen 74136) and submitted to the Nationwide Children’s Hospital Institute for Genomic Medicine for RNA quality measurements (RIN and DV200), library preparation, sequencing, and differential gene expression analysis for EF-Endo vs. EF-KD in TC71 cells. TruSeq Stranded mRNA Kit (Illumina Cat. No. 20020594) was used to prepare cDNA libraries from total RNA and sequenced on Illumina HiSeq4000 using 2X 150-bp PE run. Low-quality reads (q<10) were filtered out and adaptor sequences trimmed from raw reads using bbduk (v37.64). Each sample was aligned to HumanG1Kv37 assembly of the Homo Sapiens reference (hg19) using STAR version 2.6.0c (Dobin et al., 2013) and analyzed for differential gene expression between EF-Endo and EF-KD in TC71 cells using DESeq2. RNA-seq bigwig tracks were generated and averaged across biological replicates using bamCompare from deepTools package. Differential gene expression data for EF-Endo vs. EF-KD and EF-Rescue vs. EF-KD in A673 cells were available in the laboratory and recently published by (Boone et al., 2021). Volcano plots of DEGs were generated using EnhancedVolcano R package (Blighe et al., 2020).

### *In situ* Hi-C experiment

*In situ* Hi-C was performed in duplicate for EF-Endo, EF-KD and EF-Rescue A673 cells from 2 million frozen cells per replicate fixed in 1X PBS with 2% formaldehyde at RT for 10 minutes and quenched with 0.250M glycine at RT for 5 minutes and on ice for 15 minutes. Crosslinked cells were stored in -80°C and later processed using the Arima Hi-C kit (A410030) as per the kit protocol to obtain proximally-ligated DNA libraries. 100μL proximally-ligated DNA was fragmented using Covaris water bath sonicator (Peak Power = 70W, Duty Factor = 20%, Cycles per burst = 1000, Duration = 35 seconds) to obtain peak fragment size of 395-418bp. Fragments were size selected and biotin enriched per Arima Hi-C protocol. Dual indexed library preparation and amplification was performed using the KAPA Hyper Prep Kit with Library Amplification Module (KK8500, KK8502) according to the Arima Hi-C Kit to obtain libraries with 505-525bp average fragment size. Index sequences are listed in supplementary table S3. *In situ* Hi-C libraries were sequenced on Illumina Hi-Seq 4000 (Nationwide Children’s Hospital Institute for Genomic Medicine) using 2X 150bp PE runs.

### Hi-C data processing

Hi-C PE reads were mapped to Genome Reference Consortium Human Build 37, GRCh37 (hg19) reference genome using publicly available Arima Genomics Mapping pipeline (https://github.com/ArimaGenomics/mapping_pipeline): BWA-MEM (Li, 2013) was used to align the Hi-C PE reads to hg19. PE reads were first mapped independently as single-ends and then paired. Chimeric reads (mapping to the ligation junction) were filtered to retain the read that maps in the 5’-orientation. The filtered single-end Hi-C reads were paired, sorted and mapping quality filtered (mapping quality >10) to get PE BAM files. PCR duplicates were removed using Picard Tools (Institute, 2019). Hi-Csummary files (.hicsum) were generated from BAM (.bam) files using the publicly available iHiC pipeline (Hu, 2020): PE reads in BAM files were converted to BEDPE format using SAMTools. BEDPE files were then used to generate Hi-C Summary files (.hicsum) using the *iHiC_BEDPE2HiCSummary* function. Hi-C summary files were fed into the Homer Hi-C analysis pipeline to generate Homer style tag directories that also generates interaction frequency files (Heinz et al., 2018). These tag directories were used for downstream compartmental analysis. The *iHiC_BEDPE2III* tool from iHiC was used to create an **i**ntra-chromosomal **i**nteraction **i**ndex (III) file recording the number of PE tags (PETs) for 20kb genomic bins that interact within the same chromosome. Hi-C contact matrices (.hic files) were generated using Juicer for SLURM using the command: *juicer.sh -g hg19 -D path_to_juicer_directory -s MboI -p hg19.chrom.sizes –y path_to_restriction_site_file.txt -z path_to_reference_genome/genome.fa (Durand et al., 2016b)*. Hi-C contact maps were visualized using Juicebox (v1.9.8; Durand et al., 2016a).

### A/B compartment detection

The Homer Hi-C analysis package was used to perform Principal Component Analysis (PCA) of Hi-C data to identify chromosomal compartments using PC1 values. The runHiCpca.pl script was used with resolution 20kb, window size 40kb, and genome hg19 to generate the PC1 bedGraphs, which were converted to bigwigs using UCSC tool bedGraphToBigWig and visualized using IGV. Regions with positive PC1 values were identified as “A compartment” and regions with negative PC1 values were identified as “B compartment” where the genomic gene density regions were used to assign PC1 values to A and B compartments. To identify genomic regions that shift in compartmentalization between EF-Endo and EF-KD, and between EF-Rescue and EF-KD we used the findHiCCompartments.pl script in the Homer Hi-C analysis package to identify “A-like” and “B-like” shifting compartments. A-like compartment shifts were defined as regions changing in PC1 value (ΔPC1) by ≥ 0.4, while B-like compartment shifts were defined as regions changing in in PC1 value by ≤ -0.4 in EF-Endo vs. EF-KD and in EF-Rescue vs. EF-KD. The Juicebox visualization tool was used to coverage normalize Hi-C contact maps and generate Pearson correlation maps at 1mb resolution. ΔPC1 tracks were created by subtracting EF-KD PC1 bigwig from EF-Endo PC1 or EF-Rescue PC1 bigwig using deepTools.

### Integrative analysis of shifting compartments, EWS/FLI binding, H3K27Ac and gene expression

Genes mapping to stable and shifting compartments were annotated and overlapped with differential gene expression data from RNA-sequencing experiments using R to compare gene expression signatures for stable, A-like and B-like compartments in EF-Endo vs EF-KD and EF-Rescue vs. EF-KD. Similarly, differential H3K27Ac peaks were mapped to their respective stable, A-like and B-like compartments to compare H3K27Ac signature at stable and shifting compartments. Conserved EWS/FLI peaks mapping to shifting and stable compartments were identified using BEDTools’ intersect tool. Histogram analysis of PC1 values at all the conserved EWS/FLI binding sites as well as at only GGAA-μsats and non-μsats were performed using annotatePeaks.pl script in the Homer Hi-C analysis package. 20kb compartmental segments directly overlapping EWS/FLI peaks were identified using BEDTools’ intersect tool and were further separated into the type of EWS/FLI binding site (GGAA-μsat or non-μsat) to identify compartmental signatures of EWS/FLI binding locations genome wide.

### Detection of TADs and TAD boundaries

We identified TAD boundaries from our Hi-C data using the iHiC and TopDom packages (Hu, 2020; Shin et al., 2016). Intra-chromosomal **i**nteraction **i**ndex (III) files (see Hi-C data processing section above) were converted to interaction matrix files using *iHiC_III2MTX4TopDom* tool from iHiC package at 20kb resolution for compatibility with TopDom. The interaction matrix files were then run through the TopDom tool to identify TADs and TAD boundaries for each chromosome. TAD boundaries with p-value < 0.05 were extracted from .binSignal files using *iHiC_TADBoundary_TopDom* tool from iHiC package. Boundaries were overlapped using ChIPpeakAnno, GenomicRanges, VennDiagram, and Vennerable R packages. Boundaries common to all three EF-Endo, EF-KD and EF-Rescue conditions were termed stable boundaries, boundaries common to both the EWS/FLI containing conditions (EF-Endo and EF-Rescue) were termed EF-WT boundaries, and boundaries specific to EF-KD condition only were termed EF-KD boundaries.

### CTCF CUT&Tag peak calling and localization detection at TAD boundaries

CTCF CUT&Tag peaks were called using the following parameters for each experiment: A673 EF-Endo CTCF: FDR < 0.05, IDR < 0.01, mean counts ≥ 100, and log_2_FC ≥ 3; A673 EF-KD CTCF: FDR < 0.05, IDR < 0.01, mean counts ≥ 100, and log_2_FC ≥ 3; A673 EF-

Rescue CTCF: FDR < 0.05, IDR < 0.01, mean counts ≥ 200, and log_2_FC ≥ 3; TC71 EF-Endo CTCF: FDR 0.05, IDR < 0.01, mean counts ≥ 1300, and log_2_FC ≥ 3; TC71 EF-KD CTCF: FDR 0.05, IDR < 0.01, mean counts ≥ 1300, and log_2_FC ≥ 3. BEDTools’ intersect tool was used to identify CTCF localization in EF-Endo, EF-KD and EF-Rescue at stable, EF-WT and EF-KD boundaries. deepTools’ computeMatrix and plotHeatmap functions were used to generate heatmaps for CTCF at stable, EF-KD and EF-WT boundary regions for EF-Endo, EF-KD and EF-Rescue (Figure S4A).

### EWS/FLI enrichment at TAD boundaries

Conserved EWS/FLI peaks at stable, EF-WT and EF-KD boundaries were identified using BEDTools’ intersect tool. EWS/FLI log_2_FC enrichment over IgG was plotted to determine EWS/FLI enrichment at stable, EF-WT and EF-KD boundary regions.

### Directionality statistic calculation and generation of TAD level heatmaps

Publicly available Bioconductor diffHiC R package (Lun and Smyth, 2015) was used to partition the genome into 20kb bins. Using the domainDirections function, the total number of read pairs between each 20kb bin and a 200kb span upstream of that bin was calculated. The same analysis was repeated for a 200kb span downstream of the bin to yield two counts per 20kb bin (up and down). Low abundance bins were removed and the up and down counts were used to calculate a directionality statistic (DS) for each bin by computing the log fold change ratio between up and down counts for each bin in each condition (EF-Endo, EF-KD and EF-Rescue). The DS is similar to the directionality index defined by Dixon et al., 2012, where the magnitude of the DS identifies the size of interaction preference for a target bin, while the positive/negative signs provide information about upstream/downstream directionality preference of interaction for that target bin. The rotPlaid function in diffHiC was used to generate the Hi-C heatmaps in Figures 4 and S4.

### Detection of differential interactions

Differential interactions were detected for EF-Endo vs. EF-KD and EF-Rescue vs. EF-KD using diffHiC. The mapping location of each de-duplicated and map quality filtered read was matched to a restriction fragment in the hg19 reference genome and the resulting data were stored in index files using the HDF5 format. The HDF5 files were used to filter out reads with large (>700bp) and out of sync fragments. The reference genome was divided into either 1mb or 20kb contiguous bins and interaction intensity between any two bins (bin pair) in the genome were calculated by counting the number of read pairs that have one read successfully mapped to each of the two bins. Bin pairs with low abundance read counts were directly filtered to retain bin pairs with abundances that are at least 10- fold higher than the median abundance of inter-chromosomal interactions to account for non-specific ligations. The resulting data were TMM normalized (assumes that the intensities of most interactions do not change between experimental conditions) to reduce composition biases. Variability in read counts was modelled to reduce the significance of any differences in counts detected between biological replicates, and significance testing was performed to identify differential interactions at 20kb and 1mb resolution (FDR < 0.05 and |logFC| > 0.6) in EF-Endo vs. EF-KD and EF-Rescue vs. EF-KD using the R package edgeR (McCarthy et al., 2012; Robinson et al., 2010). Multidimensional scaling (MDS) plots of distances between Hi-C replicates were generated using the plotMDS function from the limma R package for the top 1000 interactions at 1mb resolution (Ritchie et al., 2015). Volcano plots of differential interactions were generated using EnhancedVolcano R package. 1mb loops were split into lost (FDR < 0.05 and logFC < -0.6) or gained loops (FDR < 0.05 and logFC > 0.6), inter-chromosomal loops were removed and the loop lengths were calculated by measuring the genomic distance between the midpoints of the two loop anchors spanning a loop.

### Integrative analysis of 20kb differential interactions

Homer motif finding tool findMotifsGenome.pl was used to determine enriched motifs associated with gained loop anchors at 20kb resolution. Significantly altered loops (FDR<0.05 and |logFC| >0.6) were mapped to genomic GGAA sequences of varying repeat lengths using BEDTools’ pairToBed tool. These differential interactions mapping to GGAA repeat sequences in the genome were further categorized by (i) the *total* number of genomic GGAA repeats (0-10, 11-20, 21-30, and >30 GGAA repeats) underlying the interactions (the maximum distance allowed between each GGAA motif is at most 20bp), and by (ii) the number of *consecutive* GGAA repeats (0-5, 6-10, 11-5, and 16-20 GGAA repeats) underlying the interactions (the maximum distance allowed between each GGAA motif is 0bp).

All 20kb diffHiC loops were mapped to conserved EWS/FLI sites using BEDTools’ pairToBed tool to identify loops that either overlap a conserved EWS/FLI peak or not. EWS/FLI bound loops were further categorized by the type of EWS/FLI binding site underlying the loop anchors to determine the role of EWS/FLI binding site in differential loop enrichment.

All 20kb diffHiC loops were mapped to differential enhancer regions using BEDTools’ pairToBed tool and were categorized by loops mapping to stable, gained or lost enhancers to determine the association between enhancers and loop enrichment. Gained and lost loops were annotated for genes in differential expression dataset for EF-Endo vs. EF-KD and for EF-Rescue vs. EF-KD in A673 cells using subsetByOverlaps function in GenomicRanges R package, differential expression of these genes were visualized by plotting volcano plots using the EnhancedVolcano R package.

### Identification of direct EWS/FLI target genes

Differentially expressed genes with TSS located within 20kb of a GGAA-μsat or non-μsat bound EWS/FLI conserved CUT&Tag peak were annotated using subsetByOverlaps function in R. The hg19.ensGene file containing all gene locations was downloaded from UCSC Genome Browser. The file was modified in R to identity only the starting location of all genes mapping to the hg19 reference genome. BEDTools’ intersect tool was used to identify hg19 genes with TSS mapping to a 20kb gained loop anchor (promoter proximal anchor). The merge.data.frame R base function was used to identify differentially expressed genes by merging hg19 genes mapped to 20kb gained loops to differential gene expression data from EF-Endo vs. EF-KD in A673 cells by Ensembl ID. Loop anchors paired with promoter proximal anchors (distal anchors) were identified using a unique loop identifier (LoopID). Promoter proximal and distal anchors were then annotated for the type of conserved EWS/FLI binding site underlying these loops using BEDTools’ intersect tool. Genes mapping to loops that do not overlap with an EWS/FLI binding site were removed and significantly regulated genes were identified using adjusted p-value < 0.05 and |log_2_FC|<0.3.

### Gene set enrichment analysis (GSEA)

GSEA (Version 4.0.3; Mootha et al., 2003; Subramanian et al., 2005) was used to analyze functional association between genes directly regulated by conserved EWS/FLI peaks in A673 cells and genes differentially regulated in TC71 cells upon EWS/FLI knock-down. Direct EWS/FLI regulated genes in EF-Endo vs. EF-KD A673 cells were used as gene sets for up- and down-regulated genes. Differentially expressed genes in EF-Endo vs. EF-KD TC71 cells were rank ordered by log_2_FC for comparison with the A673 EWS/FLI direct target gene sets. Significance was determined using |NES| > 1.5, FDR q-value < 0.05 and FWER p-value < 0.05.

### 4C experiment

4C sequencing libraries were prepared in duplicate following published protocol (Krijger et al., 2020). For each replicate, approximately 5 million EF-Endo, EF-KD and EF-Rescue A673 cells and EF-Endo and EF-KD TC71 cells were trypsinized and re-suspend in 2.5mL isolation buffer (10% FBS/PBS). 2.5mL of 4% fixation buffer (1mL 37% formaldehyde + 8.25 mL isolation buffer) was added and cells were cross-linked for 10 minutes. Cold glycine was added to a final concentration of 0.125M to quench the fixation reaction and cells were centrifuged at 1000g for 5 mins at 4°C. Cells were washed with ice cold PBS, pelleted, flash frozen and stored at -80°C. Thawed pellets were gently re-suspended in 1 mL lysis buffer (50mM Tris-HCl pH 7.5, 0.5% Igepal, 1% TX-100, 150mM NaCl, 5mM EDTA) with 1× cOmplete protease inhibitors (Roche 11697498001), incubated on ice for 20 minutes, spun down, supernatant removed, and the nuclei washed in 450 μl 1.2× RE1 buffer. Nuclei were re-suspended in 500 μl 1.2× RE1 buffer, 10% SDS was added to a final volume of 0.3%, and incubated while shaking at 750 rpm for 1 hour at 37°C. 20% Triton-X 100 was added to a final volume of 2.5% and incubated in a thermomixer at 750 rpm for 1 hour at 37°C. 100U of MboI (primary restriction enzyme) was added and incubated in a thermomixer at 750 rpm for 3 hours at 37°C. A second round of 100U of MboI was added and incubated in a thermomixer at 750 rpm overnight at 37°C. MboI was heat inactivated at 65°C for 20 min. Proximity ligation was performed by incubating the samples for 24 hours at 16°C using 50U of T4 DNA ligase in 7mL of 1X ligation buffer (660mM Tris-HCl, pH 7.5, 50mM MgCl2, 50mM DTT, and 10mM ATP). 30 μL Proteinase K (10 mg/ml) was added and cross-links were reversed by incubating overnight at 65°C to create 3C template DNA. The 3C template was purified using Nucleomag PCR beads (“P-beads”, Macherey-Nagel #744100.1), eluted in 450 μl 5mM Tris-HCl (pH 8.0), and stored at -20°C. Second restriction digest was performed in 500uL of 1X RE2 buffer at 37°C overnight in a thermomixer at 500rpm using 50U of NlaIII. NlaIII was heat inactivated at 65°C for 20 minutes. DNA was quantified using Qubit and a second ligation reaction was carried out in 5mL 1X ligation buffer with 25 μg of template and 50 U T4 DNA ligase at 16°C overnight. Samples were purified using P-beads as above and stored at -20°C. PCR amplification was performed in 50μL PCR reactions using 200ng of 4C template, 1μL of dNTPS (1mM), 5μL of 5uM reading primer (5’-TACACGACGCTCTTCCGATCTGACTCGTCCCAAACTCTTAGCCTC), 5μL of 5uM non reading primer (5’- ACTGGAGTTCAGACGTGTGCTCTTCCGATCTCTAGGGCAGACAGATAACAG) and 0.7μL of Expand Long Template polymerase (Roche #11759060001). PCR cycling program used: 94°C for 2 minutes; 16 cycles: 94°C for 10 seconds, 50°C for 1 minute, 68°C for 3 minutes; 68°C for 5 minutes. Libraries were purified using 0.8X AMPure XP purification and a second round of PCR was performed in 50μL of 1X PCR buffer with 1μL of 10mM dNTPs, 5μL of the purified product, 5μL of 5μM universal primer mix (forward primer contains the illumina P5 adapter end that binds to the flowcell in the sequencer, and the reverse primer contains a 6 nucleotide Truseq index sequence), 0.7μL of Roche Expand Long Template polymerase mix. PCR cycling program used for second PCR: 94°C for 2 minutes; 20 cycles: 94°C for 10 seconds, 60°C for 1 minute, 68°C for 3 minutes; 68°C for 5 minutes. PCR amplified 4C libraries were purified using Qiagen PCR purification kit and primer dimers removed using 0.8X Ampure XP bead purification. 4C libraries were run on the High Sensitivity D5000 tape for library quantitation and were sequenced with Illumina MiSeq (A673 samples) and MiniSeq (A673 and TC71 samples) sequencing systems using 2X 75bp PE runs. Only the read containing the reading primer was used for further analysis. Reading primer was used to target the GGAA-μsat at ∼60kb upstream of NKX2-2 as the region of interest (viewpoint).

### 4C data processing and analysis

4C sequencing data was processed using the pipe4C analysis pipeline. Global parameters were set using the global configuration file (conf.yml) and experiment specific parameters were set using the viewpoint file (vPFile). The pipe4C pipeline was then run using the command: Rscript pipe4C.R --vpFile=./fastq/VPinfo.txt --fqFolder=./fastq/ -- outFolder=./outF_A673_TC71_NKX2-2_4Cseq/ --cores 12 --plot –wig. The pipe4C pipeline demultiplexes the reads to retain only reads containing the reading primer. Trimming of reads was performed to extract the capture sequence between the two restriction sites. Reads were then mapped to the hg19 reference genome using Bowtie2, and unmapped reads were removed. The pipeline counts reads per fragment end and creates an *in silico* fragment end library using the Bioconductor BSgenome.Hsapiens.UCSC.hg19 R package (Team, 2020b). The fragments ends were trimmed based on capture length and only reads that mapped uniquely to the reference genome with the capture length were counted. The read counts were normalized for sequencing depth to a total sum of 1 million mapped reads per sample. The mapped data were then smoothed by averaging using a running mean of 21 fragment ends to generate RDS files. The RDS file were used to identify regions that have higher contact frequency than expected using peakC. peakC models the background contact frequency for regions upstream and downstream of the viewpoint and identifies genomic regions that are significantly contacted by computing non-parametric statistics based on ranks of coverage of 4C fragment ends with respect to the background model.

### Statistics

Significance of experimental results were carried out using unpaired t-test for comparing two groups or one-way ANOVA (with multiple comparisons) for comparing three or more groups as appropriate. Significance was determined as a p < 0.05. These statistical tests were performed using GraphPad Prism 9. For GSEA, significance was determined using a normalized enrichment score (NES), FDR q-value and FWER p-value. A result was significant if |NES| > 1.5, FWER p-value < 0.05 and FDR q-value < 0.05. Adjusted p-values (padj) for differentially expressed genes in RNA-seq were generated by DESeq2 and padj <0.05 and |log_2_FC| > 0.3 was considered significant.

## ACKNOWLEDGEMENTS

We thank A. Snedden and Y. Zhang for their support with high performance computing; Institute for Genomic Medicine at Nationwide Children’s Hospital for providing sequencing and bioinformatics support; B. Stanton, B. Sunkel, J. Tokarsky, and A. Byrum for helpful comments and discussion. Research in this publication was supported by the National Institute of Health (NIH) award U54 CA231641 to SLL. The content is solely the responsibility of the authors and does not necessarily represent the official views of the NIH.

## AUTHOR CONTRIBUTIONS

Conceptualization, IAS and SLL; Methodology, IAS and SLL; Software, IAS, CT and ERT; Formal Analysis, IAS; Investigation, IAS, JS, ERT, MAB and JCC; Writing-Original Draft, IAS; Writing-Review & Editing, IAS, JS, MAB, ERT and SLL; Visualization, IAS; Project Administration, IAS; Funding Acquisition, SLL; Supervision, SLL.

## DECLARATION OF INTERESTS

SLL declares a competing interest as a member of the advisory board for Salarius Pharmaceuticals. SLL is also a listed inventor on United States Patent No. US 7,939,253 B2, “Methods and compositions for the diagnosis and treatment of Ewing’s sarcoma,” and United States Patent No. US 8,557,532, “Diagnosis and treatment of drug-resistant Ewing’s sarcoma.”

## DATA AVAILABILITY

All RNA-seq, CUT&Tag, 4C and *in situ* Hi-C data have been deposited at GEO. Accession numbers will be made public upon publication of the manuscript. All software used in this study are published and publicly available.

