## Supplemental data for "EWS/FLI mediated reprogramming of 3D chromatin promotes an altered transcriptional state in Ewing sarcoma"

Supplementary Figure S1

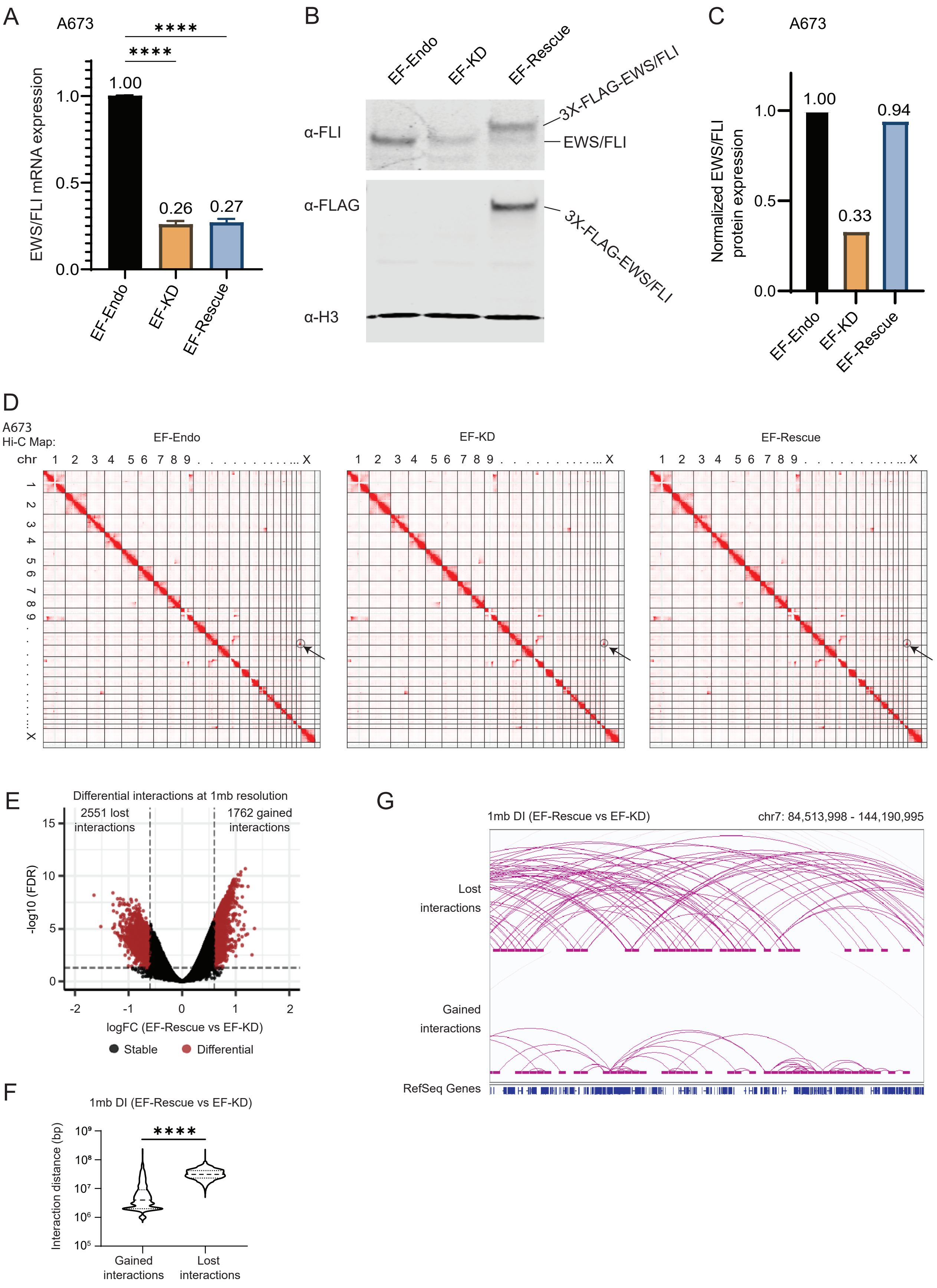

### Supplementary Figure S2

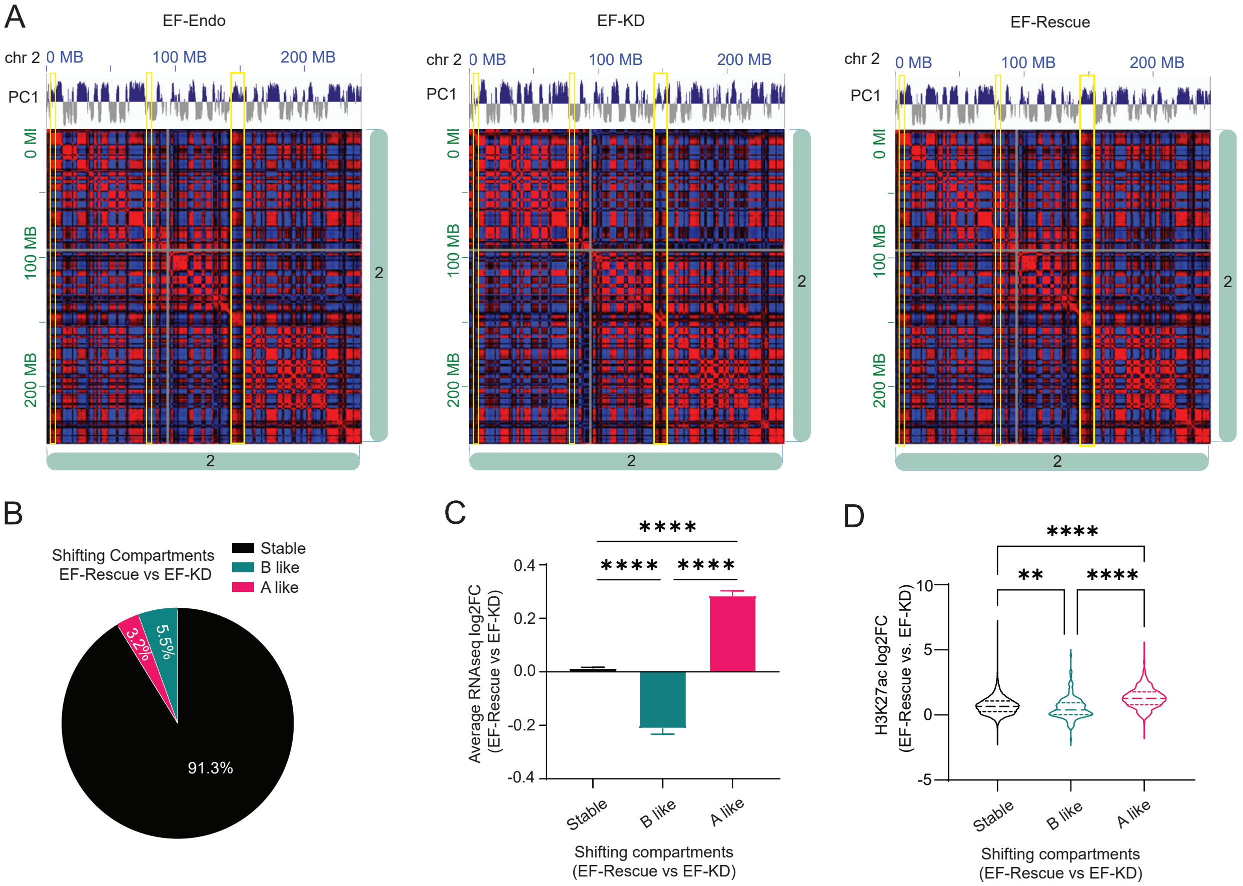

### Supplementary Figure S3

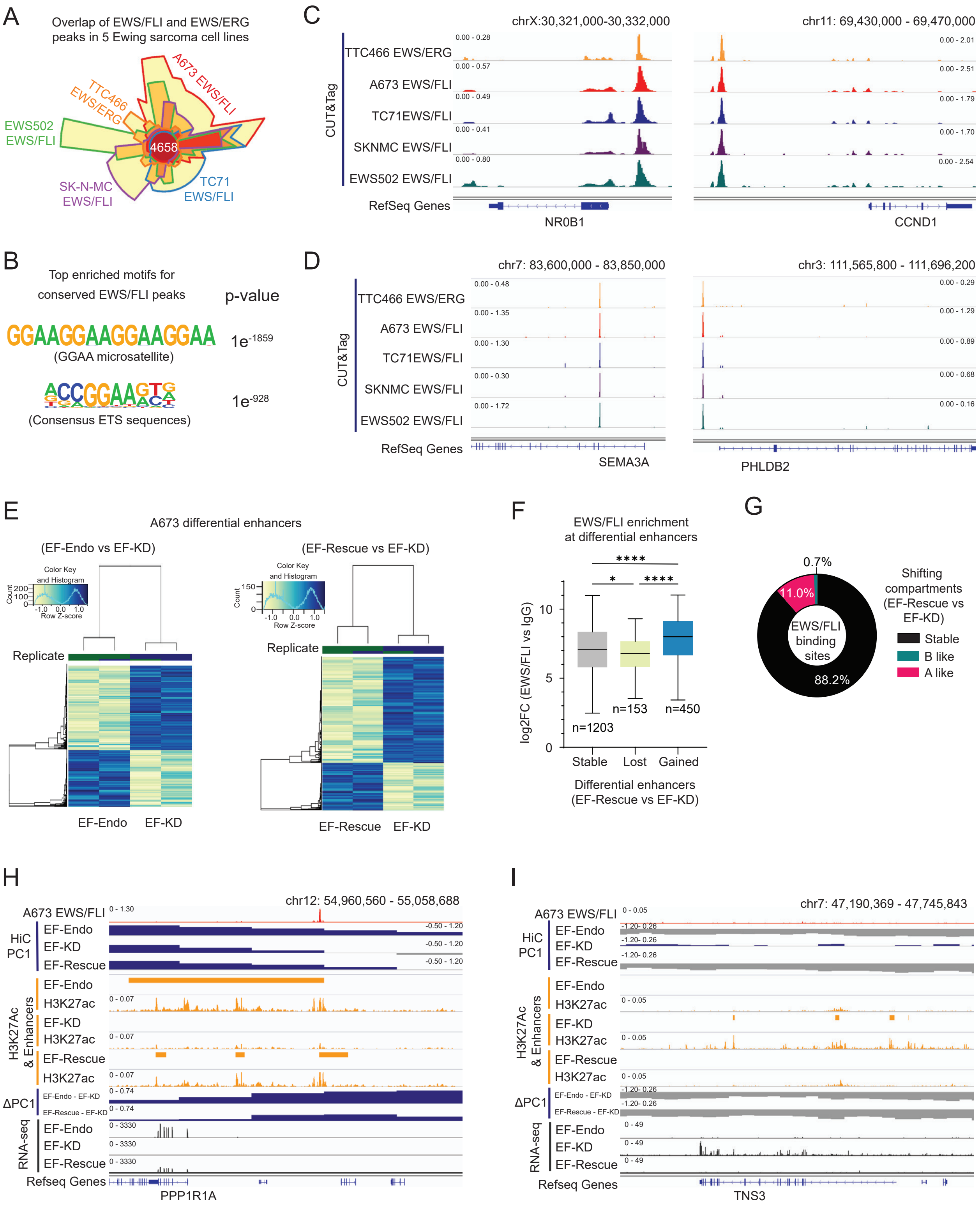

Supplementary Figure S4

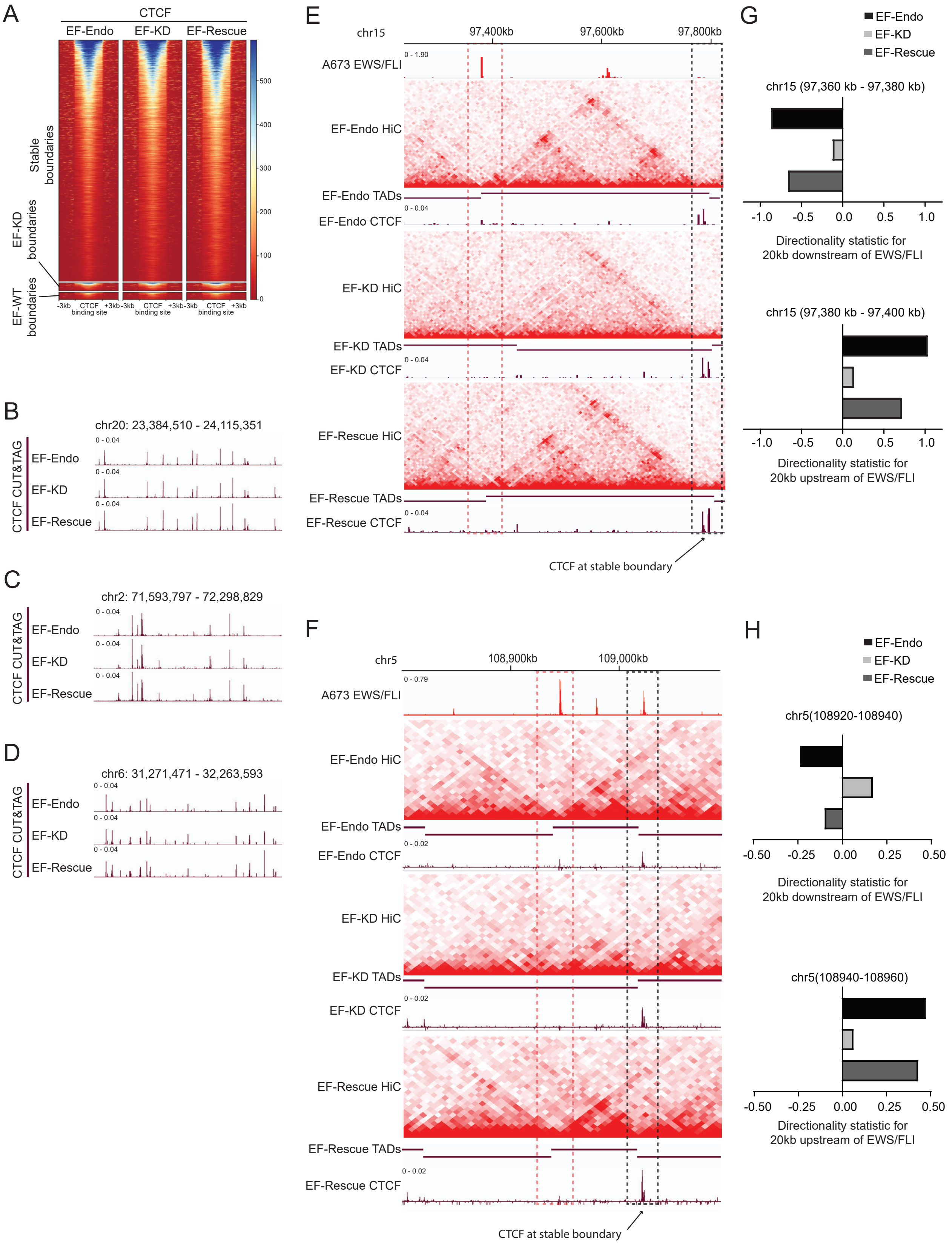

#### Supplementary Figure S5

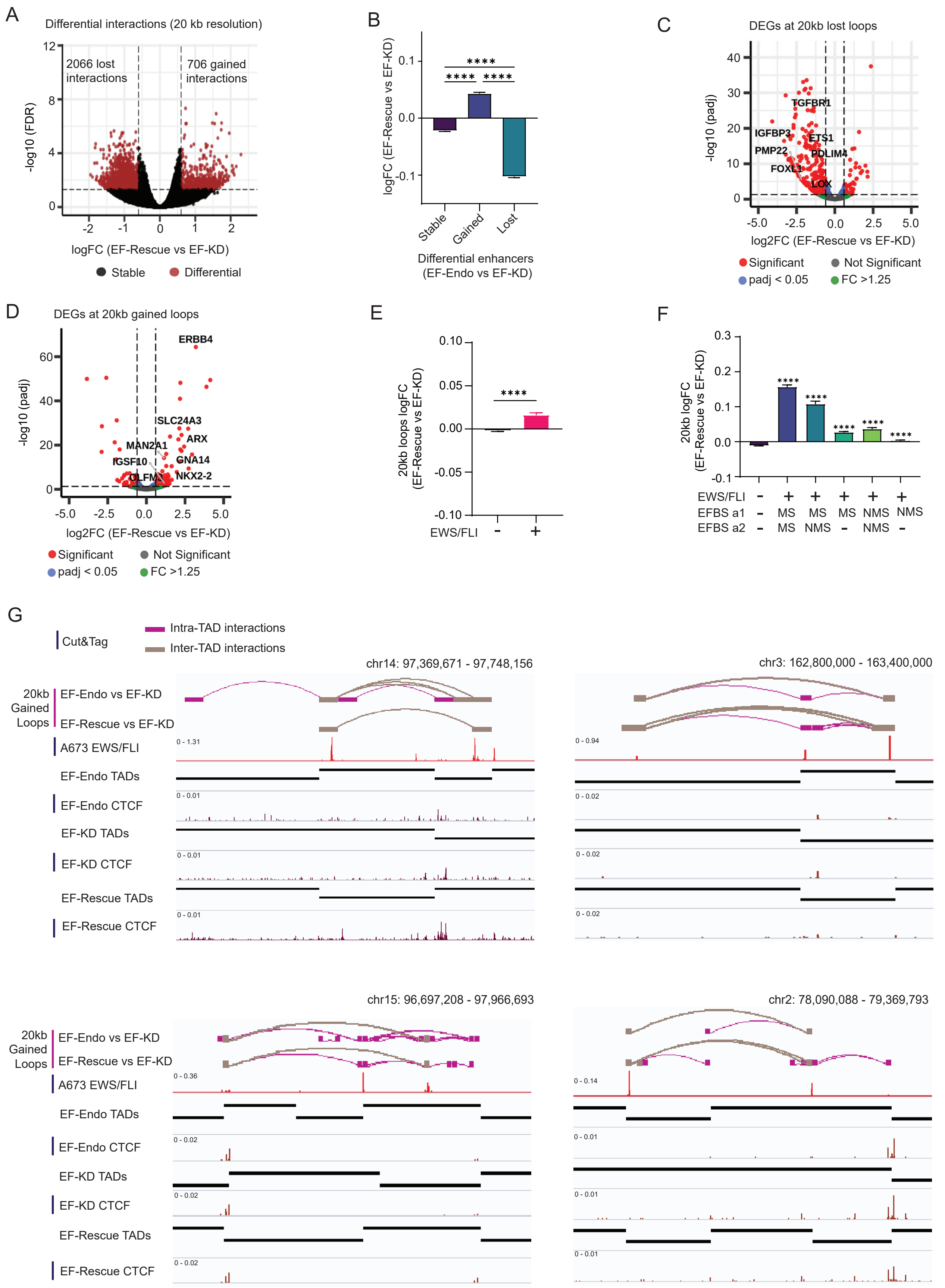

Supplementary Figure S6

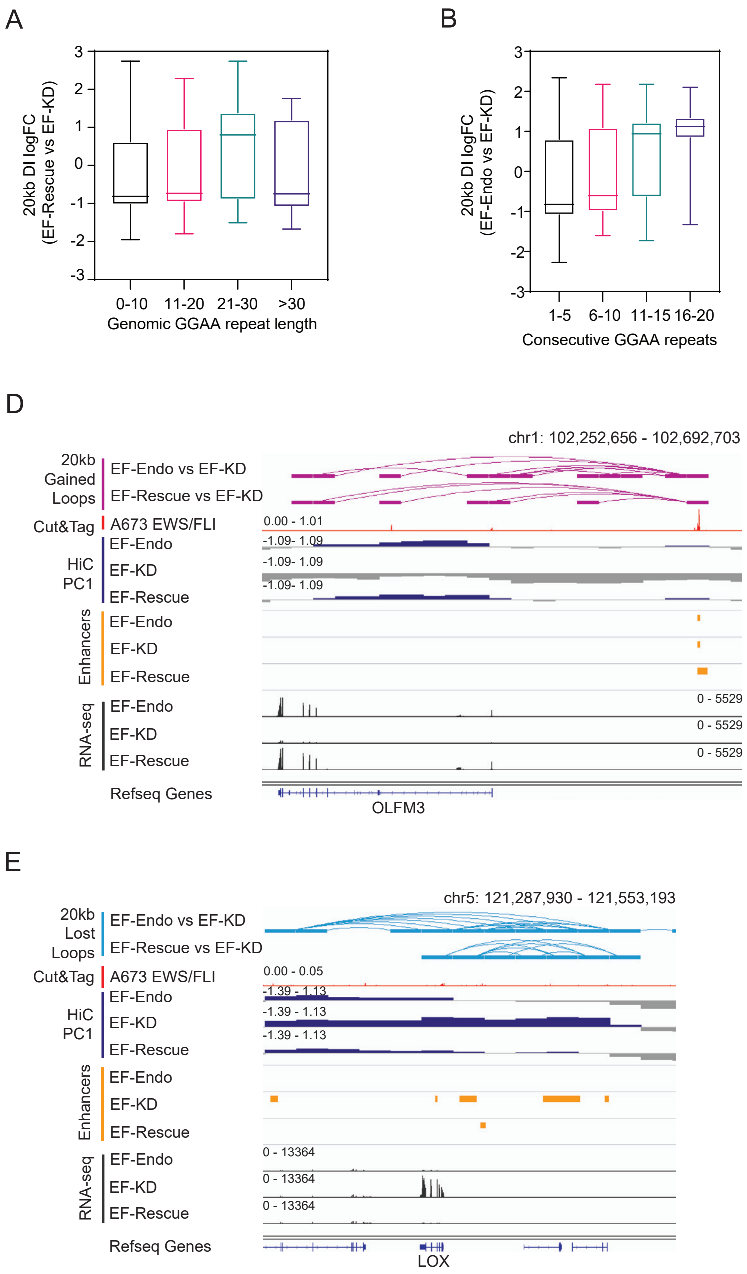

Supplementary Figure S7

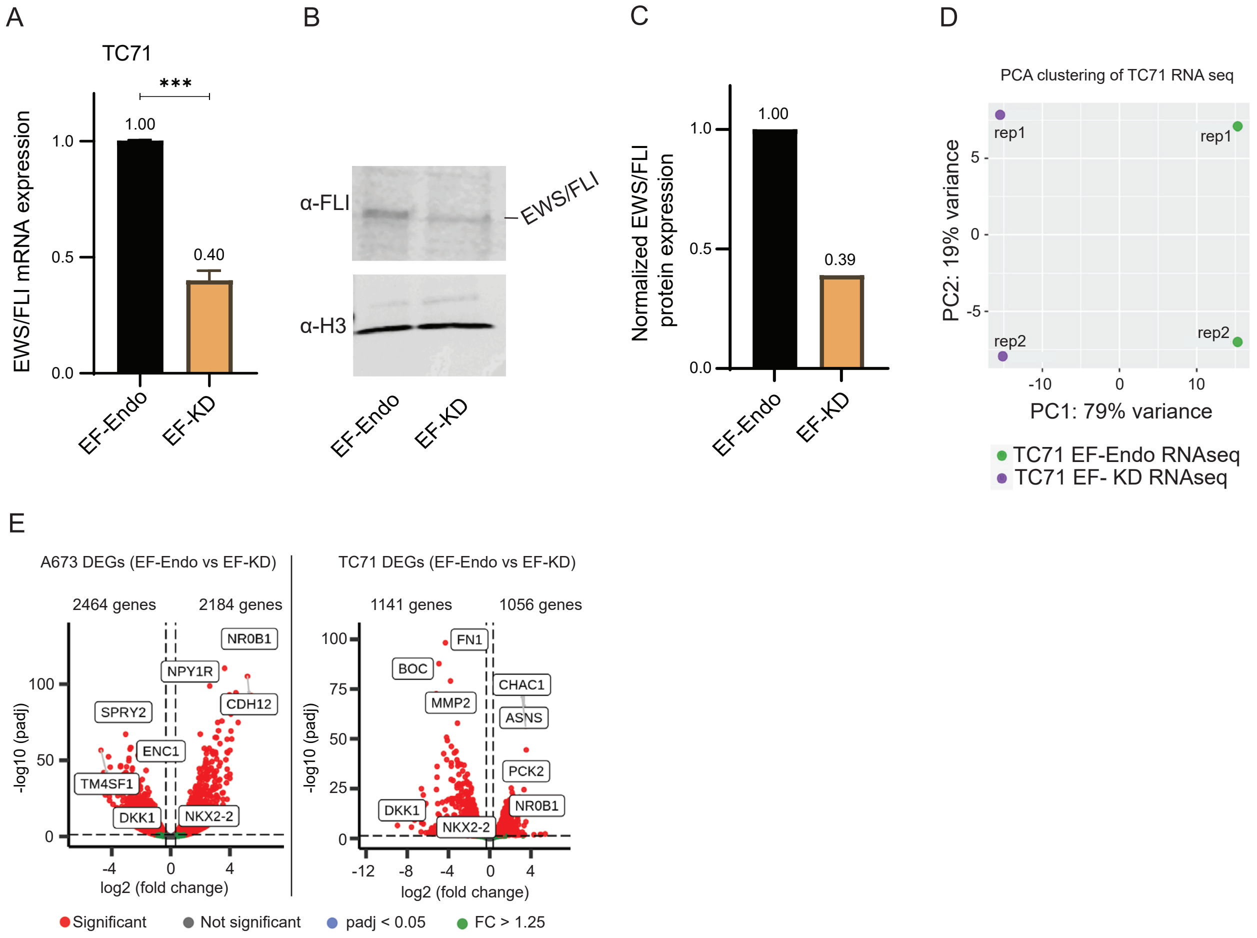

**Figure S1. EWS/FLI reprograms the global interaction profile in Ewing sarcoma**

(A) qRT-PCR validation of EWS/FLI knock-down/rescue in A673 Ewing sarcoma cells. PCR primers are specific to endogenous EWS/FLI transcript. n = 7 biological replicates. Mean ± SEM shown. **** P value<0.0001 (Dunnett's multiple comparisons test).

(B) Representative western blots for EWS/FLI in EF-Endo, EF-KD and EF-Rescue in A673 Ewing sarcoma cells. Nuclear lysates were probed with α-FLI, α-FLAG, or α-H3 antibodies. FLI and FLAG immunoblots were performed on separate membranes, both gels were loaded from the same sample tube and run simultaneously. FLAG and H3 were probed at the same time.

(C) Densitometry plot of the western blot in Figure S1B. EWS/FLI intensity was normalized against corresponding H3 band intensity.

(D) Genome-wide Hi-C maps for EF-Endo, EF-KD and EF-Rescue in A673 cells (n = 2). High to low interaction frequency (red to white). Arrows indicate the t(11;22) EWS/FLI translocation.

(E) Volcano plot of DIs at 1mb resolution (EF-Rescue vs. EF-KD). DI, FDR <0.05 & |logFC| > 0.6.

(F) Violin plots of interaction distance (bp) for DIs (EF-Rescue vs. EF-KD) at 1 mb resolution. **** P value < 0.0001 (Unpaired t-test).

(G) Example of lost and gained interactions at 1 mb resolution (EF-Rescue vs. EF-KD).

**Figure S2. Alterations in compartment structure associate highly with enhancer landscape and gene expression changes**

(A) Pearson correlation maps for intra-chromosomal interactions in chromosome 2 for EF-Endo, EF-KD and EF-Rescue. Loci with similar interaction partners are highly correlated (red). Loci with dissimilar interaction partners are anti-correlated (blue). Correlation ranges from -1 (blue) to +1 (red). Yellow boxes highlight alteration to compartmental pattern in EF-KD compared to EF-Endo and EF-Rescue.

(B) Pie chart showing percentage of the genome undergoing compartment shifting (EF-Rescue vs. EF-KD). ΔPC1 = EF-Rescue PC1 – EF-KD PC1. ΔPC1 ≥ 0.4 (A like), ΔPC1≤ -0.4 (B like), 0.4 > ΔPC1 > -0.4 (stable).

(C) Average expression of genes in EF-Rescue vs. EF-KD annotating to a shifting compartment. Mean ± SEM shown. **** P value < 0.0001 (Šídák's multiple comparisons test).

(D) Violin plots showing $log_2$FC enrichment of H3K27Ac in EF-Rescue vs. EF-KD at shifting compartments. ****P value < 0.0001, **P value < 0.01 (Tukey's multiple comparisons test).

**Figure S3. EWS/FLI promotes active compartmentalization of chromatin**

(A) Overlap of EWS/FLI and EWS/ERG genomic localization in five Ewing sarcoma cell lines to scale. 4658 overlapping regions were found to be common in all 5 cell lines, identified in red (center). Peaks within 200bp of each other were considered overlapping.

(B) Top enriched motifs for conserved EWS/FLI peaks in A673, identified using Homer.

(C, D) Representative examples of conserved EWS/FLI and EWS/ERG localization in all five cell lines at (C) GGAA-μsats and (D) non-μsats.

(E) Heatmaps showing differential enhancers (including super-enhancers) in EF-Endo vs EF-KD and in EF-Rescue vs. EF-KD. Enhancer regions showing significant alteration to H3K27Ac levels were identified as differential enhancers, FDR < 0.05.

(F) Boxplots showing EWS/FLI enrichment in A673 cells at differential enhancers (EF-Rescue vs. EF-KD). ****P value < 0.0001 (Tukey's multiple comparisons test).

(G) Doughnut chart showing proportion of EWS/FLI occupancy at shifting compartments (EF-Rescue vs. EF-KD).

(H, I) IGV traces showing EWS/FLI binding in A673, and corresponding compartment profiles (PC1), enhancers, and gene expression (RNA-seq) in EF-Endo, EF-KD and EF-Rescue. (H) Compartment activation & (I) Compartment inactivation in EF-Endo and EF-Rescue compared to EF-KD.

**Figure S4. EWS/FLI perturbs TAD boundaries**

(A) Heatmaps showing CTCF CUT&Tag signal at TAD boundaries (stable, EF-WT, and EF–KD boundaries). CTCF binding sites were clustered by respective boundary regions.

(B, C, D) Representative examples of conserved CTCF chromatin localization in EF-Endo, EF-KD, and EF-Rescue cells.

(E, F) Examples of EF-WT (dashed orange box) and stable boundaries (dashed black box) over a (E) 600 kb region in chromosome 15 and (G) 300 kb region in chromosome 5 as shown in Hi-C heatmaps and TAD calls, with corresponding EWS/FLI and CTCF localization.

(G, H) Directionality statistic (DS) calculated for 20kb regions upstream (top panel) and downstream (bottom panel) of EWS/FLI binding site (orange dashed box) in EF-Endo, EF-KD, and EF-Rescue.

**Figure S5. EWS/FLI mediates chromatin looping**

(A) Volcano plot showing differential (DL, red) and stable (gray) loops (EF-Rescue vs. EF-KD) at 20 kb resolution. DL, (FDR<0.05, |logFC| > 0.6).

(B) Average logFC enrichment of 20kb loops at differential enhancers (EF-Rescue vs. EF-KD). Mean ± SEM shown. **** P value < 0.0001 (Tukey's multiple comparisons test).

(C) & (D) Volcano plots showing differentially expressed genes (EF-Rescue vs. EF-KD) mapping to (C) lost and (D) gained loops at 20kb resolution (EF-Rescue vs. EF-KD). Differentially expressed genes (in red), adjusted p-value < 0.05 and $|\log_2 FC| > 0.3$.

(E) Average logFC enrichment of 20kb loops in EF-Rescue vs. EF-KD associated with a EWS/FLI binding site (+) or no EWS/FLI binding site (-). Mean ± SEM shown. **** P value < 0.0001 (Unpaired t-test).

(F) Average logFC (EF-Rescue vs. EF-KD) enrichment of 20kb loops associated with the type of EWS/FLI binding site at each loop anchor. Mean ± SEM shown. **** P value < 0.0001 (Dunnett's multiple comparisons test). All comparisons made to the EWS/FLI (-) loops. See Figure 5.

(G) Examples of EWS/FLI mediated inter- TAD (tan) and intra-TAD (purple) gained loops at 20kb resolution. Corresponding TAD calls (black bars), EWS/FLI localization and CTCF localization are also shown. Inter-TAD loops were identified as loops with each loop anchor mapping to a separate TAD.

**Figure S6. EWS/FLI mediates chromatin looping**

(A) Boxplots showing enrichment of 20kb DLs (EF-Rescue vs. EF-KD) mapping to *total* number of genomic GGAA motifs (sequence between two GGAA motifs ≤ 20-bp).

(B, C) Boxplots showing enrichment of 20kb DLs mapping to *consecutive* number of GGAA motifs (sequence between two GGAA motifs =0bp), independent of EWS/FLI binding. (B) EF-Endo vs. EF-KD, (C) EF-Rescue vs. EF-KD.

(D, E) Examples of 20kb DLs and corresponding EWS/FLI localization, enhancers, compartments (PC1), and gene expression (RNA-seq). (D) Gained loops, (E) Lost loops.

**Figure S7. Cell-specific local chromatin structure affects gene regulation**

(A) qRT-PCR showing depletion of EWS/FLI following knock-down with shRNA targeting EWS/FLI transcript (EF-KD) or control shRNA targeting Luciferase transcript (EF-Endo) in TC71 cells. PCR primers are specific to endogenous EWS/FLI transcript. 5 biological replicates were used. Mean ± SEM shown. *** P value < 0.001 (Welch's t-test).

(B) Representative western blot for EWS/FLI in EF-Endo and EF-KD. Nuclear lysates were probed with α-FLI and α-H3 antibodies. FLI and H3 immunoblots were performed on separate membranes, both gels were loaded at the same volume from the same sample tubes and run simultaneously.

(C) Densitometry plot of the western blot in Figure S8B. EWS/FLI intensity was normalized against corresponding H3 band intensity.

(D) Principal component analysis. PC2 vs PC1 plotted for RNA sequencing replicates of EF-Endo vs EF-KD TC71 cells.

(E) Volcano plots of differentially expressed genes (DEGs, red) in EF-Endo vs EF-KD in A673 (left) and TC71 (right). Significant, padj < 0.05 & |fold change| > 1.25.

**Supplementary table S1. Genes directly upregulated by EWS/FLI in A673**

| Ensembl ID | Gene symbol | 20kb gained loop | EWS/FLI binding site | | | | p-adj (EF-Endo vs EF-KD) |
|---|---|---|---|---|---|---|---|
| | | | Proximal | | Distal | | |
| | | | MS | NMS | MS | NMS | |
| ENSG00000169297 | NR0B1 | | ✓ | | | | 2.2133E-129 |
| ENSG00000154162 | CDH12 | ✓ | ✓ | | | | 9.46E-106 |
| ENSG00000168671 | UGT3A2 | ✓ | ✓ | ✓ | | | 1.93E-99 |
| ENSG00000234224 | TMEM229A | | ✓ | ✓ | | | 5.09217E-95 |
| ENSG00000251460 | RP11-1258F18.1 | ✓ | ✓ | | | | 1.18E-93 |
| ENSG00000165566 | AMER2 | ✓ | ✓ | | | | 4.58E-81 |
| ENSG00000166501 | PRKCB | ✓ | ✓ | ✓ | ✓ | | 1.91E-76 |
| ENSG00000135447 | PPP1R1A | ✓ | | ✓ | | | 1.74E-75 |
| ENSG00000144644 | GADL1 | | ✓ | | | | 3.929E-70 |
| ENSG00000230316 | FEZF1-AS1 | ✓ | ✓ | ✓ | | | 6.63E-70 |
| ENSG00000256553 | TRAV1-2 | | ✓ | ✓ | | | 1.0962E-65 |
| ENSG00000188992 | LIPI | | | ✓ | | | 1.30073E-65 |
| ENSG00000128610 | FEZF1 | ✓ | ✓ | ✓ | | | 1.49E-64 |
| ENSG00000134709 | HOOK1 | | ✓ | ✓ | | | 1.6142E-58 |
| ENSG00000043591 | ADRB1 | ✓ | | ✓ | ✓ | | 1.69E-58 |
| ENSG00000164647 | STEAP1 | | | ✓ | | | 1.42799E-57 |
| ENSG00000140015 | KCNH5 | ✓ | | | ✓ | | 2.46E-57 |
| ENSG00000146038 | DCDC2 | | | ✓ | | | 1.28033E-56 |
| ENSG00000157214 | STEAP2 | | ✓ | ✓ | | | 9.87659E-56 |
| ENSG00000183780 | SLC35F3 | | | ✓ | | | 1.1023E-55 |
| ENSG00000154359 | LONRF1 | ✓ | | ✓ | ✓ | | 3.88E-55 |
| ENSG00000182050 | MGAT4C | ✓ | ✓ | ✓ | | | 6.1E-55 |
| ENSG00000104870 | FCGRT | | ✓ | ✓ | | | 1.17362E-54 |
| ENSG00000196083 | IL1RAP | | ✓ | ✓ | | | 2.70373E-51 |
| ENSG00000147869 | CER1 | | ✓ | ✓ | | | 7.49398E-50 |
| ENSG00000150672 | DLG2 | | ✓ | ✓ | | | 2.0195E-49 |
| ENSG00000163406 | SLC15A2 | ✓ | ✓ | | | | 3.82E-49 |
| ENSG00000188778 | ADRB3 | ✓ | ✓ | | | ✓ | 5.53118E-47 |
| ENSG00000108176 | DNAJC12 | ✓ | ✓ | ✓ | | ✓ | 2.08226E-46 |
| ENSG00000156564 | LRFN2 | ✓ | ✓ | ✓ | | ✓ | 9.64562E-46 |
| ENSG00000240694 | PNMA2 | ✓ | | ✓ | | ✓ | 3.49658E-45 |
| ENSG00000118507 | AKAP7 | ✓ | ✓ | ✓ | ✓ | ✓ | 3.5E-45 |
| ENSG00000213626 | LBH | ✓ | ✓ | ✓ | | ✓ | 8.44938E-45 |
| ENSG00000102524 | TNFSF13B | ✓ | | ✓ | | ✓ | 1.41E-43 |
| ENSG00000146409 | SLC18B1 | ✓ | ✓ | | | ✓ | 5.39122E-43 |
| ENSG00000081803 | CADPS2 | ✓ | | ✓ | | ✓ | 1.69989E-42 |
| ENSG00000165071 | TMEM71 | ✓ | ✓ | | | ✓ | 2.4754E-42 |
| ENSG00000161681 | SHANK1 | ✓ | ✓ | | | ✓ | 9.94687E-42 |
| ENSG00000185272 | RBM11 | ✓ | | ✓ | | ✓ | 1.26876E-41 |
| ENSG00000182601 | HS3ST4 | ✓ | | ✓ | | ✓ | 8.73383E-41 |
| ENSG00000168421 | RHOH | ✓ | ✓ | | | ✓ | 1.27102E-40 |
| ENSG00000076864 | RAP1GAP | ✓ | ✓ | | | ✓ | 4.79445E-40 |

| | | | | | | | |
|---|---|---|---|---|---|---|---|
| ENSG00000036448 | MYOM2 | ✓ | | ✓ | | ✓ | 6.69E-40 |
| ENSG00000017427 | IGF1 | ✓ | ✓ | ✓ | | ✓ | 7.12184E-40 |
| ENSG00000168672 | FAM84B | ✓ | ✓ | ✓ | ✓ | ✓ | 2.22E-39 |
| ENSG00000163347 | CLDN1 | ✓ | | ✓ | | ✓ | 2.54699E-39 |
| ENSG00000181449 | SOX2 | ✓ | | ✓ | | ✓ | 4.92423E-39 |
| ENSG00000211777 | TRAV3 | ✓ | | | ✓ | ✓ | 6.77857E-39 |
| ENSG00000185052 | SLC24A3 | ✓ | ✓ | ✓ | | ✓ | 5.64E-38 |
| ENSG00000141968 | VAV1 | ✓ | | ✓ | | ✓ | 1.60241E-37 |
| ENSG00000187094 | CCK | ✓ | ✓ | | | ✓ | 1.02183E-36 |
| ENSG00000248329 | APELA | ✓ | ✓ | | | ✓ | 6.73E-36 |
| ENSG00000184613 | NELL2 | ✓ | ✓ | ✓ | | ✓ | 1.74E-34 |
| ENSG00000124171 | PARD6B | ✓ | | ✓ | | ✓ | 9.7188E-34 |
| ENSG00000157240 | FZD1 | ✓ | | ✓ | | ✓ | 2.54107E-33 |
| ENSG00000170579 | DLGAP1 | ✓ | ✓ | ✓ | ✓ | ✓ | 3.42E-33 |
| ENSG00000171766 | GATM | ✓ | | ✓ | | ✓ | 1.30105E-32 |
| ENSG00000110092 | CCND1 | ✓ | ✓ | ✓ | ✓ | ✓ | 1.65E-32 |
| ENSG00000198848 | CES1 | ✓ | ✓ | | | ✓ | 4.81908E-32 |
| ENSG00000253888 | RP11-521M14.1 | ✓ | ✓ | ✓ | ✓ | ✓ | 1.26E-31 |
| ENSG00000133878 | DUSP26 | ✓ | ✓ | ✓ | | ✓ | 2.98412E-31 |
| ENSG00000116774 | OLFML3 | ✓ | | ✓ | | ✓ | 5.21997E-30 |
| ENSG00000250305 | KIAA1456 | ✓ | ✓ | ✓ | | ✓ | 5.35171E-30 |
| ENSG00000164418 | GRIK2 | ✓ | ✓ | ✓ | | ✓ | 9.58E-30 |
| ENSG00000231532 | LINC01249 | ✓ | | ✓ | | ✓ | 1.32268E-29 |
| ENSG00000079101 | CLUL1 | ✓ | ✓ | ✓ | | ✓ | 3.13E-29 |
| ENSG00000159023 | EPB41 | ✓ | | ✓ | | ✓ | 3.2884E-29 |
| ENSG00000102445 | RUBCNL | ✓ | ✓ | | | ✓ | 6.18E-29 |
| ENSG00000168765 | GSTM4 | ✓ | | ✓ | | ✓ | 7.43554E-29 |
| ENSG00000138030 | KHK | ✓ | | | ✓ | ✓ | 1.58138E-28 |
| ENSG00000182752 | PAPPA | ✓ | ✓ | | ✓ | ✓ | 1.69E-28 |
| ENSG00000118420 | UBE3D | | | ✓ | | | 1.83727E-28 |
| ENSG00000107736 | CDH23 | | ✓ | | | | 2.96714E-28 |
| ENSG00000105642 | KCNN1 | | ✓ | | | | 6.25568E-28 |
| ENSG00000228354 | RP11-552J9.9 | | ✓ | | | | 5.75917E-27 |
| ENSG00000140600 | SH3GL3 | ✓ | ✓ | | | | 7.93E-27 |
| ENSG00000169618 | PROKR1 | ✓ | | ✓ | | | 8.87E-27 |
| ENSG00000144036 | EXOC6B | | ✓ | ✓ | | | 1.77938E-26 |
| ENSG00000184368 | MAP7D2 | | ✓ | | | | 2.42946E-26 |
| ENSG00000130529 | TRPM4 | | ✓ | | | | 2.93322E-26 |
| ENSG00000165588 | OTX2 | ✓ | ✓ | | | | 4.09E-26 |
| ENSG00000176595 | KBTBD11 | ✓ | | ✓ | | | 4.7E-26 |
| ENSG00000167210 | LOXHD1 | ✓ | ✓ | ✓ | | | 3.56E-25 |
| ENSG00000009709 | PAX7 | | ✓ | ✓ | | | 3.77262E-25 |
| ENSG00000145247 | OCIAD2 | | ✓ | | | | 1.55157E-24 |
| ENSG00000078618 | NRDC | | ✓ | ✓ | | | 1.67262E-24 |
| ENSG00000183250 | LINC01547 | | ✓ | ✓ | | | 3.24075E-24 |
| ENSG00000105472 | CLEC11A | | ✓ | | | | 8.67519E-24 |

| | | | | | | | |
|---|---|---|---|---|---|---|---|
| ENSG00000175538 | KCNE3 | | ✓ | ✓ | | | 1.20964E-23 |
| ENSG00000169744 | LDB2 | | ✓ | ✓ | | | 1.21199E-23 |
| ENSG00000064835 | POU1F1 | ✓ | | ✓ | ✓ | | 2.08E-23 |
| ENSG00000058866 | DGKG | ✓ | ✓ | | ✓ | | 5.03E-23 |
| ENSG00000123977 | DAW1 | ✓ | | | ✓ | | 5.03467E-23 |
| ENSG00000072415 | MPP5 | | | ✓ | | | 8.72539E-23 |
| ENSG00000182957 | SPATA13 | | ✓ | | | | 1.02989E-22 |
| ENSG00000185668 | POU3F1 | | ✓ | ✓ | | | 1.66198E-22 |
| ENSG00000227932 | RP13-16H11.9 | | ✓ | | | | 2.32871E-22 |
| ENSG00000196730 | DAPK1 | | ✓ | ✓ | | | 2.81129E-22 |
| ENSG00000050628 | PTGER3 | | | ✓ | | | 4.73081E-22 |
| ENSG00000099284 | H2AFY2 | | ✓ | ✓ | | | 6.94015E-22 |
| ENSG00000161031 | PGLYRP2 | | ✓ | ✓ | | | 1.21248E-21 |
| ENSG00000224940 | PRRT4 | | ✓ | ✓ | | | 2.42972E-21 |
| ENSG00000254416 | RP11-347E10.1 | | ✓ | ✓ | | | 3.74024E-21 |
| ENSG00000156521 | TYSND1 | | | ✓ | | | 4.99904E-21 |
| ENSG00000105974 | CAV1 | ✓ | ✓ | ✓ | ✓ | | 1.05E-20 |
| ENSG00000174469 | CNTNAP2 | | ✓ | | | | 1.0666E-20 |
| ENSG00000164989 | CCDC171 | | ✓ | | | | 1.07149E-20 |
| ENSG00000173868 | PHOSPHO1 | | ✓ | | | | 1.58043E-20 |
| ENSG00000134443 | GRP | | ✓ | ✓ | | | 2.45957E-20 |
| ENSG00000250488 | RP11-6C14.1 | | ✓ | ✓ | | | 3.80294E-20 |
| ENSG00000112149 | CD83 | | ✓ | ✓ | | | 6.38609E-20 |
| ENSG00000251002 | AE000661.37 | ✓ | ✓ | ✓ | | | 1.16E-19 |
| ENSG00000205808 | PLPP6 | | ✓ | ✓ | | | 1.44272E-19 |
| ENSG00000182667 | NTM | | | ✓ | | | 1.82569E-19 |
| ENSG00000157557 | ETS2 | | ✓ | | | | 3.33616E-19 |
| ENSG00000130558 | OLFM1 | | ✓ | | | | 4.14968E-19 |
| ENSG00000171954 | CYP4F22 | | ✓ | ✓ | | | 5.07476E-19 |
| ENSG00000134207 | SYT6 | ✓ | | | ✓ | | 1.80198E-18 |
| ENSG00000164619 | BMPER | | ✓ | ✓ | | | 2.62865E-18 |
| ENSG00000106686 | SPATA6L | | ✓ | ✓ | | | 4.54217E-18 |
| ENSG00000258343 | RP11-536G4.2 | | | ✓ | | | 6.12272E-18 |
| ENSG00000248636 | RP11-768F21.1 | ✓ | | ✓ | | | 3.95E-17 |
| ENSG00000056998 | GYG2 | | ✓ | | | | 4.19694E-17 |
| ENSG00000091490 | SEL1L3 | | ✓ | | | | 5.08411E-17 |
| ENSG00000198756 | COLGALT2 | | ✓ | ✓ | | | 7.01013E-17 |
| ENSG00000155011 | DKK2 | ✓ | | | ✓ | | 8.09793E-17 |
| ENSG00000106069 | CHN2 | ✓ | | | | | 9.9183E-17 |
| ENSG00000129991 | TNNI3 | ✓ | | ✓ | | | 1.18E-16 |
| ENSG00000119801 | YPEL5 | | ✓ | ✓ | | | 2.04405E-16 |
| ENSG00000178662 | CSRNP3 | | ✓ | ✓ | | | 4.50756E-16 |
| ENSG00000057294 | PKP2 | | ✓ | ✓ | | | 4.66194E-16 |
| ENSG00000117407 | ARTN | | | ✓ | | | 5.17229E-16 |
| ENSG00000171094 | ALK | | | ✓ | | | 8.06187E-16 |
| ENSG00000166828 | SCNN1G | | ✓ | | | | 8.39583E-16 |

| | | | | | | | |
|---|---|---|---|---|---|---|---|
| ENSG00000115425 | PECR | | ✓ | | | | 8.84119E-16 |
| ENSG00000196139 | AKR1C3 | | | ✓ | | | 1.62238E-15 |
| ENSG00000089847 | ANKRD24 | | ✓ | ✓ | | | 2.07098E-15 |
| ENSG00000272825 | LL21NC02-1C16.2 | | ✓ | ✓ | | | 2.24144E-15 |
| ENSG00000165061 | ZMAT4 | | ✓ | ✓ | | | 2.26561E-15 |
| ENSG00000117009 | KMO | | ✓ | ✓ | | | 3.99905E-15 |
| ENSG00000227039 | ITGB2-AS1 | | ✓ | ✓ | | | 4.44821E-15 |
| ENSG00000133740 | E2F5 | | ✓ | | | | 4.80124E-15 |
| ENSG00000123338 | NCKAP1L | ✓ | | ✓ | ✓ | | 5.73E-15 |
| ENSG00000251448 | RP11-71E19.2 | | ✓ | | | | 2.98838E-14 |
| ENSG00000171843 | MLLT3 | ✓ | | ✓ | ✓ | | 3.17E-14 |
| ENSG00000030419 | IKZF2 | | | ✓ | | | 3.40733E-14 |
| ENSG00000137714 | FDX1 | | | ✓ | | | 4.40229E-14 |
| ENSG00000160255 | ITGB2 | | ✓ | ✓ | | | 6.14647E-14 |
| ENSG00000165995 | CACNB2 | ✓ | ✓ | | | | 4.43E-13 |
| ENSG00000136167 | LCP1 | ✓ | | ✓ | | | 6.76E-13 |
| ENSG00000127324 | TSPAN8 | ✓ | | | | | 7.62154E-13 |
| ENSG00000112893 | MAN2A1 | ✓ | | ✓ | ✓ | | 9.27E-13 |
| ENSG00000080618 | CPB2 | | ✓ | ✓ | | | 1.28496E-12 |
| ENSG00000172216 | CEBPB | | | ✓ | | | 1.78562E-12 |
| ENSG00000227646 | STEAP2-AS1 | | ✓ | ✓ | | | 3.35547E-12 |
| ENSG00000186153 | WWOX | | ✓ | ✓ | | | 5.53705E-12 |
| ENSG00000110002 | VWA5A | | ✓ | | | | 5.74716E-12 |
| ENSG00000175893 | ZDHHC21 | | | ✓ | | | 5.80903E-12 |
| ENSG00000152580 | IGSF10 | ✓ | | | ✓ | | 6.3E-12 |
| ENSG00000165757 | KIAA1462 | | ✓ | ✓ | | | 6.38264E-12 |
| ENSG00000251468 | RP11-369K16.1 | | | ✓ | | | 6.98702E-12 |
| ENSG00000149809 | TM7SF2 | | | ✓ | | | 7.3975E-12 |
| ENSG00000204950 | LRRC10B | | | ✓ | | | 1.04651E-11 |
| ENSG00000163630 | SYNPR | | | ✓ | | | 1.14797E-11 |
| ENSG00000123360 | PDE1B | ✓ | | | | | 1.15245E-11 |
| ENSG00000124126 | PREX1 | | ✓ | | | | 1.50517E-11 |
| ENSG00000006459 | KDM7A | | ✓ | | | | 1.81153E-11 |
| ENSG00000132646 | PCNA | | | ✓ | | | 1.95097E-11 |
| ENSG00000227674 | LINC00355 | | ✓ | | | | 2.27519E-11 |
| ENSG00000154856 | APCDD1 | ✓ | ✓ | ✓ | | | 2.59E-11 |
| ENSG00000137571 | SLCO5A1 | | ✓ | ✓ | | | 2.65438E-11 |
| ENSG00000166535 | A2ML1 | | ✓ | | | | 3.44431E-11 |
| ENSG00000204518 | AADACL4 | | ✓ | | | | 3.79946E-11 |
| ENSG00000109854 | HTATIP2 | | | ✓ | | | 5.00605E-11 |
| ENSG00000130449 | ZSWIM6 | | | ✓ | | | 6.83266E-11 |
| ENSG00000149781 | FERMT3 | | | ✓ | | | 8.09278E-11 |
| ENSG00000165821 | SALL2 | | | ✓ | | | 8.85819E-11 |
| ENSG00000049247 | UTS2 | | ✓ | ✓ | | | 8.91389E-11 |
| ENSG00000065150 | IPO5 | | ✓ | ✓ | | | 1.00728E-10 |
| ENSG00000188039 | NWD1 | | ✓ | ✓ | | | 1.10394E-10 |

| | | | | | | | |
|---|---|---|---|---|---|---|---|
| ENSG00000197837 | HIST4H4 | | | √ | | | 1.76469E-10 |
| ENSG00000138074 | SLC5A6 | | √ | | | | 2.51593E-10 |
| ENSG00000144820 | ADGRG7 | | √ | | | | 3.10926E-10 |
| ENSG00000119537 | KDSR | | √ | √ | | | 3.31484E-10 |
| ENSG00000163497 | FEV | | | √ | | | 4.6034E-10 |
| ENSG00000121749 | TBC1D15 | √ | √ | √ | | | 5.6E-10 |
| ENSG00000120594 | PLXDC2 | | | √ | | | 7.59314E-10 |
| ENSG00000135362 | PRR5L | | √ | √ | | | 7.74615E-10 |
| ENSG00000163659 | TIPARP | | | √ | | | 9.79376E-10 |
| ENSG00000017483 | SLC38A5 | | √ | | | | 9.84105E-10 |
| ENSG00000253111 | RP11-136O12.2 | | √ | √ | | | 1.00669E-09 |
| ENSG00000003096 | KLHL13 | | √ | √ | | | 1.12884E-09 |
| ENSG00000132915 | PDE6A | | √ | | | | 1.15759E-09 |
| ENSG00000235903 | CPB2-AS1 | | | √ | | | 2.00718E-09 |
| ENSG00000120519 | SLC10A7 | | √ | √ | | | 2.0979E-09 |
| ENSG00000132541 | RIDA | | √ | √ | | | 2.12437E-09 |
| ENSG00000168032 | ENTPD3 | | √ | | | | 2.63505E-09 |
| ENSG00000164076 | CAMKV | | | √ | | | 3.43278E-09 |
| ENSG00000232053 | AC009784.3 | | √ | √ | | | 3.44743E-09 |
| ENSG00000143995 | MEIS1 | | | √ | | | 3.81007E-09 |
| ENSG00000077522 | ACTN2 | | | √ | | | 4.4529E-09 |
| ENSG00000147853 | AK3 | | | √ | | | 5.21043E-09 |
| ENSG00000088035 | ALG6 | | √ | | | | 6.22056E-09 |
| ENSG00000079557 | AFM | | | √ | | | 7.31248E-09 |
| ENSG00000173212 | MAB21L3 | √ | | √ | | | 7.68E-09 |
| ENSG00000047579 | DTNBP1 | | | √ | | | 8.04381E-09 |
| ENSG00000004848 | ARX | √ | | | √ | | 8.19E-09 |
| ENSG00000123607 | TTC21B | | | √ | | | 1.04635E-08 |
| ENSG00000083807 | SLC27A5 | | | √ | | | 1.06686E-08 |
| ENSG00000175984 | DENND2C | √ | √ | √ | | | 1.12E-08 |
| ENSG00000115942 | ORC2 | | | √ | | | 1.32159E-08 |
| ENSG00000026103 | FAS | | | √ | | | 1.56977E-08 |
| ENSG00000165804 | ZNF219 | | | √ | | | 1.62573E-08 |
| ENSG00000166869 | CHP2 | | √ | √ | | | 1.68188E-08 |
| ENSG00000151572 | ANO4 | | √ | | | | 2.23444E-08 |
| ENSG00000146049 | KAAG1 | | | √ | | | 2.23491E-08 |
| ENSG00000124243 | BCAS4 | | √ | √ | | | 2.38245E-08 |
| ENSG00000176826 | FKBP9P1 | | √ | √ | | | 2.4327E-08 |
| ENSG00000204099 | NEU4 | | | √ | | | 2.46535E-08 |
| ENSG00000128274 | A4GALT | √ | √ | √ | | | 2.75E-08 |
| ENSG00000224982 | TMEM233 | √ | | √ | | | 3.52E-08 |
| ENSG00000253696 | KBTBD11-OT1 | | | √ | | | 3.63877E-08 |
| ENSG00000100749 | VRK1 | | | √ | | | 3.73335E-08 |
| ENSG00000253880 | RP11-124B13.1 | | √ | | | | 3.89792E-08 |
| ENSG00000221977 | OR4E2 | √ | √ | √ | | | 4.18E-08 |
| ENSG00000109586 | GALNT7 | | | √ | | | 4.87064E-08 |

| | | | | | | | |
|---|---|---|---|---|---|---|---|
| ENSG00000127328 | RAB3IP | | | ✓ | | | 5.26995E-08 |
| ENSG00000118733 | OLFM3 | ✓ | | | ✓ | | 7.26E-08 |
| ENSG00000187754 | SSX7 | | ✓ | | | | 7.74866E-08 |
| ENSG00000105048 | TNNT1 | ✓ | | ✓ | | | 9.17E-08 |
| ENSG00000052850 | ALX4 | | ✓ | ✓ | | | 9.81894E-08 |
| ENSG00000110315 | RNF141 | | ✓ | ✓ | | | 1.01375E-07 |
| ENSG00000166313 | APBB1 | | ✓ | ✓ | | | 1.01867E-07 |
| ENSG00000173153 | ESRRA | | | ✓ | | | 1.17258E-07 |
| ENSG00000234056 | LINC00463 | ✓ | ✓ | | | | 0.000000133 |
| ENSG00000197641 | SERPINB13 | | ✓ | | | | 1.7618E-07 |
| ENSG00000233332 | RP4-799P18.2 | | | ✓ | | | 2.13453E-07 |
| ENSG00000237732 | AC010980.2 | | ✓ | ✓ | | | 2.59096E-07 |
| ENSG00000204228 | HSD17B8 | | | ✓ | | | 2.73718E-07 |
| ENSG00000187742 | SECISBP2 | | | ✓ | | | 2.88721E-07 |
| ENSG00000228705 | LINC00659 | | | ✓ | | | 2.98395E-07 |
| ENSG00000205045 | SLFN12L | | ✓ | | | | 3.27683E-07 |
| ENSG00000176788 | BASP1 | | | ✓ | | | 3.7958E-07 |
| ENSG00000090863 | GLG1 | | ✓ | ✓ | | | 4.42968E-07 |
| ENSG00000198400 | NTRK1 | | ✓ | | | | 4.57017E-07 |
| ENSG00000164684 | ZNF704 | | ✓ | ✓ | | | 4.67948E-07 |
| ENSG00000105880 | DLX5 | ✓ | | | ✓ | | 5.04728E-07 |
| ENSG00000215196 | AC091878.1 | | | ✓ | | | 5.04739E-07 |
| ENSG00000225508 | SSXP5 | | ✓ | | | | 5.11939E-07 |
| ENSG00000168216 | LMBRD1 | | ✓ | ✓ | | | 5.2785E-07 |
| ENSG00000173473 | SMARCC1 | | ✓ | ✓ | | | 5.46482E-07 |
| ENSG00000174032 | SLC25A30 | | | ✓ | | | 5.67252E-07 |
| ENSG00000154269 | ENPP3 | | | ✓ | | | 6.76464E-07 |
| ENSG00000165943 | MOAP1 | | | ✓ | | | 7.19933E-07 |
| ENSG00000133104 | SPG20 | ✓ | | | ✓ | | 0.000000722 |
| ENSG00000164442 | CITED2 | ✓ | | ✓ | ✓ | | 0.000000735 |
| ENSG00000138814 | PPP3CA | | | ✓ | | | 8.00211E-07 |
| ENSG00000184787 | UBE2G2 | | | ✓ | | | 8.21686E-07 |
| ENSG00000257953 | RP11-620J15.1 | | | ✓ | | | 8.3892E-07 |
| ENSG00000167588 | GPD1 | | | ✓ | | | 8.42354E-07 |
| ENSG00000185518 | SV2B | ✓ | ✓ | ✓ | | | 0.000000877 |
| ENSG00000164600 | NEUROD6 | | ✓ | ✓ | | | 9.43787E-07 |
| ENSG00000022556 | NLRP2 | | | ✓ | | | 9.43787E-07 |
| ENSG00000154065 | ANKRD29 | | ✓ | | | | 1.09761E-06 |
| ENSG00000156925 | ZIC3 | | | ✓ | | | 1.37472E-06 |
| ENSG00000138190 | EXOC6 | | | ✓ | | | 1.47909E-06 |
| ENSG00000154760 | SLFN13 | | | ✓ | | | 1.63408E-06 |
| ENSG00000100078 | PLA2G3 | | | ✓ | | | 2.00609E-06 |
| ENSG00000157168 | NRG1 | | ✓ | ✓ | | | 2.09845E-06 |
| ENSG00000172927 | MYEOV | ✓ | | ✓ | ✓ | | 0.00000228 |
| ENSG00000100593 | ISM2 | | ✓ | | | | 2.35424E-06 |
| ENSG00000189120 | SP6 | | | ✓ | | | 2.66799E-06 |

| | | | | | | | |
|---|---|---|---|---|---|---|---|
| ENSG00000143341 | HMCN1 | | ✓ | ✓ | | | 2.88318E-06 |
| ENSG00000111615 | KRR1 | | | ✓ | | | 3.05871E-06 |
| ENSG00000160131 | VMA21 | | | ✓ | | | 3.29678E-06 |
| ENSG00000180573 | HIST1H2AC | | ✓ | ✓ | | | 3.56662E-06 |
| ENSG00000125816 | NKX2-4 | ✓ | | | ✓ | | 3.79331E-06 |
| ENSG00000162434 | JAK1 | ✓ | | ✓ | | | 0.00000382 |
| ENSG00000198885 | ITPRIPL1 | | | ✓ | | | 3.98173E-06 |
| ENSG00000156049 | GNA14 | ✓ | ✓ | ✓ | ✓ | | 0.00000418 |
| ENSG00000231817 | LINC01198 | ✓ | | | | | 0.00000465 |
| ENSG00000059588 | TARBP1 | | | ✓ | | | 6.41141E-06 |
| ENSG00000148362 | C9orf142 | | | ✓ | | | 6.57357E-06 |
| ENSG00000125820 | NKX2-2 | ✓ | | ✓ | ✓ | | 0.00000667 |
| ENSG00000072657 | TRHDE | ✓ | ✓ | ✓ | | | 0.00000681 |
| ENSG00000236708 | AC073316.1 | | ✓ | | | | 7.79992E-06 |
| ENSG00000227115 | LINC01630 | | ✓ | | | | 8.4071E-06 |
| ENSG00000166311 | SMPD1 | | ✓ | ✓ | | | 8.42288E-06 |
| ENSG00000183154 | RP11-863K10.7 | | ✓ | | | | 8.7916E-06 |
| ENSG00000064300 | NGFR | | | ✓ | | | 8.94998E-06 |
| ENSG00000205835 | GMNC | ✓ | | | ✓ | | 9.49402E-06 |
| ENSG00000163995 | ABLIM2 | ✓ | ✓ | | | | 0.00000976 |
| ENSG00000167083 | GNGT2 | | ✓ | ✓ | | | 1.00169E-05 |
| ENSG00000163002 | NUP35 | | | ✓ | | | 1.05059E-05 |
| ENSG00000104687 | GSR | | | ✓ | | | 1.21588E-05 |
| ENSG00000101109 | STK4 | | | ✓ | | | 1.28825E-05 |
| ENSG00000229051 | RP5-952N6.1 | | ✓ | | | | 1.33643E-05 |
| ENSG00000131771 | PPP1R1B | | | ✓ | | | 1.35905E-05 |
| ENSG00000127989 | MTERF1 | ✓ | ✓ | ✓ | ✓ | | 0.0000137 |
| ENSG00000135519 | KCNH3 | | | ✓ | | | 1.58322E-05 |
| ENSG00000157538 | DSCR3 | | | ✓ | | | 1.75276E-05 |
| ENSG00000145604 | SKP2 | | | ✓ | | | 1.84982E-05 |
| ENSG00000120616 | EPC1 | | | ✓ | | | 1.87065E-05 |
| ENSG00000236069 | RP5-1011O1.3 | | ✓ | ✓ | | | 1.90183E-05 |
| ENSG00000123200 | ZC3H13 | | | ✓ | | | 1.90183E-05 |
| ENSG00000171877 | FRMD5 | ✓ | ✓ | | | | 0.0000202 |
| ENSG00000149150 | SLC43A1 | | ✓ | ✓ | | | 2.23158E-05 |
| ENSG00000250303 | RP11-356J5.12 | | ✓ | ✓ | | | 2.28726E-05 |
| ENSG00000125864 | BFSP1 | ✓ | | ✓ | | | 0.0000232 |
| ENSG00000173334 | TRIB1 | | | ✓ | | | 2.33065E-05 |
| ENSG00000188559 | RALGAPA2 | | | ✓ | | | 2.36678E-05 |
| ENSG00000105369 | CD79A | | ✓ | ✓ | | | 2.37481E-05 |
| ENSG00000259977 | AL121578.2 | | ✓ | | | | 2.53262E-05 |
| ENSG00000037280 | FLT4 | ✓ | | ✓ | | | 0.0000258 |
| ENSG00000105643 | ARRDC2 | | | ✓ | | | 2.61163E-05 |
| ENSG00000272755 | RP11-326G21.1 | | ✓ | | | | 2.73791E-05 |
| ENSG00000213366 | GSTM2 | | | ✓ | | | 2.78161E-05 |
| ENSG00000075914 | EXOSC7 | | ✓ | | | | 2.84856E-05 |

| | | | | | | | |
|---|---|---|---|---|---|---|---|
| ENSG00000165795 | NDRG2 | | | ✓ | | | 2.87527E-05 |
| ENSG00000163743 | RCHY1 | | | ✓ | | | 3.13329E-05 |
| ENSG00000101638 | ST8SIA5 | | ✓ | ✓ | | | 3.17475E-05 |
| ENSG00000267034 | RP11-384O8.1 | | ✓ | ✓ | | | 3.21255E-05 |
| ENSG00000253974 | NRG1-IT1 | | ✓ | ✓ | | | 3.29309E-05 |
| ENSG00000269933 | RP3-333A15.2 | | ✓ | ✓ | | | 3.67353E-05 |
| ENSG00000144749 | LRIG1 | | | ✓ | | | 3.77469E-05 |
| ENSG00000172554 | SNTG2 | | | ✓ | | | 3.85175E-05 |
| ENSG00000255664 | ARL6IP1P1 | | ✓ | ✓ | | | 3.88269E-05 |
| ENSG00000167995 | BEST1 | | ✓ | ✓ | | | 3.90162E-05 |
| ENSG00000181284 | TMEM102 | | ✓ | ✓ | | | 4.05E-05 |
| ENSG00000077254 | USP33 | | ✓ | ✓ | | | 4.2256E-05 |
| ENSG00000158373 | HIST1H2BD | | | ✓ | | | 4.35092E-05 |
| ENSG00000227418 | PCGEM1 | | | ✓ | | | 4.55434E-05 |
| ENSG00000196636 | SDHAF3 | | | ✓ | | | 4.71213E-05 |
| ENSG00000131037 | EPS8L1 | | | ✓ | | | 4.82747E-05 |
| ENSG00000183145 | RIPPLY3 | | | ✓ | | | 5.10006E-05 |
| ENSG00000008118 | CAMK1G | | ✓ | | | | 5.6628E-05 |
| ENSG00000100346 | CACNA1I | | ✓ | | | | 5.87924E-05 |
| ENSG00000167528 | ZNF641 | | | ✓ | | | 5.95535E-05 |
| ENSG00000122877 | EGR2 | | | ✓ | | | 6.08335E-05 |
| ENSG00000116679 | IVNS1ABP | ✓ | | ✓ | ✓ | | 0.000061 |
| ENSG00000180447 | GAS1 | | | ✓ | | | 6.88957E-05 |
| ENSG00000258661 | RP11-964E11.2 | ✓ | | | ✓ | | 7.05272E-05 |
| ENSG00000165312 | OTUD1 | | | ✓ | | | 7.17862E-05 |
| ENSG00000212766 | EWSAT1 | | ✓ | | | | 7.2995E-05 |
| ENSG00000248550 | OTX2-AS1 | ✓ | ✓ | | | | 0.0000746 |
| ENSG00000232599 | RP1-161N10.1 | | ✓ | | | | 7.60769E-05 |
| ENSG00000072954 | TMEM38A | | | ✓ | | | 7.69019E-05 |
| ENSG00000015413 | DPEP1 | | | ✓ | | | 7.7598E-05 |
| ENSG00000260714 | RP11-266L9.1 | | ✓ | ✓ | | | 8.56443E-05 |
| ENSG00000116748 | AMPD1 | | ✓ | ✓ | | | 0.000101791 |
| ENSG00000114503 | NCBP2 | | | ✓ | | | 0.00010708 |
| ENSG00000091583 | APOH | | ✓ | | | | 0.000109247 |
| ENSG00000267279 | RP11-879F14.2 | | | ✓ | | | 0.000122236 |
| ENSG00000189403 | HMGB1 | | | ✓ | | | 0.000122956 |
| ENSG00000106066 | CPVL | ✓ | | | | | 0.000136074 |
| ENSG00000163399 | ATP1A1 | | ✓ | ✓ | | | 0.000139892 |
| ENSG00000131174 | COX7B | | | ✓ | | | 0.00014537 |
| ENSG00000169621 | APLF | | ✓ | ✓ | | | 0.000149041 |
| ENSG00000175105 | ZNF654 | | | ✓ | | | 0.00015108 |
| ENSG00000215630 | GUSBP9 | | | ✓ | | | 0.00015374 |
| ENSG00000267041 | ZNF850 | | | ✓ | | | 0.000157833 |
| ENSG00000174652 | ZNF266 | | | ✓ | | | 0.00015876 |
| ENSG00000175536 | LIPT2 | | | ✓ | | | 0.000164576 |
| ENSG00000177301 | KCNA2 | | ✓ | ✓ | | | 0.000166845 |

| | | | | | | | |
|---|---|---|---|---|---|---|---|
| ENSG00000164307 | ERAP1 | | | ✓ | | | 0.000181435 |
| ENSG00000173093 | CCDC63 | | | ✓ | | | 0.000184015 |
| ENSG00000124191 | TOX2 | | ✓ | ✓ | | | 0.000185063 |
| ENSG00000135903 | PAX3 | | | ✓ | | | 0.000191942 |
| ENSG00000164180 | TMEM161B | | | ✓ | | | 0.000193226 |
| ENSG00000140263 | SORD | ✓ | ✓ | | ✓ | | 0.00019842 |
| ENSG00000188906 | LRRK2 | ✓ | | | ✓ | | 0.000204013 |
| ENSG00000197299 | BLM | | | ✓ | | | 0.000210158 |
| ENSG00000168275 | COA6 | | | ✓ | | | 0.000213081 |
| ENSG00000197961 | ZNF121 | | | ✓ | | | 0.000216459 |
| ENSG00000146701 | MDH2 | | | ✓ | | | 0.000217639 |
| ENSG00000224577 | LINC01117 | | ✓ | ✓ | | | 0.000241417 |
| ENSG00000145555 | MYO10 | | ✓ | ✓ | | | 0.000247531 |
| ENSG00000105851 | PIK3CG | | | ✓ | | | 0.000248443 |
| ENSG00000228630 | HOTAIR | | | ✓ | | | 0.000257053 |
| ENSG00000162631 | NTNG1 | ✓ | ✓ | ✓ | ✓ | | 0.000261223 |
| ENSG00000197457 | STMN3 | | | ✓ | | | 0.000265556 |
| ENSG00000114315 | HES1 | | | ✓ | | | 0.00026609 |
| ENSG00000162924 | REL | | | ✓ | | | 0.000282605 |
| ENSG00000139287 | TPH2 | | | ✓ | | | 0.00028747 |
| ENSG00000267040 | RP11-35G9.3 | | | ✓ | | | 0.000298073 |
| ENSG00000259110 | RP11-359N5.1 | | ✓ | | | | 0.000300787 |
| ENSG00000164062 | APEH | | | ✓ | | | 0.000308328 |
| ENSG00000130312 | MRPL34 | | | ✓ | | | 0.000311371 |
| ENSG00000176927 | EFCAB5 | ✓ | ✓ | ✓ | | | 0.000312989 |
| ENSG00000121579 | NAA50 | | | ✓ | | | 0.000314261 |
| ENSG00000090776 | EFNB1 | | | ✓ | | | 0.000322572 |
| ENSG00000163812 | ZDHHC3 | | ✓ | | | | 0.000332973 |
| ENSG00000124635 | HIST1H2BJ | | | ✓ | | | 0.000340889 |
| ENSG00000128714 | HOXD13 | ✓ | | | | | 0.000341556 |
| ENSG00000128609 | NDUFA5 | | | ✓ | | | 0.000347381 |
| ENSG00000073169 | SELO | | | ✓ | | | 0.000349013 |
| ENSG00000165819 | METTL3 | ✓ | | ✓ | | | 0.000395918 |
| ENSG00000168918 | INPP5D | | | ✓ | | | 0.000458332 |
| ENSG00000032742 | IFT88 | | | ✓ | | | 0.000468468 |
| ENSG00000143319 | ISG20L2 | | | ✓ | | | 0.000498129 |
| ENSG00000168298 | HIST1H1E | | ✓ | ✓ | | | 0.000505598 |
| ENSG00000117143 | UAP1 | ✓ | | ✓ | ✓ | | 0.000525238 |
| ENSG00000132849 | PATJ | ✓ | | ✓ | | | 0.000537809 |
| ENSG00000115687 | PASK | | | ✓ | | | 0.000545085 |
| ENSG00000106268 | NUDT1 | | | ✓ | | | 0.000549163 |
| ENSG00000104731 | KLHDC4 | | | ✓ | | | 0.000553852 |
| ENSG00000198807 | PAX9 | ✓ | | | ✓ | | 0.000558268 |
| ENSG00000163623 | NKX6-1 | | | ✓ | | | 0.00055925 |
| ENSG00000167904 | TMEM68 | | | ✓ | | | 0.000570492 |
| ENSG00000135299 | ANKRD6 | | | ✓ | | | 0.000574079 |

| | | | | | | | |
|---|---|---|---|---|---|---|---|
| ENSG00000124784 | RIOK1 | | | √ | | | 0.000574685 |
| ENSG00000093072 | CECR1 | | | √ | | | 0.000580238 |
| ENSG00000188707 | ZBED6CL | | √ | | | | 0.000626579 |
| ENSG00000183048 | SLC25A10 | | | √ | | | 0.000631468 |
| ENSG00000227693 | GSTM3P1 | √ | | √ | √ | | 0.000639624 |
| ENSG00000100938 | GMPR2 | | | √ | | | 0.000652808 |
| ENSG00000132357 | CARD6 | | | √ | | | 0.000667635 |
| ENSG00000166012 | TAF1D | | | √ | | | 0.000672101 |
| ENSG00000175879 | HOXD8 | | | √ | | | 0.000715124 |
| ENSG00000165480 | SKA3 | | | √ | | | 0.000715465 |
| ENSG00000165792 | METTL17 | | | √ | | | 0.000733713 |
| ENSG00000198673 | FAM19A2 | | | √ | | | 0.000745539 |
| ENSG00000135269 | TES | | | √ | | | 0.000747839 |
| ENSG00000175198 | PCCA | √ | √ | √ | √ | | 0.000781127 |
| ENSG00000266578 | RP11-838N2.5 | | | √ | | | 0.000786103 |
| ENSG00000166398 | KIAA0355 | | | √ | | | 0.000790142 |
| ENSG00000143942 | CHAC2 | | | √ | | | 0.000811389 |
| ENSG00000085840 | ORC1 | √ | | √ | | | 0.00085012 |
| ENSG00000123388 | HOXC11 | | | √ | | | 0.000874406 |
| ENSG00000044524 | EPHA3 | | | √ | | | 0.000876301 |
| ENSG00000237813 | AC002066.1 | √ | √ | | √ | | 0.000891772 |
| ENSG00000246223 | LINC01550 | | | √ | | | 0.000893002 |
| ENSG00000163320 | CGGBP1 | √ | | √ | | | 0.000933184 |
| ENSG00000148704 | VAX1 | √ | | √ | | | 0.000947199 |
| ENSG00000137871 | ZNF280D | √ | √ | | | | 0.000953556 |
| ENSG00000083097 | DOPEY1 | | | √ | | | 0.000982133 |
| ENSG00000161958 | FGF11 | | √ | √ | | | 0.000989905 |
| ENSG00000005436 | GCFC2 | | | √ | | | 0.000990443 |
| ENSG00000238117 | AP004372.1 | | | √ | | | 0.001008956 |
| ENSG00000246528 | RP11-159H10.3 | | √ | √ | | | 0.001020617 |
| ENSG00000185238 | PRMT3 | | | √ | | | 0.001020658 |
| ENSG00000173598 | NUDT4 | | | √ | | | 0.00106793 |
| ENSG00000168067 | MAP4K2 | | | √ | | | 0.001073469 |
| ENSG00000244879 | GABPB1-AS1 | | | √ | | | 0.00109232 |
| ENSG00000167646 | DNAAF3 | √ | | √ | | | 0.00113 |
| ENSG00000236013 | RP3-332B22.1 | √ | √ | √ | | | 0.001135794 |
| ENSG00000182315 | MBD3L3 | | | √ | | | 0.001153285 |
| ENSG00000218351 | RPS3AP23 | √ | √ | √ | √ | | 0.001189315 |
| ENSG00000155592 | ZKSCAN2 | | | √ | | | 0.001233514 |
| ENSG00000095794 | CREM | | | √ | | | 0.001259529 |
| ENSG00000009844 | VTA1 | | | √ | | | 0.001316927 |
| ENSG00000168411 | RFWD3 | | | √ | | | 0.001381993 |
| ENSG00000164329 | PAPD4 | | | √ | | | 0.001401861 |
| ENSG00000105483 | CARD8 | | | √ | | | 0.001429337 |
| ENSG00000183283 | DAZAP2 | | | √ | | | 0.001437042 |
| ENSG00000211965 | IGHV3-49 | √ | √ | | | | 0.00148 |

| | | | | | | | |
|---|---|---|---|---|---|---|---|
| ENSG00000166821 | PEX11A | | | ✓ | | | 0.001486355 |
| ENSG00000253764 | RP11-439C15.4 | ✓ | | ✓ | | | 0.001524193 |
| ENSG00000175455 | CCDC14 | | | ✓ | | | 0.001527309 |
| ENSG00000180537 | RNF182 | | ✓ | ✓ | | | 0.001532469 |
| ENSG00000101972 | STAG2 | ✓ | | | ✓ | | 0.001537559 |
| ENSG00000267390 | RP11-635N19.1 | | ✓ | ✓ | | | 0.001540675 |
| ENSG00000178722 | C5orf64 | | ✓ | ✓ | | | 0.001572541 |
| ENSG00000159224 | GIP | | ✓ | ✓ | | | 0.001603139 |
| ENSG00000153774 | CFDP1 | | | ✓ | | | 0.001606787 |
| ENSG00000200879 | SNORD14E | | | ✓ | | | 0.001758792 |
| ENSG00000115902 | SLC1A4 | | ✓ | | | | 0.001885536 |
| ENSG00000189046 | ALKBH2 | | | ✓ | | | 0.001945853 |
| ENSG00000136521 | NDUFB5 | | | ✓ | | | 0.001963373 |
| ENSG00000187239 | FNBP1 | | | ✓ | | | 0.001977827 |
| ENSG00000260231 | JHDM1D-AS1 | | ✓ | | | | 0.001992557 |
| ENSG00000108001 | EBF3 | | | ✓ | | | 0.002024033 |
| ENSG00000240792 | RP11-418B12.1 | ✓ | ✓ | | ✓ | | 0.002109119 |
| ENSG00000124802 | EEF1E1 | | | ✓ | | | 0.002128353 |
| ENSG00000235996 | RP11-97N19.2 | ✓ | ✓ | ✓ | | | 0.002183569 |
| ENSG00000115875 | SRSF7 | | | ✓ | | | 0.002220828 |
| ENSG00000221978 | CCNL2 | | | ✓ | | | 0.002246777 |
| ENSG00000123689 | G0S2 | | | ✓ | | | 0.002321853 |
| ENSG00000125630 | POLR1B | | | ✓ | | | 0.002385161 |
| ENSG00000214553 | LRRC37A11P | | ✓ | | | | 0.002420169 |
| ENSG00000142856 | ITGB3BP | | | ✓ | | | 0.002506984 |
| ENSG00000176907 | C8orf4 | | | ✓ | | | 0.002584607 |
| ENSG00000214145 | LINC00887 | | | ✓ | | | 0.002844273 |
| ENSG00000172236 | TPSAB1 | | ✓ | | | | 0.00290887 |
| ENSG00000072832 | CRMP1 | | | ✓ | | | 0.002916684 |
| ENSG00000242808 | SOX2-OT | | | ✓ | | | 0.002978382 |
| ENSG00000070018 | LRP6 | | ✓ | | | | 0.002996734 |
| ENSG00000149716 | ORAOV1 | ✓ | | ✓ | ✓ | | 0.00300732 |
| ENSG00000187323 | DCC | | | ✓ | | | 0.003058328 |
| ENSG00000170270 | GON7 | | | ✓ | | | 0.003205035 |
| ENSG00000183434 | TFDP3 | | ✓ | | | | 0.003244031 |
| ENSG00000183032 | SLC25A21 | ✓ | ✓ | | | | 0.003244775 |
| ENSG00000101868 | POLA1 | ✓ | | ✓ | | | 0.003268045 |
| ENSG00000147465 | STAR | | ✓ | | | | 0.00338853 |
| ENSG00000132424 | PNISR | | | ✓ | | | 0.003414813 |
| ENSG00000118965 | WDR35 | | | ✓ | | | 0.003429975 |
| ENSG00000188001 | TPRG1 | ✓ | ✓ | ✓ | ✓ | | 0.00349 |
| ENSG00000205544 | TMEM256 | | | ✓ | | | 0.003682936 |
| ENSG00000135387 | CAPRIN1 | | ✓ | ✓ | | | 0.003727482 |
| ENSG00000125863 | MKKS | | | ✓ | | | 0.004057113 |
| ENSG00000232310 | RP11-557H15.4 | | ✓ | | | | 0.004107333 |
| ENSG00000235590 | GNAS-AS1 | ✓ | | ✓ | | | 0.0041364 |

| | | | | | | | |
|---|---|---|---|---|---|---|---|
| ENSG00000138653 | NDST4 | | | ✓ | | | 0.004299369 |
| ENSG00000187783 | TMEM72 | | | ✓ | | | 0.004348296 |
| ENSG00000198053 | SIRPA | | | ✓ | | | 0.004385303 |
| ENSG00000184575 | XPOT | | | ✓ | | | 0.004471173 |
| ENSG00000092140 | G2E3 | | | ✓ | | | 0.004545705 |
| ENSG00000177034 | MTX3 | | | ✓ | | | 0.004545705 |
| ENSG00000101890 | GUCY2F | | ✓ | | | | 0.004592215 |
| ENSG00000173376 | NDNF | | | ✓ | | | 0.004606197 |
| ENSG00000112874 | NUDT12 | | | ✓ | | | 0.004661856 |
| ENSG00000154265 | ABCA5 | | ✓ | ✓ | | | 0.004803446 |
| ENSG00000005801 | ZNF195 | | | ✓ | | | 0.004876313 |
| ENSG00000169783 | LINGO1 | ✓ | ✓ | ✓ | ✓ | | 0.00501 |
| ENSG00000167196 | FBXO22 | | | ✓ | | | 0.005230057 |
| ENSG00000260255 | RP11-185O21.1 | | | ✓ | | | 0.005242209 |
| ENSG00000177459 | ERICH5 | | | ✓ | | | 0.005349487 |
| ENSG00000089902 | RCOR1 | | ✓ | ✓ | | | 0.005463509 |
| ENSG00000163882 | POLR2H | | | ✓ | | | 0.005521081 |
| ENSG00000183674 | LINC00518 | | | ✓ | | | 0.005598013 |
| ENSG00000176102 | CSTF3 | | | ✓ | | | 0.005821177 |
| ENSG00000176853 | FAM91A1 | | | ✓ | | | 0.005821177 |
| ENSG00000164818 | DNAAF5 | | | ✓ | | | 0.006011717 |
| ENSG00000062822 | POLD1 | | | ✓ | | | 0.006047594 |
| ENSG00000103534 | TMC5 | ✓ | | | | | 0.006071386 |
| ENSG00000128652 | HOXD3 | | | ✓ | | | 0.006181566 |
| ENSG00000211776 | TRAV2 | ✓ | | | ✓ | | 0.006196245 |
| ENSG00000232000 | CLCN3P1 | | ✓ | ✓ | | | 0.006211278 |
| ENSG00000123009 | NME2P1 | | ✓ | ✓ | | | 0.006305577 |
| ENSG00000186487 | MYT1L | ✓ | ✓ | ✓ | | | 0.006306826 |
| ENSG00000230552 | AC092162.1 | ✓ | | ✓ | | | 0.006346361 |
| ENSG00000164967 | RPP25L | | | ✓ | | | 0.006494381 |
| ENSG00000101019 | UQCC1 | ✓ | | ✓ | | | 0.00663993 |
| ENSG00000117154 | IGSF21 | | ✓ | | | | 0.006749347 |
| ENSG00000245213 | RP11-10K16.1 | | | ✓ | | | 0.007052881 |
| ENSG00000248479 | RP11-807H7.2 | | ✓ | | | | 0.007054872 |
| ENSG00000233896 | RP4-684O24.5 | | ✓ | ✓ | | | 0.007223239 |
| ENSG00000250988 | SNHG21 | | | ✓ | | | 0.007573133 |
| ENSG00000188428 | BLOC1S5 | | | ✓ | | | 0.007701945 |
| ENSG00000128272 | ATF4 | | | ✓ | | | 0.00786721 |
| ENSG00000204356 | NELFE | | | ✓ | | | 0.007885553 |
| ENSG00000233862 | AC016907.3 | | ✓ | ✓ | | | 0.007966568 |
| ENSG00000171606 | ZNF274 | | | ✓ | | | 0.008205182 |
| ENSG00000165240 | ATP7A | | | ✓ | | | 0.00839978 |
| ENSG00000225783 | MIAT | | | ✓ | | | 0.008487657 |
| ENSG00000168434 | COG7 | | | ✓ | | | 0.008514453 |
| ENSG00000211961 | IGHV1-45 | ✓ | | | ✓ | | 0.008537371 |
| ENSG00000197555 | SIPA1L1 | | ✓ | ✓ | | | 0.008592953 |

| | | | | | | | |
|---|---|---|---|---|---|---|---|
| ENSG00000172840 | PDP2 | | ✓ | ✓ | | | 0.008830307 |
| ENSG00000104660 | LEPROTL1 | | | ✓ | | | 0.009074216 |
| ENSG00000092850 | TEKT2 | | | ✓ | | | 0.009147179 |
| ENSG00000164081 | TEX264 | | ✓ | | | | 0.009163876 |
| ENSG00000159208 | CIART | | | ✓ | | | 0.009168513 |
| ENSG00000170906 | NDUFA3 | | | ✓ | | | 0.009300474 |
| ENSG00000164284 | GRPEL2 | | | ✓ | | | 0.009906054 |
| ENSG00000184304 | PRKD1 | | ✓ | ✓ | | | 0.009990719 |
| ENSG00000179091 | CYC1 | | | ✓ | | | 0.0106213 |
| ENSG00000101361 | NOP56 | | | ✓ | | | 0.01070943 |
| ENSG00000267215 | RP11-685A21.1 | | ✓ | ✓ | | | 0.010828628 |
| ENSG00000232243 | RP11-408E5.5 | | | ✓ | | | 0.011268587 |
| ENSG00000263724 | DLGAP1-AS3 | | | ✓ | | | 0.011385071 |
| ENSG00000145777 | TSLP | | | ✓ | | | 0.011386085 |
| ENSG00000129071 | MBD4 | | ✓ | ✓ | | | 0.011423958 |
| ENSG00000139800 | ZIC5 | | | ✓ | | | 0.011589532 |
| ENSG00000173559 | NABP1 | | | ✓ | | | 0.011604508 |
| ENSG00000135119 | RNFT2 | | | ✓ | | | 0.011614405 |
| ENSG00000168016 | TRANK1 | | | ✓ | | | 0.011719695 |
| ENSG00000144908 | ALDH1L1 | | | ✓ | | | 0.011921141 |
| ENSG00000176105 | YES1 | | | ✓ | | | 0.011976708 |
| ENSG00000224367 | OACYLP | | | ✓ | | | 0.012365818 |
| ENSG00000095059 | DHPS | ✓ | | ✓ | | | 0.012451109 |
| ENSG00000225039 | LINC01058 | | ✓ | ✓ | | | 0.012509216 |
| ENSG00000165609 | NUDT5 | | | ✓ | | | 0.013187198 |
| ENSG00000204348 | DXO | | | ✓ | | | 0.013219723 |
| ENSG00000118515 | SGK1 | | | ✓ | | | 0.01322162 |
| ENSG00000149761 | NUDT22 | | | ✓ | | | 0.013395718 |
| ENSG00000247240 | UBL7-AS1 | | | ✓ | | | 0.013398835 |
| ENSG00000170345 | FOS | | ✓ | | | | 0.013422353 |
| ENSG00000100985 | MMP9 | | ✓ | ✓ | | | 0.013802486 |
| ENSG00000196189 | SEMA4A | ✓ | | ✓ | | | 0.0139 |
| ENSG00000231721 | LINC-PINT | | ✓ | ✓ | | | 0.01420732 |
| ENSG00000253438 | PCAT1 | | | ✓ | | | 0.014247454 |
| ENSG00000149346 | SLX4IP | | | ✓ | | | 0.014416394 |
| ENSG00000212366 | RNU6-1246P | | ✓ | ✓ | | | 0.01465573 |
| ENSG00000148942 | SLC5A12 | | ✓ | ✓ | | | 0.014726631 |
| ENSG00000121989 | ACVR2A | | | ✓ | | | 0.014742309 |
| ENSG00000147804 | SLC39A4 | | | ✓ | | | 0.014770513 |
| ENSG00000163376 | KBTBD8 | | | ✓ | | | 0.014836342 |
| ENSG00000250791 | RP11-55L3.1 | | | ✓ | | | 0.015007747 |
| ENSG00000111897 | SERINC1 | | | ✓ | | | 0.015164552 |
| ENSG00000089057 | SLC23A2 | | ✓ | | | | 0.015264164 |
| ENSG00000152147 | GEMIN6 | | ✓ | ✓ | | | 0.015275504 |
| ENSG00000166261 | ZNF202 | | | ✓ | | | 0.0153929 |
| ENSG00000197410 | DCHS2 | | ✓ | | | | 0.01557743 |

| | | | | | | | |
|---|---|---|---|---|---|---|---|
| ENSG00000111364 | DDX55 | | | ✓ | | | 0.015861781 |
| ENSG00000235695 | RP11-395P16.1 | | ✓ | ✓ | | | 0.015906856 |
| ENSG00000152332 | UHMK1 | | | ✓ | | | 0.015938041 |
| ENSG00000138670 | RASGEF1B | | | ✓ | | | 0.015941984 |
| ENSG00000198818 | SFT2D1 | | | ✓ | | | 0.016008869 |
| ENSG00000242593 | RP5-921G16.1 | | ✓ | ✓ | | | 0.016120119 |
| ENSG00000251857 | RNA5SP362 | ✓ | ✓ | ✓ | | | 0.0166 |
| ENSG00000124587 | PEX6 | | | ✓ | | | 0.016764063 |
| ENSG00000160688 | FLAD1 | | | ✓ | | | 0.016939915 |
| ENSG00000183971 | NPW | | | ✓ | | | 0.017050763 |
| ENSG00000164796 | CSMD3 | | | ✓ | | | 0.017366746 |
| ENSG00000159176 | CSRP1 | | ✓ | | | | 0.017370894 |
| ENSG00000250436 | RP11-622A1.2 | | | ✓ | | | 0.017878396 |
| ENSG00000101752 | MIB1 | | ✓ | | | | 0.018339165 |
| ENSG00000134802 | SLC43A3 | | | ✓ | | | 0.018818507 |
| ENSG00000254286 | RP11-89K10.1 | ✓ | ✓ | ✓ | ✓ | | 0.018958752 |
| ENSG00000232677 | LINC00665 | | | ✓ | | | 0.019196553 |
| ENSG00000149485 | FADS1 | | | ✓ | | | 0.019198698 |
| ENSG00000257218 | GATC | | | ✓ | | | 0.019351089 |
| ENSG00000128713 | HOXD11 | ✓ | | | ✓ | | 0.019432927 |
| ENSG00000250838 | RP11-501C14.5 | | ✓ | ✓ | | | 0.019463615 |
| ENSG00000086548 | CEACAM6 | | ✓ | ✓ | | | 0.019634265 |
| ENSG00000196787 | HIST1H2AG | | | ✓ | | | 0.020353292 |
| ENSG00000077514 | POLD3 | | | ✓ | | | 0.020499207 |
| ENSG00000137574 | TGS1 | | | ✓ | | | 0.021125606 |
| ENSG00000162227 | TAF6L | | | ✓ | | | 0.021205005 |
| ENSG00000167384 | ZNF180 | | ✓ | ✓ | | | 0.021358751 |
| ENSG00000102531 | FNDC3A | | | ✓ | | | 0.021441656 |
| ENSG00000197253 | TPSB2 | | ✓ | | | | 0.021612232 |
| ENSG00000163864 | NMNAT3 | | | ✓ | | | 0.021949553 |
| ENSG00000170231 | FABP6 | | | ✓ | | | 0.022891486 |
| ENSG00000142623 | PADI1 | | ✓ | | | | 0.02293395 |
| ENSG00000162086 | ZNF75A | | | ✓ | | | 0.023002059 |
| ENSG00000163322 | FAM175A | | | ✓ | | | 0.023252689 |
| ENSG00000125885 | MCM8 | | | ✓ | | | 0.023369682 |
| ENSG00000145388 | METTL14 | | | ✓ | | | 0.023615022 |
| ENSG00000049541 | RFC2 | | | ✓ | | | 0.024945179 |
| ENSG00000107779 | BMPR1A | | | ✓ | | | 0.025116358 |
| ENSG00000159217 | IGF2BP1 | | ✓ | ✓ | | | 0.025394483 |
| ENSG00000196878 | LAMB3 | | ✓ | ✓ | | | 0.026020545 |
| ENSG00000141837 | CACNA1A | | ✓ | | | | 0.02644947 |
| ENSG00000147419 | CCDC25 | | | ✓ | | | 0.02644947 |
| ENSG00000132485 | ZRANB2 | | | ✓ | | | 0.026673951 |
| ENSG00000166508 | MCM7 | | | ✓ | | | 0.026785434 |
| ENSG00000159086 | PAXBP1 | | | ✓ | | | 0.026958122 |
| ENSG00000167220 | HDHD2 | | | ✓ | | | 0.027276336 |

| | | | | | | | |
|---|---|---|---|---|---|---|---|
| ENSG00000161996 | WDR90 | | | ✓ | | | 0.027296003 |
| ENSG00000125967 | NECAB3 | | | ✓ | | | 0.027355736 |
| ENSG00000158296 | SLC13A3 | | | ✓ | | | 0.027563841 |
| ENSG00000169718 | DUS1L | | | ✓ | | | 0.027748822 |
| ENSG00000253203 | GUSBP3 | | ✓ | | | | 0.027829208 |
| ENSG00000221914 | PPP2R2A | ✓ | | ✓ | | | 0.028374755 |
| ENSG00000104915 | STX10 | | | ✓ | | | 0.02845857 |
| ENSG00000147874 | HAUS6 | | | ✓ | | | 0.028591601 |
| ENSG00000173660 | UQCRH | | | ✓ | | | 0.029002563 |
| ENSG00000118680 | MYL12B | | | ✓ | | | 0.029002563 |
| ENSG00000149476 | TKFC | | | ✓ | | | 0.029334075 |
| ENSG00000235092 | ID2-AS1 | ✓ | | ✓ | | | 0.029443656 |
| ENSG00000109084 | TMEM97 | | | ✓ | | | 0.030053496 |
| ENSG00000108106 | UBE2S | | | ✓ | | | 0.030291726 |
| ENSG00000103994 | ZNF106 | | | ✓ | | | 0.03037316 |
| ENSG00000124571 | XPO5 | | | ✓ | | | 0.030402511 |
| ENSG00000144395 | CCDC150 | | | ✓ | | | 0.030662187 |
| ENSG00000144401 | METTL21A | | | ✓ | | | 0.031339578 |
| ENSG00000163938 | GNL3 | | | ✓ | | | 0.031589359 |
| ENSG00000172123 | SLFN12 | | | ✓ | | | 0.031589359 |
| ENSG00000050393 | MCUR1 | | | ✓ | | | 0.031901192 |
| ENSG00000237950 | RP11-7O11.3 | | | ✓ | | | 0.031901192 |
| ENSG00000149823 | VPS51 | | | ✓ | | | 0.031919366 |
| ENSG00000211892 | IGHG4 | | | ✓ | | | 0.032307142 |
| ENSG00000143756 | FBXO28 | | | ✓ | | | 0.033508787 |
| ENSG00000249878 | CTD-2244C20.1 | | ✓ | ✓ | | | 0.033802808 |
| ENSG00000070476 | ZXDC | | | ✓ | | | 0.03395039 |
| ENSG00000130303 | BST2 | | | ✓ | | | 0.03415668 |
| ENSG00000238169 | LINC01053 | ✓ | ✓ | | | | 0.034243151 |
| ENSG00000105372 | RPS19 | | ✓ | ✓ | | | 0.035814417 |
| ENSG00000174669 | SLC29A2 | | ✓ | | | | 0.036325122 |
| ENSG00000130561 | SAG | | ✓ | ✓ | | | 0.036441734 |
| ENSG00000052723 | SIKE1 | | | ✓ | | | 0.03739427 |
| ENSG00000138101 | DTNB | | ✓ | | | | 0.037518138 |
| ENSG00000103005 | USB1 | | | ✓ | | | 0.037700851 |
| ENSG00000144034 | TPRKB | | | ✓ | | | 0.038362419 |
| ENSG00000254143 | RP11-470M17.2 | | | ✓ | | | 0.038729378 |
| ENSG00000237401 | LINC01304 | ✓ | | ✓ | ✓ | | 0.039325318 |
| ENSG00000132680 | KIAA0907 | | | ✓ | | | 0.039412534 |
| ENSG00000100246 | DNAL4 | | ✓ | ✓ | | | 0.039909514 |
| ENSG00000216937 | CCDC7 | ✓ | | ✓ | ✓ | | 0.040000641 |
| ENSG00000170638 | TRABD | | | ✓ | | | 0.040096551 |
| ENSG00000140859 | KIFC3 | | ✓ | ✓ | | | 0.040165959 |
| ENSG00000103356 | EARS2 | | | ✓ | | | 0.040331093 |
| ENSG00000259042 | AE000661.50 | | ✓ | ✓ | | | 0.040762671 |
| ENSG00000138769 | CDKL2 | | ✓ | | | | 0.041128745 |

| | | | | | | | |
|---|---|---|---|---|---|---|---|
| ENSG00000167964 | RAB26 | | | ✓ | | | 0.041313236 |
| ENSG00000236751 | LINC01186 | | ✓ | ✓ | | | 0.041369204 |
| ENSG00000076928 | ARHGEF1 | | ✓ | ✓ | | | 0.041524461 |
| ENSG00000113790 | EHHADH | | | ✓ | | | 0.042882034 |
| ENSG00000247626 | MARS2 | | | ✓ | | | 0.042976727 |
| ENSG00000149925 | ALDOA | | | ✓ | | | 0.04313866 |
| ENSG00000136270 | TBRG4 | | | ✓ | | | 0.043150691 |
| ENSG00000178295 | GEN1 | | | ✓ | | | 0.043175323 |
| ENSG00000169604 | ANTXR1 | | ✓ | ✓ | | | 0.044599569 |
| ENSG00000135390 | ATP5G2 | | ✓ | | | | 0.04478402 |
| ENSG00000246022 | ALDH1L1-AS2 | | | ✓ | | | 0.04478402 |
| ENSG00000225492 | GBP1P1 | | ✓ | | | | 0.04516779 |
| ENSG00000114573 | ATP6V1A | | | ✓ | | | 0.04525793 |
| ENSG00000049246 | PER3 | | ✓ | ✓ | | | 0.045396171 |
| ENSG00000197903 | HIST1H2BK | | | ✓ | | | 0.045666781 |
| ENSG00000204444 | APOM | | | ✓ | | | 0.045813206 |
| ENSG00000236780 | AC078941.1 | ✓ | | | ✓ | | 0.046397248 |
| ENSG00000181418 | DDN | | | ✓ | | | 0.046919605 |
| ENSG00000237053 | FUCA1P1 | | | ✓ | | | 0.047537216 |
| ENSG00000012504 | NR1H4 | | ✓ | ✓ | | | 0.049358773 |
| ENSG00000258913 | RP11-260M19.2 | | | ✓ | | | 0.049420466 |

MS = GGAA-µsat, NMS = non-µsat, p-adj = adjusted p-value ( < 0.05)

**Supplementary table S2. Genes directly downregulated by EWS/FLI in A673**

| Ensembl ID | Gene symbol | 20kb gained loop | EWS/FLI binding site | | | | p-adj (EF-Endo vs EF-KD) |
|---|---|---|---|---|---|---|---|
| | | | Proximal | | Distal | | |
| | | | MS | NMS | MS | NMS | |
| ENSG00000136158 | SPRY2 | | | ✓ | | | 2.69627E-77 |
| ENSG00000135363 | LMO2 | ✓ | | | | ✓ | 4.04E-58 |
| ENSG00000144824 | PHLDB2 | ✓ | | ✓ | | | 6.85E-54 |
| ENSG00000075213 | SEMA3A | | | ✓ | | | 9.27395E-45 |
| ENSG00000150687 | PRSS23 | ✓ | | ✓ | ✓ | | 7.69E-42 |
| ENSG00000151491 | EPS8 | | | ✓ | | | 1.91435E-36 |
| ENSG00000039560 | RAI14 | | | ✓ | | | 1.3673E-33 |
| ENSG00000181982 | CCDC149 | | | ✓ | | | 3.79083E-33 |
| ENSG00000188452 | CERKL | | ✓ | | | | 2.94472E-30 |
| ENSG00000064042 | LIMCH1 | | ✓ | ✓ | | | 2.13458E-25 |
| ENSG00000075223 | SEMA3C | | | ✓ | | | 2.73347E-25 |
| ENSG00000123989 | CHPF | | | ✓ | | | 5.42718E-25 |
| ENSG00000181104 | F2R | | ✓ | | | | 2.60693E-24 |
| ENSG00000138347 | MYPN | | ✓ | | | | 2.06314E-23 |
| ENSG00000120833 | SOCS2 | | | ✓ | | | 1.29662E-22 |
| ENSG00000144724 | PTPRG | | | ✓ | | | 2.52453E-22 |
| ENSG00000106397 | PLOD3 | | | ✓ | | | 5.23517E-22 |
| ENSG00000185624 | P4HB | | | ✓ | | | 3.43677E-21 |
| ENSG00000150347 | ARID5B | ✓ | | | ✓ | ✓ | 3.87E-21 |
| ENSG00000116991 | SIPA1L2 | | | ✓ | | | 3.00794E-20 |
| ENSG00000205730 | ITPRIPL2 | | | ✓ | | | 2.4538E-19 |
| ENSG00000150551 | LYPD1 | | | ✓ | | | 2.49615E-18 |
| ENSG00000146147 | MLIP | | ✓ | | | | 5.97417E-18 |
| ENSG00000148154 | UGCG | | | ✓ | | | 3.61502E-17 |
| ENSG00000170445 | HARS | | | ✓ | | | 5.71288E-17 |
| ENSG00000165617 | DACT1 | | ✓ | | | | 5.88569E-17 |
| ENSG00000128595 | CALU | | | ✓ | | | 6.45929E-17 |
| ENSG00000154188 | ANGPT1 | | ✓ | | | | 5.37258E-16 |
| ENSG00000181826 | RELL1 | | | ✓ | | | 1.09588E-15 |
| ENSG00000189129 | PLAC9 | | | ✓ | | | 1.40804E-15 |
| ENSG00000138674 | SEC31A | | | ✓ | | | 4.53196E-15 |
| ENSG00000137573 | SULF1 | | | ✓ | | | 1.96688E-14 |
| ENSG00000102554 | KLF5 | | | ✓ | | | 3.10617E-14 |
| ENSG00000172380 | GNG12 | | | ✓ | | | 5.64097E-14 |
| ENSG00000095752 | IL11 | | | ✓ | | | 7.71635E-14 |
| ENSG00000137331 | IER3 | ✓ | | ✓ | | ✓ | 2.47E-13 |
| ENSG00000235162 | C12orf75 | | | ✓ | | | 1.19038E-12 |
| ENSG00000160886 | LY6K | | | ✓ | | | 1.97291E-12 |
| ENSG00000120913 | PDLIM2 | | | ✓ | | | 2.79074E-12 |
| ENSG00000184349 | EFNA5 | | | ✓ | | | 4.13229E-12 |
| ENSG00000091136 | LAMB1 | | | ✓ | | | 6.72601E-12 |

| | | | | | | | |
|---|---|---|---|---|---|---|---|
| ENSG00000161638 | ITGA5 | ✓ | | ✓ | | ✓ | 9.02E-12 |
| ENSG00000122218 | COPA | | | ✓ | | | 9.77763E-12 |
| ENSG00000196159 | FAT4 | | | ✓ | | | 2.2824E-11 |
| ENSG00000027075 | PRKCH | ✓ | | ✓ | | | 4.49E-11 |
| ENSG00000165895 | ARHGAP42 | | | ✓ | | | 5.51939E-11 |
| ENSG00000142867 | BCL10 | | | ✓ | | | 5.54094E-11 |
| ENSG00000168702 | LRP1B | | | ✓ | | | 6.39544E-11 |
| ENSG00000018408 | WWTR1 | | | ✓ | | | 7.47586E-11 |
| ENSG00000196975 | ANXA4 | | | ✓ | | | 8.90418E-11 |
| ENSG00000158109 | TPRG1L | | | ✓ | | | 1.69968E-10 |
| ENSG00000170153 | RNF150 | ✓ | | ✓ | | | 1.9E-10 |
| ENSG00000182621 | PLCB1 | | | ✓ | | | 4.6101E-10 |
| ENSG00000197959 | DNM3 | | | ✓ | | | 4.90092E-10 |
| ENSG00000138835 | RGS3 | | ✓ | | | | 5.24854E-10 |
| ENSG00000135862 | LAMC1 | | | ✓ | | | 8.22572E-10 |
| ENSG00000137501 | SYTL2 | | | ✓ | | | 1.24337E-09 |
| ENSG00000128590 | DNAJB9 | | | ✓ | | | 1.78724E-09 |
| ENSG00000181789 | COPG1 | | | ✓ | | | 2.30781E-09 |
| ENSG00000177283 | FZD8 | | ✓ | ✓ | | | 2.55904E-09 |
| ENSG00000168675 | LDLRAD4 | | | ✓ | | | 4.56435E-09 |
| ENSG00000163479 | SSR2 | | | ✓ | | | 6.96006E-09 |
| ENSG00000116106 | EPHA4 | ✓ | | | ✓ | | 8.28346E-09 |
| ENSG00000184432 | COPB2 | | | ✓ | | | 9.32093E-09 |
| ENSG00000140612 | SEC11A | | | ✓ | | | 9.73731E-09 |
| ENSG00000137269 | LRRC1 | | | ✓ | | | 1.04211E-08 |
| ENSG00000115129 | TP53I3 | | | ✓ | | | 1.20156E-08 |
| ENSG00000122642 | FKBP9 | | ✓ | | | | 1.35357E-08 |
| ENSG00000095370 | SH2D3C | | | ✓ | | | 1.39155E-08 |
| ENSG00000163485 | ADORA1 | | | ✓ | | | 1.55367E-08 |
| ENSG00000173068 | BNC2 | | | ✓ | | | 1.93982E-08 |
| ENSG00000162576 | MXRA8 | | | ✓ | | | 2.43299E-08 |
| ENSG00000108387 | SEPT4 | | | ✓ | | | 2.93196E-08 |
| ENSG00000058262 | SEC61A1 | | | ✓ | | | 3.89153E-08 |
| ENSG00000130429 | ARPC1B | | | ✓ | | | 7.70269E-08 |
| ENSG00000139970 | RTN1 | | | ✓ | | | 9.07593E-08 |
| ENSG00000171940 | ZNF217 | | | ✓ | | | 1.26275E-07 |
| ENSG00000179222 | MAGED1 | | | ✓ | | | 1.33161E-07 |
| ENSG00000107796 | ACTA2 | | | ✓ | | | 1.50884E-07 |
| ENSG00000140945 | CDH13 | ✓ | | | ✓ | | 1.73727E-07 |
| ENSG00000109680 | TBC1D19 | | | ✓ | | | 4.0424E-07 |
| ENSG00000113721 | PDGFRB | | | ✓ | | | 4.12753E-07 |
| ENSG00000124120 | TTPAL | | | ✓ | | | 4.36788E-07 |
| ENSG00000150867 | PIP4K2A | | | ✓ | | | 8.72848E-07 |
| ENSG00000140682 | TGFB1I1 | | | ✓ | | | 1.09714E-06 |
| ENSG00000151240 | DIP2C | | | ✓ | | | 1.57594E-06 |

| | | | | | | | |
|---|---|---|---|---|---|---|---|
| ENSG00000196526 | AFAP1 | ✓ | | ✓ | ✓ | | 0.00000176 |
| ENSG00000108829 | LRRC59 | | | ✓ | | | 1.78161E-06 |
| ENSG00000077238 | IL4R | | | ✓ | | | 1.8528E-06 |
| ENSG00000102575 | ACP5 | | | ✓ | | | 2.0714E-06 |
| ENSG00000148180 | GSN | | | ✓ | | | 2.08133E-06 |
| ENSG00000137872 | SEMA6D | ✓ | ✓ | | ✓ | | 0.00000232 |
| ENSG00000242247 | ARFGAP3 | ✓ | | | ✓ | ✓ | 2.53515E-06 |
| ENSG00000171055 | FEZ2 | | | ✓ | | | 3.10695E-06 |
| ENSG00000076513 | ANKRD13A | | | ✓ | | | 3.51285E-06 |
| ENSG00000065485 | PDIA5 | | ✓ | ✓ | | | 4.01568E-06 |
| ENSG00000008196 | TFAP2B | | | ✓ | | | 4.03447E-06 |
| ENSG00000102359 | SRPX2 | | | ✓ | | | 4.78842E-06 |
| ENSG00000257219 | RP11-54A9.1 | | | ✓ | | | 5.34353E-06 |
| ENSG00000168615 | ADAM9 | | | ✓ | | | 5.60925E-06 |
| ENSG00000138760 | SCARB2 | | | ✓ | | | 5.63519E-06 |
| ENSG00000163697 | APBB2 | ✓ | | ✓ | | ✓ | 0.00000584 |
| ENSG00000112473 | SLC39A7 | | | ✓ | | | 5.89282E-06 |
| ENSG00000124006 | OBSL1 | | | ✓ | | | 6.09538E-06 |
| ENSG00000185989 | RASA3 | ✓ | | ✓ | | | 0.00000701 |
| ENSG00000134516 | DOCK2 | ✓ | ✓ | | ✓ | | 0.000008 |
| ENSG00000168374 | ARF4 | | | ✓ | | | 9.59481E-06 |
| ENSG00000166949 | SMAD3 | ✓ | | ✓ | | | 0.0000123 |
| ENSG00000229989 | MIR181A1HG | | ✓ | ✓ | | | 1.23013E-05 |
| ENSG00000071127 | WDR1 | | | ✓ | | | 1.49106E-05 |
| ENSG00000172037 | LAMB2 | | | ✓ | | | 1.50853E-05 |
| ENSG00000091129 | NRCAM | | | ✓ | | | 2.0245E-05 |
| ENSG00000197757 | HOXC6 | | | ✓ | | | 2.14E-05 |
| ENSG00000103855 | CD276 | | | ✓ | | | 2.37493E-05 |
| ENSG00000139971 | C14orf37 | ✓ | ✓ | | ✓ | ✓ | 0.0000254 |
| ENSG00000090530 | P3H2 | | | ✓ | | | 2.67966E-05 |
| ENSG00000182287 | AP1S2 | | | ✓ | | | 3.13159E-05 |
| ENSG00000166250 | CLMP | | | ✓ | | | 3.35602E-05 |
| ENSG00000064932 | SBNO2 | | | ✓ | | | 3.85914E-05 |
| ENSG00000107175 | CREB3 | | | ✓ | | | 3.90217E-05 |
| ENSG00000099999 | RNF215 | | | ✓ | | | 3.97321E-05 |
| ENSG00000166148 | AVPR1A | | ✓ | | | | 4.14129E-05 |
| ENSG00000132128 | LRRC41 | | | ✓ | | | 4.18921E-05 |
| ENSG00000137509 | PRCP | | | ✓ | | | 4.34582E-05 |
| ENSG00000223891 | OSER1-AS1 | | | ✓ | | | 5.18456E-05 |
| ENSG00000149289 | ZC3H12C | | | ✓ | | | 5.9167E-05 |
| ENSG00000115216 | NRBP1 | | | ✓ | | | 6.78283E-05 |
| ENSG00000005238 | FAM214B | | | ✓ | | | 6.97034E-05 |
| ENSG00000158828 | PINK1 | | | ✓ | | | 9.63464E-05 |
| ENSG00000149260 | CAPN5 | | | ✓ | | | 0.00010665 |
| ENSG00000130733 | YIPF2 | | | ✓ | | | 0.000118536 |

| | | | | | | | |
|---|---|---|---|---|---|---|---|
| ENSG00000130703 | OSBPL2 | | | ✓ | | | 0.000130199 |
| ENSG00000271122 | RP11-379H18.1 | | | ✓ | | | 0.000135694 |
| ENSG00000126878 | AIF1L | | | ✓ | | | 0.000139779 |
| ENSG00000130309 | COLGALT1 | | | ✓ | | | 0.000145282 |
| ENSG00000165915 | SLC39A13 | | | ✓ | | | 0.000183335 |
| ENSG00000143079 | CTTNBP2NL | | | ✓ | | | 0.000184268 |
| ENSG00000106976 | DNM1 | | | ✓ | | | 0.000212471 |
| ENSG00000168090 | COPS6 | | | ✓ | | | 0.000237044 |
| ENSG00000068912 | ERLEC1 | | | ✓ | | | 0.000256374 |
| ENSG00000090615 | GOLGA3 | | | ✓ | | | 0.000271567 |
| ENSG00000179889 | PDXDC1 | | | ✓ | | | 0.000291689 |
| ENSG00000169760 | NLGN1 | | ✓ | ✓ | | | 0.000343662 |
| ENSG00000113578 | FGF1 | | | ✓ | | | 0.000353076 |
| ENSG00000223768 | LINC00205 | | | ✓ | | | 0.000362412 |
| ENSG00000148248 | SURF4 | | | ✓ | | | 0.000372783 |
| ENSG00000122557 | HERPUD2 | | | ✓ | | | 0.000385794 |
| ENSG00000131018 | SYNE1 | | ✓ | | | | 0.000432738 |
| ENSG00000106636 | YKT6 | | | ✓ | | | 0.000444309 |
| ENSG00000156050 | FAM161B | | | ✓ | | | 0.000551858 |
| ENSG00000163466 | ARPC2 | | | ✓ | | | 0.000607826 |
| ENSG00000170776 | AKAP13 | ✓ | ✓ | ✓ | | | 0.0007 |
| ENSG00000095383 | TBC1D2 | | | ✓ | | | 0.000780056 |
| ENSG00000100325 | ASCC2 | | | ✓ | | | 0.000789669 |
| ENSG00000099250 | NRP1 | | | ✓ | | | 0.000810608 |
| ENSG00000072958 | AP1M1 | | | ✓ | | | 0.000814437 |
| ENSG00000138696 | BMPR1B | | ✓ | | | | 0.000829158 |
| ENSG00000130164 | LDLR | | | ✓ | | | 0.000845729 |
| ENSG00000186866 | POFUT2 | | | ✓ | | | 0.000852962 |
| ENSG00000215447 | BX322557.10 | | | ✓ | | | 0.000908296 |
| ENSG00000198198 | SZT2 | | | ✓ | | | 0.000989917 |
| ENSG00000006744 | ELAC2 | | | ✓ | | | 0.00104779 |
| ENSG00000100417 | PMM1 | | | ✓ | | | 0.001082236 |
| ENSG00000120742 | SERP1 | | | ✓ | | | 0.001091096 |
| ENSG00000169047 | IRS1 | | | ✓ | | | 0.001109005 |
| ENSG00000152763 | WDR78 | | | ✓ | | | 0.001214045 |
| ENSG00000122359 | ANXA11 | | | ✓ | | | 0.001235595 |
| ENSG00000197496 | SLC2A10 | | | ✓ | | | 0.001312637 |
| ENSG00000143641 | GALNT2 | | ✓ | ✓ | | | 0.001319818 |
| ENSG00000175215 | CTDSP2 | | | ✓ | | | 0.001337825 |
| ENSG00000130176 | CNN1 | | | ✓ | | | 0.001384892 |
| ENSG00000108946 | PRKAR1A | | | ✓ | | | 0.001386113 |
| ENSG00000113615 | SEC24A | | | ✓ | | | 0.00144094 |
| ENSG00000117475 | BLZF1 | | ✓ | ✓ | | | 0.001564808 |
| ENSG00000146112 | PPP1R18 | ✓ | | | | ✓ | 0.001639296 |
| ENSG00000175294 | CATSPER1 | | | ✓ | | | 0.001649295 |

| | | | | | | | |
|---|---|---|---|---|---|---|---|
| ENSG00000144566 | RAB5A | | | ✓ | | | 0.001665756 |
| ENSG00000160691 | SHC1 | | | ✓ | | | 0.001673495 |
| ENSG00000116299 | KIAA1324 | | | ✓ | | | 0.001686277 |
| ENSG00000251258 | RFPL4B | | | ✓ | | | 0.001753372 |
| ENSG00000110880 | CORO1C | | | ✓ | | | 0.001758843 |
| ENSG00000132561 | MATN2 | | | ✓ | | | 0.001783952 |
| ENSG00000087191 | PSMC5 | | | ✓ | | | 0.001796693 |
| ENSG00000236120 | RP11-733O18.1 | ✓ | | ✓ | | | 0.001904202 |
| ENSG00000180354 | MTURN | | | ✓ | | | 0.002046959 |
| ENSG00000177374 | HIC1 | | | ✓ | | | 0.002064441 |
| ENSG00000169750 | RAC3 | | | ✓ | | | 0.002106943 |
| ENSG00000102316 | MAGED2 | | | ✓ | | | 0.002143061 |
| ENSG00000223969 | AC002456.2 | ✓ | | ✓ | ✓ | ✓ | 0.002217917 |
| ENSG00000140575 | IQGAP1 | ✓ | | | ✓ | ✓ | 0.002328772 |
| ENSG00000156273 | BACH1 | | | ✓ | | | 0.00247067 |
| ENSG00000165152 | TMEM246 | | | ✓ | | | 0.002524239 |
| ENSG00000121879 | PIK3CA | | | ✓ | | | 0.002558307 |
| ENSG00000170458 | CD14 | | | ✓ | | | 0.002869644 |
| ENSG00000159579 | RSPRY1 | | | ✓ | | | 0.002914138 |
| ENSG00000184640 | SEPT9 | | | ✓ | | | 0.002964689 |
| ENSG00000137076 | TLN1 | | | ✓ | | | 0.002966974 |
| ENSG00000166997 | CNPY4 | | | ✓ | | | 0.003082753 |
| ENSG00000166444 | ST5 | | | ✓ | | | 0.0032449 |
| ENSG00000185565 | LSAMP | | ✓ | ✓ | | | 0.003333828 |
| ENSG00000147324 | MFHAS1 | | | ✓ | | | 0.003500301 |
| ENSG00000148832 | PAOX | | | ✓ | | | 0.003534803 |
| ENSG00000177963 | RIC8A | | | ✓ | | | 0.003543626 |
| ENSG00000150093 | ITGB1 | | | ✓ | | | 0.003608133 |
| ENSG00000171451 | DSEL | | | ✓ | | | 0.003629009 |
| ENSG00000170017 | ALCAM | | | ✓ | | | 0.003630989 |
| ENSG00000151348 | EXT2 | ✓ | ✓ | | | | 0.003656737 |
| ENSG00000198663 | C6orf89 | | | ✓ | | | 0.003815177 |
| ENSG00000204767 | FAM196B | ✓ | | | ✓ | | 0.004001858 |
| ENSG00000105321 | CCDC9 | | ✓ | ✓ | | | 0.004182484 |
| ENSG00000063322 | MED29 | | | ✓ | | | 0.004182484 |
| ENSG00000182272 | B4GALNT4 | | | ✓ | | | 0.004223157 |
| ENSG00000107185 | RGP1 | | | ✓ | | | 0.004541061 |
| ENSG00000136908 | DPM2 | | | ✓ | | | 0.004642232 |
| ENSG00000185909 | KLHDC8B | | | ✓ | | | 0.004719766 |
| ENSG00000196923 | PDLIM7 | | ✓ | | | | 0.005006073 |
| ENSG00000173614 | NMNAT1 | | | ✓ | | | 0.00512232 |
| ENSG00000100284 | TOM1 | | | ✓ | | | 0.005149844 |
| ENSG00000062598 | ELMO2 | | | ✓ | | | 0.005385626 |
| ENSG00000167695 | FAM57A | | | ✓ | | | 0.005529141 |
| ENSG00000154930 | ACSS1 | | | ✓ | | | 0.005559371 |

| | | | | | | | |
|---|---|---|---|---|---|---|---|
| ENSG00000106436 | MYL10 | | ✓ | | | | 0.005662159 |
| ENSG00000114353 | GNAI2 | | | ✓ | | | 0.00599371 |
| ENSG00000120656 | TAF12 | | | ✓ | | | 0.006030139 |
| ENSG00000164946 | FREM1 | | | ✓ | | | 0.006128099 |
| ENSG00000106266 | SNX8 | | | ✓ | | | 0.00623375 |
| ENSG00000163602 | RYBP | | | ✓ | | | 0.006259737 |
| ENSG00000175592 | FOSL1 | | | ✓ | | | 0.006296682 |
| ENSG00000127952 | STYXL1 | | | ✓ | | | 0.006471829 |
| ENSG00000168505 | GBX2 | | | ✓ | | | 0.006743743 |
| ENSG00000176887 | SOX11 | | | ✓ | | | 0.006976582 |
| ENSG00000198961 | PJA2 | | | ✓ | | | 0.006999097 |
| ENSG00000163902 | RPN1 | | | ✓ | | | 0.007113106 |
| ENSG00000260032 | NORAD | | | ✓ | | | 0.007202985 |
| ENSG00000188215 | DCUN1D3 | | | ✓ | | | 0.007462641 |
| ENSG00000090097 | PCBP4 | | | ✓ | | | 0.007666655 |
| ENSG00000114354 | TFG | | | ✓ | | | 0.008054431 |
| ENSG00000131446 | MGAT1 | | | ✓ | | | 0.00832568 |
| ENSG00000187678 | SPRY4 | | | ✓ | | | 0.008383833 |
| ENSG00000172247 | C1QTNF4 | | | ✓ | | | 0.008829241 |
| ENSG00000086598 | TMED2 | | | ✓ | | | 0.008843753 |
| ENSG00000142453 | CARM1 | | | ✓ | | | 0.009092757 |
| ENSG00000160408 | ST6GALNAC6 | | | ✓ | | | 0.009483151 |
| ENSG00000117614 | SYF2 | | | ✓ | | | 0.009537173 |
| ENSG00000168734 | PKIG | | | ✓ | | | 0.009635278 |
| ENSG00000137275 | RIPK1 | | | ✓ | | | 0.009831331 |
| ENSG00000109079 | TNFAIP1 | | | ✓ | | | 0.009866802 |
| ENSG00000183010 | PYCR1 | | | ✓ | | | 0.010434686 |
| ENSG00000154258 | ABCA9 | | | ✓ | | | 0.010552994 |
| ENSG00000173818 | ENDOV | | | ✓ | | | 0.010573861 |
| ENSG00000186001 | LRCH3 | | | ✓ | | | 0.010695377 |
| ENSG00000116649 | SRM | | | ✓ | | | 0.011311095 |
| ENSG00000121653 | MAPK8IP1 | | | ✓ | | | 0.012706089 |
| ENSG00000068308 | OTUD5 | | | ✓ | | | 0.01301553 |
| ENSG00000172667 | ZMAT3 | | | ✓ | | | 0.013104319 |
| ENSG00000163346 | PBXIP1 | | | ✓ | | | 0.014454047 |
| ENSG00000185189 | NRBP2 | | | ✓ | | | 0.014706223 |
| ENSG00000110536 | PTPMT1 | | | ✓ | | | 0.01476827 |
| ENSG00000171067 | C11orf24 | | | ✓ | | | 0.014779532 |
| ENSG00000177548 | RABEP2 | | | ✓ | | | 0.015019596 |
| ENSG00000112855 | HARS2 | | | ✓ | | | 0.015034391 |
| ENSG00000115977 | AAK1 | | | ✓ | | | 0.015440781 |
| ENSG00000169994 | MYO7B | | ✓ | | | | 0.015474478 |
| ENSG00000143315 | PIGM | | | ✓ | | | 0.015546758 |
| ENSG00000062716 | VMP1 | | | ✓ | | | 0.015600211 |
| ENSG00000116667 | C1orf21 | | ✓ | | | | 0.015682143 |

| | | | | | | | |
|---|---|---|---|---|---|---|---|
| ENSG00000110108 | TMEM109 | | | ✓ | | | 0.016288457 |
| ENSG00000171552 | BCL2L1 | | | ✓ | | | 0.016291534 |
| ENSG00000132581 | SDF2 | | | ✓ | | | 0.016875375 |
| ENSG00000105404 | RABAC1 | ✓ | ✓ | ✓ | | | 0.01728954 |
| ENSG00000101310 | SEC23B | | | ✓ | | | 0.018132594 |
| ENSG00000068028 | RASSF1 | | | ✓ | | | 0.018495992 |
| ENSG00000254560 | BBOX1-AS1 | | ✓ | | | | 0.018961443 |
| ENSG00000145431 | PDGFC | | | ✓ | | | 0.019101208 |
| ENSG00000188580 | NKAIN2 | ✓ | | ✓ | | | 0.01999429 |
| ENSG00000237765 | FAM200B | | | ✓ | | | 0.020070513 |
| ENSG00000135241 | PNPLA8 | | | ✓ | | | 0.02022049 |
| ENSG00000175220 | ARHGAP1 | | | ✓ | | | 0.02088622 |
| ENSG00000119820 | YIPF4 | | | ✓ | | | 0.021222324 |
| ENSG00000167881 | SRP68 | | | ✓ | | | 0.021384855 |
| ENSG00000109458 | GAB1 | | | ✓ | | | 0.021545004 |
| ENSG00000172638 | EFEMP2 | | | ✓ | | | 0.021616569 |
| ENSG00000256268 | RP11-221N13.3 | ✓ | | ✓ | | ✓ | 0.0219 |
| ENSG00000113812 | ACTR8 | | | ✓ | | | 0.022354634 |
| ENSG00000148634 | HERC4 | | | ✓ | | | 0.023666178 |
| ENSG00000197530 | MIB2 | | | ✓ | | | 0.024081593 |
| ENSG00000002822 | MAD1L1 | | | ✓ | | | 0.024175467 |
| ENSG00000150967 | ABCB9 | | | ✓ | | | 0.024945179 |
| ENSG00000197021 | CXorf40B | | | ✓ | | | 0.025399477 |
| ENSG00000172500 | FIBP | | | ✓ | | | 0.025579948 |
| ENSG00000180806 | HOXC9 | | | ✓ | | | 0.025888455 |
| ENSG00000157259 | GATAD1 | | | ✓ | | | 0.025987068 |
| ENSG00000167614 | TTYH1 | | | ✓ | | | 0.026131729 |
| ENSG00000225407 | CTD-2384B11.2 | | ✓ | | | | 0.026791189 |
| ENSG00000107651 | SEC23IP | | | ✓ | | | 0.027276336 |
| ENSG00000123342 | MMP19 | | | ✓ | | | 0.027524544 |
| ENSG00000136731 | UGGT1 | | | ✓ | | | 0.028037464 |
| ENSG00000172260 | NEGR1 | ✓ | | | ✓ | ✓ | 0.028123779 |
| ENSG00000136827 | TOR1A | | | ✓ | | | 0.028512492 |
| ENSG00000108433 | GOSR2 | | | ✓ | | | 0.028966071 |
| ENSG00000259969 | RP11-999E24.3 | ✓ | ✓ | | ✓ | ✓ | 0.0291856 |
| ENSG00000260136 | CTD-2270L9.4 | | | ✓ | | | 0.029544776 |
| ENSG00000092295 | TGM1 | | | ✓ | | | 0.029560628 |
| ENSG00000134153 | EMC7 | | | ✓ | | | 0.029659604 |
| ENSG00000236056 | GAPDHP14 | | | ✓ | | | 0.029746972 |
| ENSG00000272622 | RP11-395N3.2 | | | ✓ | | | 0.030746753 |
| ENSG00000186973 | FAM183A | | | ✓ | | | 0.031901192 |
| ENSG00000111752 | PHC1 | | | ✓ | | | 0.031901192 |
| ENSG00000131238 | PPT1 | | | ✓ | | | 0.031901192 |
| ENSG00000116954 | RRAGC | | | ✓ | | | 0.033680099 |
| ENSG00000101850 | GPR143 | | | ✓ | | | 0.034850555 |

| | | | | | | | |
|---|---|---|---|---|---|---|---|
| ENSG00000264247 | LINC00909 | | | ✓ | | | 0.034915992 |
| ENSG00000162591 | MEGF6 | | | ✓ | | | 0.035113634 |
| ENSG00000165861 | ZFYVE1 | | ✓ | | | | 0.035487724 |
| ENSG00000102100 | SLC35A2 | ✓ | | ✓ | | | 0.035742973 |
| ENSG00000182197 | EXT1 | | ✓ | ✓ | | | 0.035754833 |
| ENSG00000196981 | WDR5B | | | ✓ | | | 0.0363894 |
| ENSG00000213523 | SRA1 | ✓ | | ✓ | | | 0.03644822 |
| ENSG00000253161 | LINC01605 | | ✓ | ✓ | | | 0.036617723 |
| ENSG00000272455 | RP4-758J18.13 | | | ✓ | | | 0.037373918 |
| ENSG00000198551 | ZNF627 | | | ✓ | | | 0.037459511 |
| ENSG00000015532 | XYLT2 | | | ✓ | | | 0.038602218 |
| ENSG00000259877 | RP11-46C24.7 | | | ✓ | | | 0.038854845 |
| ENSG00000122545 | SEPT7 | | ✓ | ✓ | | | 0.039299384 |
| ENSG00000139318 | DUSP6 | ✓ | ✓ | ✓ | | ✓ | 0.040590686 |
| ENSG00000131370 | SH3BP5 | | | ✓ | | | 0.040682372 |
| ENSG00000101294 | HM13 | | | ✓ | | | 0.04117118 |
| ENSG00000149548 | CCDC15 | | | ✓ | | | 0.041616292 |
| ENSG00000115652 | UXS1 | | | ✓ | | | 0.043476592 |
| ENSG00000260482 | CTD-2196E14.9 | | | ✓ | | | 0.044084238 |
| ENSG00000127554 | GFER | | | ✓ | | | 0.045240386 |
| ENSG00000106617 | PRKAG2 | | ✓ | | | | 0.045573978 |
| ENSG00000006451 | RALA | | | ✓ | | | 0.046042341 |
| ENSG00000167977 | KCTD5 | | | ✓ | | | 0.047220845 |
| ENSG00000101940 | WDR13 | ✓ | | ✓ | | | 0.047220845 |
| ENSG00000152422 | XRCC4 | | | ✓ | | | 0.048199375 |
| ENSG00000182934 | SRPRA | | | ✓ | | | 0.048821255 |
| ENSG00000145882 | PCYOX1L | | | ✓ | | | 0.049105938 |
| ENSG00000253746 | RP11-527N22.2 | | | ✓ | | | 0.049171405 |

MS = GGAA-μsat, NMS = non-μsat, p-adj = adjusted p-value ( < 0.05)

**Supplementary table S3. Dual indexing primers used in CUT&Tag experiments (index sequences in bold)**

| **CUT&Tag sample** | **Forward primer (P5)** | **Reverse primer (P7)** |
|---|---|---|
| A673 EWS/FLI rep1 | AATGATACGGCGACCACCGAGATCTACAC**AGAGTAGA**TCGTCGGCAGCGTCAGATGTGTAT | CAAGCAGAAGACGGCATACGAGAT**CTAGTACG**GTCTCGTGGGCTCGGAGATGTG |
| A673 EWS/FLI rep2 | AATGATACGGCGACCACCGAGATCTACAC**GTAAGGAG**TCGTCGGCAGCGTCAGATGTGTAT | CAAGCAGAAGACGGCATACGAGAT**CTAGTACG**GTCTCGTGGGCTCGGAGATGTG |
| A673 Rb-IgG rep1 | AATGATACGGCGACCACCGAGATCTACAC**AGAGTAGA**TCGTCGGCAGCGTCAGATGTGTAT | CAAGCAGAAGACGGCATACGAGAT**TCGCCTTA**GTCTCGTGGGCTCGGAGATGTG |
| A673 Rb-IgG rep2 | AATGATACGGCGACCACCGAGATCTACAC**GTAAGGAG**TCGTCGGCAGCGTCAGATGTGTAT | CAAGCAGAAGACGGCATACGAGAT**TCGCCTTA**GTCTCGTGGGCTCGGAGATGTG |
| TC71 EWS/FLI rep1 | AATGATACGGCGACCACCGAGATCTACAC**CTCTCTAT**TCGTCGGCAGCGTCAGATGTGTAT | CAAGCAGAAGACGGCATACGAGAT**CATGCCTA**GTCTCGTGGGCTCGGAGATGTG |
| TC71 EWS/FLI rep2 | AATGATACGGCGACCACCGAGATCTACAC**CTCTCTAT**TCGTCGGCAGCGTCAGATGTGTAT | CAAGCAGAAGACGGCATACGAGAT**CCTCTCTG**GTCTCGTGGGCTCGGAGATGTG |

| | | |
|---|---|---|
| TC71 Rb-IgG rep1 | AATGATACGGCGACCACCGAGATCTACAC**CTCTCTAT**TCGTCGGCAGCGTCAGATGTGTAT | CAAGCAGAAGACGGCATACGAGAT**AGGAGTCC**GTCTCGTGGGCTCGGAGATGTG |
| TC71 Rb-IgG rep2 | AATGATACGGCGACCACCGAGATCTACAC**CTCTCTAT**TCGTCGGCAGCGTCAGATGTGTAT | CAAGCAGAAGACGGCATACGAGAT**GTAGAGAG**GTCTCGTGGGCTCGGAGATGTG |
| SK-N-MC EWS/FLI rep1 | AATGATACGGCGACCACCGAGATCTACAC**TATCCTCT**TCGTCGGCAGCGTCAGATGTGTAT | CAAGCAGAAGACGGCATACGAGAT**CTAGTACG**GTCTCGTGGGCTCGGAGATGTG |
| SK-N-MC EWS/FLI rep2 | AATGATACGGCGACCACCGAGATCTACAC**TATCCTCT**TCGTCGGCAGCGTCAGATGTGTAT | CAAGCAGAAGACGGCATACGAGAT**GCTCAGGA**GTCTCGTGGGCTCGGAGATGTG |
| SK-N-MC Rb-IgG rep1 | AATGATACGGCGACCACCGAGATCTACAC**TATCCTCT**TCGTCGGCAGCGTCAGATGTGTAT | CAAGCAGAAGACGGCATACGAGAT**TCGCCTTA**GTCTCGTGGGCTCGGAGATGTG |
| SK-N-MC Rb-IgG rep2 | AATGATACGGCGACCACCGAGATCTACAC**TATCCTCT**TCGTCGGCAGCGTCAGATGTGTAT | CAAGCAGAAGACGGCATACGAGAT**TTCTGCCT**GTCTCGTGGGCTCGGAGATGTG |
| EWS-502 EWS/FLI rep1 | AATGATACGGCGACCACCGAGATCTACAC**CTCTCTAT**TCGTCGGCAGCGTCAGATGTGTAT | CAAGCAGAAGACGGCATACGAGAT**GCTCAGGA**GTCTCGTGGGCTCGGAGATGTG |

| | | |
|---|---|---|
| EWS-502 EWS/FLI rep2 | AATGATACGGCGACCACCGAG ATCTACAC**GTAAGGAG**TCGTC GGCAGCGTCAGATGTGTAT | CAAGCAGAAGACGGCATAC GAGAT**GCTCAGGA**GTCTCG TGGGCTCGGAGATGTG |
| EWS-502 Rb-IgG rep1 | AATGATACGGCGACCACCGAG ATCTACAC**CTCTCTAT**TCGTCG GCAGCGTCAGATGTGTAT | CAAGCAGAAGACGGCATAC GAGAT**TTCTGCCT**GTCTCGT GGGCTCGGAGATGTG |
| EWS-502 Rb-IgG rep2 | AATGATACGGCGACCACCGAG ATCTACAC**GTAAGGAG**TCGTC GGCAGCGTCAGATGTGTAT | CAAGCAGAAGACGGCATAC GAGAT**TTCTGCCT**GTCTCGT GGGCTCGGAGATGTG |
| TTC-466 EWS/ERG rep1 | AATGATACGGCGACCACCGAG ATCTACAC**AACATGAT**TCGTCG GCAGCGTCAGATGTGTAT | CAAGCAGAAGACGGCATAC GAGAT**TCCTCTAC**GTCTCGT GGGCTCGGAGATGTG |
| TTC-466 EWS/ERG rep2 | AATGATACGGCGACCACCGAG ATCTACAC**TGGAAATC**TCGTCG GCAGCGTCAGATGTGTAT | CAAGCAGAAGACGGCATAC GAGAT**TCCTCTAC**GTCTCGT GGGCTCGGAGATGTG |
| TTC-466 Rb-IgG rep1 | AATGATACGGCGACCACCGAG ATCTACAC**AACATGAT**TCGTCG GCAGCGTCAGATGTGTAT | CAAGCAGAAGACGGCATAC GAGAT**TGCCTCTT**GTCTCGT GGGCTCGGAGATGTG |
| TTC-466 Rb-IgG rep2 | AATGATACGGCGACCACCGAG ATCTACAC**TGGAAATC**TCGTCG GCAGCGTCAGATGTGTAT | CAAGCAGAAGACGGCATAC GAGAT**TGCCTCTT**GTCTCGT GGGCTCGGAGATGTG |

| | | |
|---|---|---|
| A673_EF-Endo_CTCF_rep1 | AATGATACGGCGACCACCGAGATCTACAC**TAGATCGC**TCGTCGGCAGCGTCAGATGTGTAT | CAAGCAGAAGACGGCATACGAGAT**TCGCCTTA**GTCTCGTGGGCTCGGAGATGTG |
| A673_EF-Endo_CTCF_rep2 | AATGATACGGCGACCACCGAGATCTACAC**AGAGTAGA**TCGTCGGCAGCGTCAGATGTGTAT | CAAGCAGAAGACGGCATACGAGAT**CATGCCTA**GTCTCGTGGGCTCGGAGATGTG |
| A673_EF-Endo_Rb-IgG_rep1 | AATGATACGGCGACCACCGAGATCTACAC**TAGATCGC**TCGTCGGCAGCGTCAGATGTGTAT | CAAGCAGAAGACGGCATACGAGAT**CTAGTACG**GTCTCGTGGGCTCGGAGATGTG |
| A673_EF-Endo_Rb-IgG_rep2 | AATGATACGGCGACCACCGAGATCTACAC**AGAGTAGA**TCGTCGGCAGCGTCAGATGTGTAT | CAAGCAGAAGACGGCATACGAGAT**AGGAGTCC**GTCTCGTGGGCTCGGAGATGTG |
| A673_EF-KD_CTCF_rep1 | AATGATACGGCGACCACCGAGATCTACAC**CTCTCTAT**TCGTCGGCAGCGTCAGATGTGTAT | CAAGCAGAAGACGGCATACGAGAT**TCGCCTTA**GTCTCGTGGGCTCGGAGATGTG |
| A673_EF-KD_CTCF_rep2 | AATGATACGGCGACCACCGAGATCTACAC**GTAAGGAG**TCGTCGGCAGCGTCAGATGTGTAT | CAAGCAGAAGACGGCATACGAGAT**CATGCCTA**GTCTCGTGGGCTCGGAGATGTG |
| A673_EF-KD_Rb-IgG_rep1 | AATGATACGGCGACCACCGAGATCTACAC**TATCCTCT**TCGTCGGCAGCGTCAGATGTGTAT | CAAGCAGAAGACGGCATACGAGAT**CTAGTACG**GTCTCGTGGGCTCGGAGATGTG |

| | | |
|---|---|---|
| A673_EF-KD_Rb-IgG_rep2 | AATGATACGGCGACCACCGAGATCTACAC**GTAAGGAG**TCGTCGGCAGCGTCAGATGTGTAT | CAAGCAGAAGACGGCATACGAGAT**AGGAGTCC**GTCTCGTGGGCTCGGAGATGTG |
| A673_EF-Rescue_CTCF_rep1 | AATGATACGGCGACCACCGAGATCTACAC**TATCCTCT**TCGTCGGCAGCGTCAGATGTGTAT | CAAGCAGAAGACGGCATACGAGAT**TCGCCTTA**GTCTCGTGGGCTCGGAGATGTG |
| A673_EF-Rescue_CTCF_rep2 | AATGATACGGCGACCACCGAGATCTACAC**ACTGCATA**TCGTCGGCAGCGTCAGATGTGTAT | CAAGCAGAAGACGGCATACGAGAT**CATGCCTA**GTCTCGTGGGCTCGGAGATGTG |
| A673_EF-Rescue_Rb-IgG_rep1 | AATGATACGGCGACCACCGAGATCTACAC**CTCTCTAT**TCGTCGGCAGCGTCAGATGTGTAT | CAAGCAGAAGACGGCATACGAGAT**CTAGTACG**GTCTCGTGGGCTCGGAGATGTG |
| A673_EF-Rescue_Rb-IgG_rep2 | AATGATACGGCGACCACCGAGATCTACAC**ACTGCATA**TCGTCGGCAGCGTCAGATGTGTAT | CAAGCAGAAGACGGCATACGAGAT**AGGAGTCC**GTCTCGTGGGCTCGGAGATGTG |
| TC71_EF-Endo_CTCF_rep1 | AATGATACGGCGACCACCGAGATCTACAC**TGGAAATC**TCGTCGGCAGCGTCAGATGTGTAT | CAAGCAGAAGACGGCATACGAGAT**TTCTGCCT**GTCTCGTGGGCTCGGAGATGTG |
| TC71_EF-Endo_CTCF_rep2 | AATGATACGGCGACCACCGAGATCTACAC**TGGAAATC**TCGTCGGCAGCGTCAGATGTGTAT | CAAGCAGAAGACGGCATACGAGAT**CCTCTCTG**GTCTCGTGGGCTCGGAGATGTG |

| | | |
|---|---|---|
| TC71_EF-Endo_Rb-IgG_rep1 | AATGATACGGCGACCACCGAGATCTACAC**TGGAAATC**TCGTCGGCAGCGTCAGATGTGTAT | CAAGCAGAAGACGGCATACGAGAT**TCGCCTTA**GTCTCGTGGGCTCGGAGATGTG |
| TC71_EF-Endo_Rb-IgG_rep2 | AATGATACGGCGACCACCGAGATCTACAC**TGGAAATC**TCGTCGGCAGCGTCAGATGTGTAT | CAAGCAGAAGACGGCATACGAGAT**CATGCCTA**GTCTCGTGGGCTCGGAGATGTG |
| TC71_EF-KD_CTCF_rep1 | AATGATACGGCGACCACCGAGATCTACAC**AACATGAT**TCGTCGGCAGCGTCAGATGTGTAT | CAAGCAGAAGACGGCATACGAGAT**TTCTGCCT**GTCTCGTGGGCTCGGAGATGTG |
| TC71_EF-KD_CTCF_rep2 | AATGATACGGCGACCACCGAGATCTACAC**AACATGAT**TCGTCGGCAGCGTCAGATGTGTAT | CAAGCAGAAGACGGCATACGAGAT**CCTCTCTG**GTCTCGTGGGCTCGGAGATGTG |
| TC71_EF-KD_Rb-IgG_rep1 | AATGATACGGCGACCACCGAGATCTACAC**AACATGAT**TCGTCGGCAGCGTCAGATGTGTAT | CAAGCAGAAGACGGCATACGAGAT**TCGCCTTA**GTCTCGTGGGCTCGGAGATGTG |
| TC71_EF-KD_Rb-IgG_rep2 | AATGATACGGCGACCACCGAGATCTACAC**AACATGAT**TCGTCGGCAGCGTCAGATGTGTAT | CAAGCAGAAGACGGCATACGAGAT**CATGCCTA**GTCTCGTGGGCTCGGAGATGTG |

**Supplementary table S4. Dual index sequences used in Hi-C libraries**

| ***In situ* Hi-C sample** | **P5 index** | **P7 index** |
|---|---|---|
| A673_EF-Endo_Replicate1 | AGGCGAAG | CGCTCATT |
| A673_EF-Endo_Replicate2 | AGGCGAAG | GAGATTCC |
| A673_EF-KD_Replicate1 | TAATCTTA | ATTACTCG |
| A673_EF-KD_Replicate2 | TAATCTTA | TCCGGAGA |
| A673_EF-Rescue_Replicate1 | TAATCTTA | CGCTCATT |
| A673_ EF-Rescue _Replicate2 | TAATCTTA | GAGATTCC |

**Supplementary table S5. Primers used in 2nd PCR amplification of 4C libraries**

| 4C sample | Universal forward primer | Reverse primer (index in bold) |
|---|---|---|
| A673 EF-Endo rep1 | AATGATACGGCGACCACCGAGATCTACACTCTTTCCCTA | CAAGCAGAAGACGGCATACGAGAT**CGTGAT**GTGACTGGAGT |
| A673 EF-KD rep1 | AATGATACGGCGACCACCGAGATCTACACTCTTTCCCTA | CAAGCAGAAGACGGCATACGAGAT**CACTGT**GTGACTGGAGT |
| A673 EF-Rescue rep1 | AATGATACGGCGACCACCGAGATCTACACTCTTTCCCTA | CAAGCAGAAGACGGCATACGAGAT**CTGATC**GTGACTGGAGT |
| A673 EF-Endo rep2 | AATGATACGGCGACCACCGAGATCTACACTCTTTCCCTA | CAAGCAGAAGACGGCATACGAGATTG**TTGACT**GTGACTGGA |
| A673 EF-KD rep2 | AATGATACGGCGACCACCGAGATCTACACTCTTTCCCTA | CAAGCAGAAGACGGCATACGAGATGT**GCGGAC**GTGACTGG |
| A673 EF-Rescue rep2 | AATGATACGGCGACCACCGAGATCTACACTCTTTCCCTA | CAAGCAGAAGACGGCATACGAGATTA**CGTACG**GTGACTGGA |
| TC71 EF-Endo rep1 | AATGATACGGCGACCACCGAGATCTACACTCTTTCCCTA | CAAGCAGAAGACGGCATACGAGAT**CGTGAT**GTGACTGGAGT |
| TC71 EF-Endo rep1 | AATGATACGGCGACCACCGAGATCTACACTCTTTCCCTA | CAAGCAGAAGACGGCATACGAGAT**CACTGT**GTGACTGGAGT |
| TC71 EF-KD rep2 | AATGATACGGCGACCACCGAGATCTACACTCTTTCCCTA | CAAGCAGAAGACGGCATACGAGAT**CTGATC**GTGACTGGAGT |
| TC71 EF-KD rep2 | AATGATACGGCGACCACCGAGATCTACACTCTTTCCCTA | CAAGCAGAAGACGGCATACGAGATTG**TTGACT**GTGACTGGA |
